# Markers of biosynthetic innovation in the Cyanobacteria

**DOI:** 10.64898/2026.09.06.749771

**Authors:** Rachel Mizzi, Brett Anthony Neilan, Leanne Andrea Pearson, Verlaine Joy Timms

## Abstract

Cyanobacteria are a rich source of specialised metabolites of ecological, biomedical and industrial significance, yet discovery of novel compounds is constrained by rediscovery and uneven taxonomic exploration. Diversification of specialised metabolism constitutes a key evolutionary innovation associated with ecological expansion. Further, understanding how biosynthetic capacity is distributed across cyanobacterial lineages and environments is critical for guiding natural product discovery. We therefore analysed 939 cyanobacterial genomes spanning 11 orders and five habitat types to identify taxonomic and ecological hotspots of biosynthetic potential. We evaluated several indicators of biosynthetic innovation, including singleton BGCs and specialized enzymes and assessed the distribution of cyanobacterial toxin BGCs. Conserved terpene and ribosomally synthesised and post-translationally modified peptide (RiPP) clusters dominated, accounting for >50% of identified BGCs. These two classes exhibited high sequence conservation, averaging 16.41 and 28.10 similarity links per BGC, respectively. Conversely, NRPS and polyketide synthase clusters were less common (18.5% and 2.5% of total BGCs, respectively) but displayed considerable sequence diversity, with a mean of 1.9 and 0.79 links per BGC, respectively. Enzymes associated with potentially novel chemistries were rare, with halogenases and cytochrome P450s present in 4.9% and 1.27% of BGCs, respectively, whereas amidinotransferases were detected in only 0.42% of BGCs. Symbiotic, terrestrial and thermal spring cyanobacteria, particularly Nostocales, showed elevated biosynthetic potential, whereas marine Synechococcales genomes showed comparatively low biosynthetic potential. Putative toxin BGCs were enriched in freshwater Nostocales, Chroococcales and Oscillatoriales. These findings link phylogeny, ecology and genomic features to prioritise taxa, habitats and biosynthetic pathways for future natural product discovery.

## Introduction

Cyanobacteria exist in diverse aquatic and terrestrial habitats, ranging from thermal springs to polar deserts (Walter, et al. 2017; Willis and Woodhouse 2020). They were the first organisms to perform oxygenic photosynthesis (Cardona 2019) and their lengthy evolutionary history and broad dispersion have led to significant morphological and metabolic diversity within the phylum. It is hypothesized that the ecological success of cyanobacteria is underpinned by their capacity to synthesize specialized metabolites – small bioactive compounds, such as non-ribosomal peptides, polyketides, alkaloids, siderophores and mycosporine-like amino acids (Dittmann, et al. 2015; Belknap, et al. 2020). Some of these metabolites (e.g., microcystin and saxitoxin) are highly toxic and are an ecological and public health risk (Codd, et al. 2005; O’Neil, et al. 2012; Hackett, et al. 2013). Others have potentially beneficial applications as active ingredients in pharmaceuticals, cosmetics and agrochemicals (Balskus and Walsh 2010; Katoch, et al. 2016). Furthermore, certain species of cyanobacteria are capable of biosorption or degradation of harmful environmental contaminants, making them useful for bioremediation in wastewater or other contaminated sites (Deviram, et al. 2020). However, in several instances, the biosynthetic pathways associated with these compounds are poorly understood. The ecological, public health, and industrial relevance of cyanobacteria underscores the importance of investigating their diverse metabolic potential, which may offer valuable insights for environmental management, medical research and biotechnology.

The phylum has undergone significant taxonomic revision in the face of gene and, more recently, genome sequencing and these revisions are ongoing ^16^. The literature currently recognises 11 orders, including Chroococcales, Chroococcidiopsidales, Gloeobacterales, Gloeomargaritales, Nostocales, Oscillatoriales, Pleurocapsales, Pseudanabaenales, Spirulinales, Synechococcales and Thermostichales, with several orders displaying polyphyly (e.g., Synechococcales and Oscillatoriales) (Willis and Woodhouse 2020; Strunecký, et al. 2023). Certain taxa have been associated with the production of specialized metabolites, however, research has disproportionately focused on toxic bloom-forming freshwater genera such as *Microcystis*, *Planktothrix*, and *Dolichospermum* due to their direct impact on water quality and public health.

Predicting metabolic capabilities in cyanobacteria remains challenging, although certain patterns emerge in relation to both habitat and phylogeny. For example, the cyanotoxin microcystin is typically associated with freshwater taxa, whereas lyngbyatoxin is typically produced by marine species (Pearson, et al. 2016; Ding, et al. 2018). Nonetheless, toxicity within these aquatic environments during blooms can be sporadic. Phylogenetic relationships offer a stronger framework for predicting biosynthetic potential, however, this approach is limited by confounding factors such as horizontal gene transfer, gene deletion events and convergent evolution, all of which can obscure true evolutionary and functional relationships (Patel 2016; Chevrette, et al. 2020). Understanding the distribution of these pathways may assist in understanding their evolution, identifying novel pathways for their production or tracing occurrence of outbreaks.

A significantly more robust predictor of specialized metabolite production is the presence of biosynthetic gene clusters (BGCs) (Blin, et al. 2021). However, only a small percentage of cyanobacterial BGCs have been characterised to date (Robert, et al. 2010; Dittmann, et al. 2015; Kultschar 2018). The elucidation of BGCs responsible for major cyanotoxins (microcystin, nodularin, cylindrospermopsin, saxitoxin, anatoxin, lyngbyatoxin and guanitoxin), has enabled the development of rapid molecular detection methods, particularly qPCR-based assays (e.g., Phytoxigene™ CyanoDTec (Romanis, et al. 2021; Pinheiro, et al. 2025)), for monitoring harmful cyanobacterial populations. These advances have significantly improved the management of water quality and seafood safety. Collectively, they highlight the utility and importance of a BGC-centric approach in cyanobacterial natural product discovery and environmental monitoring.

Furthermore, rediscovery of metabolites is a frequent challenge within the field of secondary metabolite research (Krug and Müller 2014). The prioritization of strains with novel BGCs, and therefore, potentially novel chemistries, can minimize compound rediscovery (a common problem with traditional bioassay-guided discovery pipelines). Therefore, increasing scientific understanding of BGC diversity in a larger dataset of cyanobacteria may assist in reducing wasted efforts on potentially non-novel clusters. Furthermore, comparisons between characterised databases of BGCs and newly sequenced genomes can assist in identifying new variants or indicate that a pathway might be novel when it does not cluster with known BGCs (known as singletons). Additionally, genome-based approaches can provide access to specialized metabolites that are poorly expressed (or not expressed) under laboratory culture conditions (Yu, et al. 2024). This can be achieved through strain engineering or the heterologous expression of BGCs in industrial hosts, such as *E. coli* (Ongley, et al. 2013; Soeriyadi, et al. 2021). While the technology is rapidly advancing, several limitations remain. For example, genomic studies frequently rely on reference databases or arbitrary cut-off thresholds for matching genes and BGCs to previously characterised counterparts. This approach can be problematic when analyzing diverse microbial phyla such as the Cyanobacteria, where species or niche-specific variations may hinder the detection of homologous genes or clusters, leading to underestimation of biosynthetic potential.

The aim of this study was to identify environmental sources and cyanobacterial taxa with high biosynthetic potential, particularly those potentially capable of producing novel, toxic or underexplored specialized metabolites. To investigate this, we systematically mined 939 cyanobacterial genomes using AntiSMASH (Blin, et al. 2021) in combination with the BigSCAPE pipeline (Navarro-Munoz, et al. 2020) to annotate and classify BGCs. Furthermore, Hmmer (Finn, et al. 2011; Potter, et al. 2018) was used to identify candidate enzymes associated with novel biosynthetic pathways. Characterized BGCs of interest from the MIBiG database were also used as references to find these pathways within our data. Statistical analyses were employed to assess the relationships between BGC diversity and ecological origin. Our results provide a comprehensive picture of the biosynthetic landscape across ecological and phylogenetic subgroups of cyanobacteria and yield important findings that can inform future bioprospecting efforts, as well as environmental monitoring and management strategies.

## Methods

### Dataset curation and quality assessment

A dataset of 860 cyanobacterial genomes previously curated by Timms, et al., 2023 (Timms, et al. 2023) was the foundation for this study. This core dataset was supplemented with 56 recently sequenced genomes from the Commonwealth Scientific and Industrial Research Organisation (CSIRO) Australian National Algae Culture Collection and 86 additional closed genomes from GenBank. Public genome data was downloaded and curated from GenBank on the 15^th^ of March 2023 using filters for closed *Cyanobacteriota* genomes so that additions to the previous dataset would be high quality. Assemblies were assessed with QUAST (version 5.0.2) (Gurevich, et al. 2013). As a quality control measure, assemblies exceeding 400 contigs were excluded from the final dataset. Genomes passing this filter were retained for BGC analysis, annotation, and phylogenetic profiling.

### Phylogenomic tree reconstruction

Genome annotations were performed using Prokka (version 1.14.5) (Seemann 2014). For phylogenomic inference, we used PhyloPhlAn (version 3.0.60) (Asnicar, et al. 2020) which builds species trees based on a concatenated alignment of 400 universal proteins conserved across bacterial genomes. The aligned protein sequences were used as input for IQ-TREE (version 1.6.7) (Nguyen, et al. 2015). The ModelFinder option was used in IQ-Tree to determine the most appropriate model for the data (LG+R10) and the resulting maximum likelihood tree was visualised and annotated using Interactive Tree of Life (iTOL) (Letunic and Bork 2019).

### Genome mining

The standalone version of AntiSMASH (version 6.6.0) (Blin, et al. 2021) was used to identify BGCs in the cyanobacterial genomes using the default parameters. The resulting AntiSMASH output files were processed using BiG-SCAPE (version 1.1.5) (Navarro-Munoz, et al. 2020) for evolutionary context and visualization of gene cluster architecture. CORe Analysis of Syntenic Orthologs to prioritize Natural Product Biosynthetic Gene Clusters (CORASON), implemented within the BiG-SCAPE pipeline, was used to visualize gene clusters (Navarro-Munoz, et al. 2020). Within the network analysis, a 0.3 distance threshold (loose) was used to determine gene cluster families. Any BGCs that did not have any connections to other BGCs that met this threshold were considered singletons.

### Statistical analysis

To evaluate whether BGC abundance varied significantly between genomes derived from different environmental sources or taxonomic orders, two-way analysis of variance (ANOVA) was conducted in R (version 4.4.0). Homoscedasticity was assessed by viewing residuals vs fitting plot Q-Q residuals lots. The two independent variables were environmental origin and order. The dependent variable was total BGC count per genome, as identified by AntiSMASH. Tukey’s Honestly Significant Difference (Tukey’s HSD) post-hoc test was used for pairwise comparisons. Statistical significance was defined as *P* <0.05.

### Genome mining for target biosynthetic enzymes

To investigate the distribution and abundance Tryptophan halogenases PF04820 and Acyl halogenase CylC-like A0A0A1VQW2 and amidinotransferases SxtG Aphanizomenongracile UAM529 (A0A1M4BLL8) within cyanobacterial BGCs, reference protein sequences corresponding to the target enzyme classes were retrieved from the InterPro or Hmmer database and used as queries. Protein sequences were extracted from AntiSMASH-generated .gbk output files using GenBank_to (version 0.42, available https://github.com/linsalrob/genbank_to) and queried using hmmscan (part of the HMMER suite) (Finn, et al. 2011; Potter, et al. 2018). An E-value threshold of 0.001, identity ≥70%, query coverage ≥85% and was applied to retain only statistically significant matches.

For identification of cytochrome P450 monooxygenases, we incorporated a curated set of 341 cyanobacterial protein sequences from (Khumalo, et al. 2020). These sequences were used as queries in phmmer against the custom BGC database. More stringent filtering criteria were applied to retain only high-confidence hits (E-value ≥1×10^−5^, identity ≥90%, query coverage ≥85%) given that these had previously been found in cyanobacterial genomes.

### Identification of putative cyanotoxin BGCs

To identify gene cluster families (GCFs) that contained a known toxin BGC, BiG-SCAPE was used with the database flag. This incorporated the Minimum Information about a Biosynthetic Gene (MIBiG) (Zdouc, et al. 2025) database into the analysis. Within the MIBiG database, cyanotoxin clusters were identified (Table 1). BGCs that were grouped into the same GCF as a cyanotoxin within the bigscape network were designated as putative toxin BGCs.

**Table 1.**
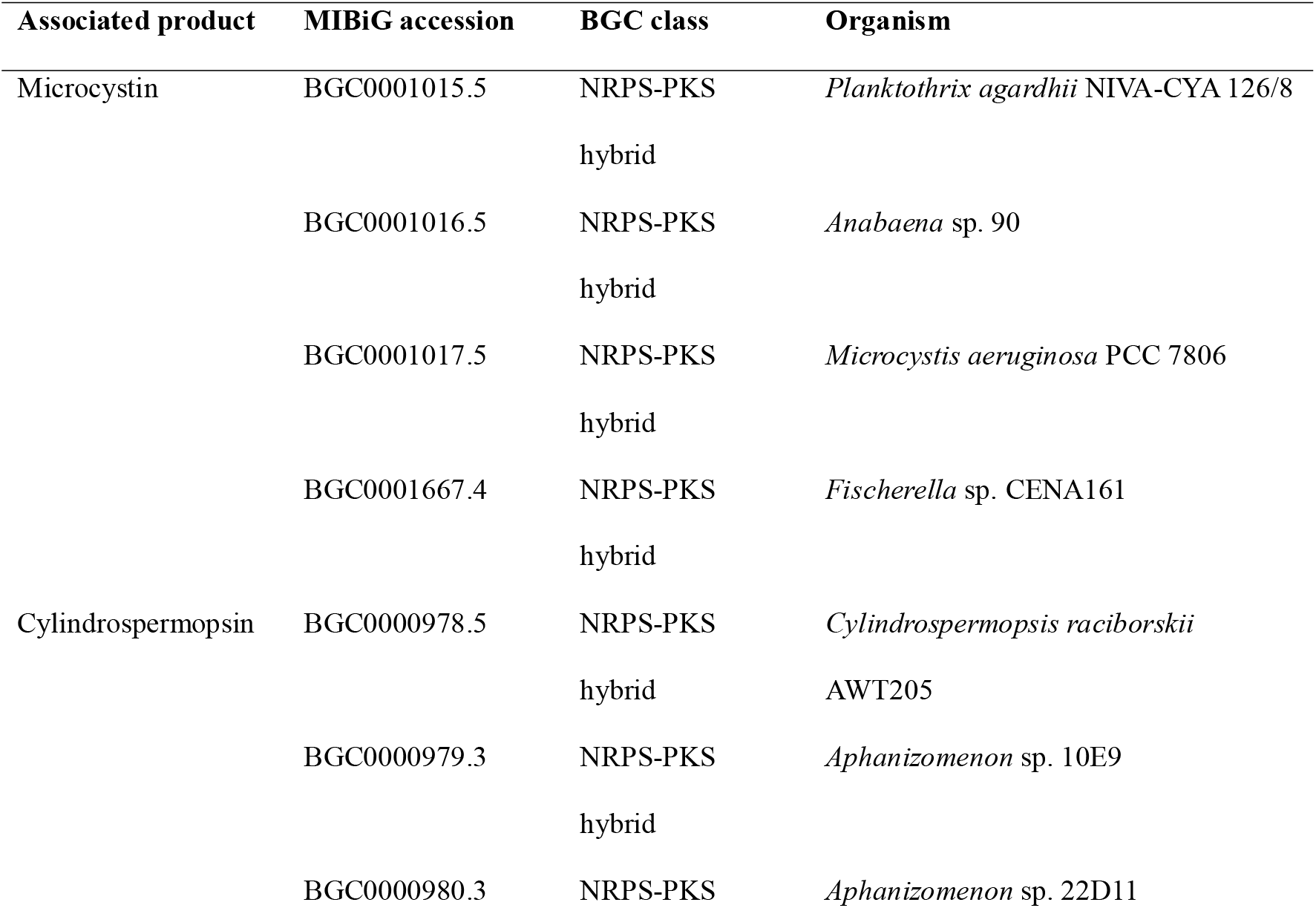
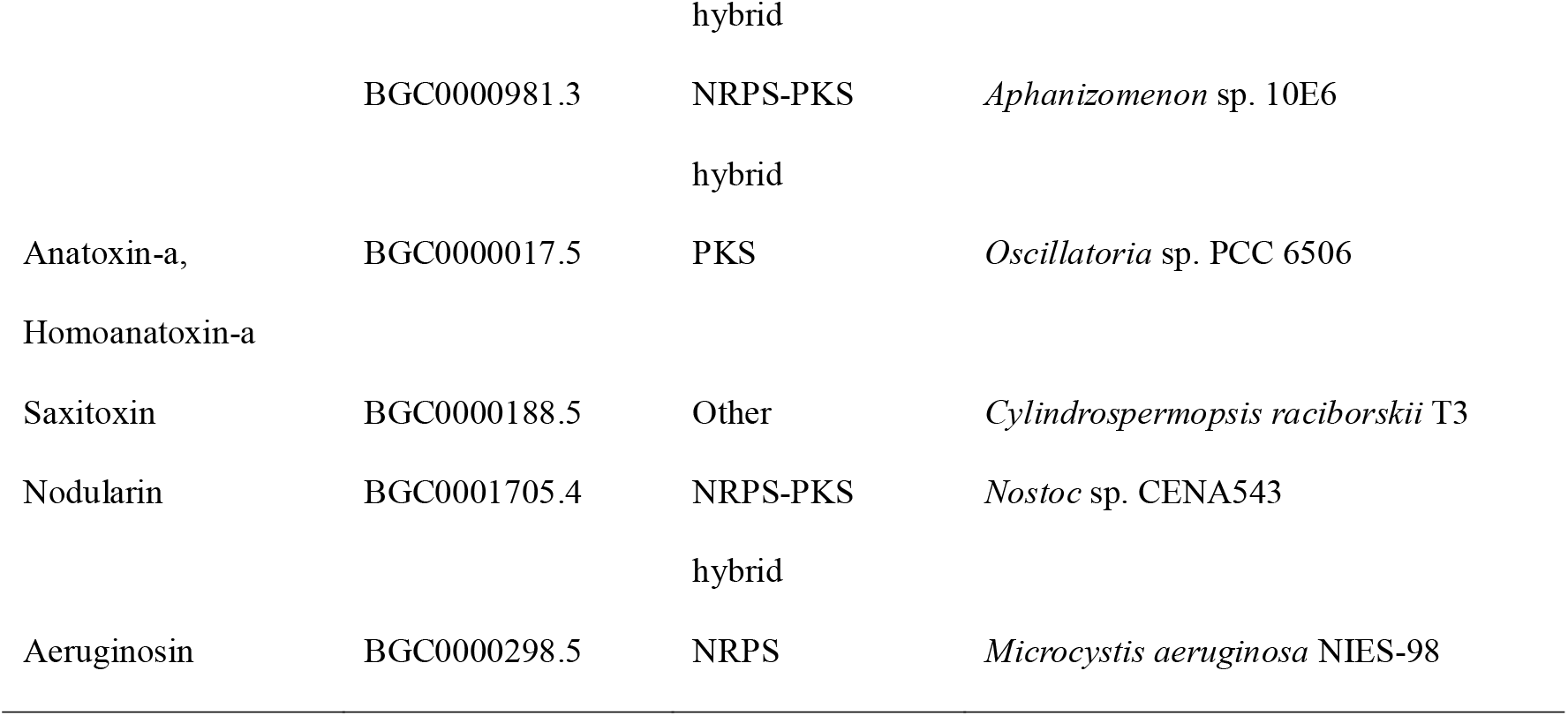
Cyanotoxin BGCs from the MIBiG repository retrieved 04/08/2025.

## Results and discussion

### Study objectives and impact

The aim of this study was to identify markers of BGC innovation in Cyanobacteria and environmental sources and cyanobacterial lineages with high biosynthetic potential. Specifically, we were interested in those capable of producing harmful, novel or underexplored specialised metabolites, such as cyanotoxins, guanidinylated and halogenated compounds. Analysis of 939 genomes spanning 11 orders and five habitat categories, several trends emerged in the distribution, abundance and diversity of biosynthetic gene clusters (BGCs) and markers of novelty such as rare enzymes and singleton BGCs.

### Dataset curation and general statistics

The curated dataset contained 939 genomes. Nostocales (n = 326, 35%) dominated the dataset, followed by Synechococcales (257, 27%), Oscillatoriales (125, 13%), Chroococcales (108, 11%) and Pseudanabaenales (93, 10%). Several orders, Chroococcidiopsidales (8, <1%), Gloeobacterales (4, <1%), Gloeomargaritales (2, < 1%), Spirulinales (6, < 1%), Pleurocapsales, (8, < 1%) and Thermostichales (2, < 1%), were sparsely represented. (Figure 1a). Habitat representation was similarly uneven. Freshwater (n = 235, 25%) and marine isolates (227, 26%) comprised half the dataset, while thermal spring isolates (47, 5%), symbiotic (46, 5%) and terrestrial isolates (127, 14%) made up a smaller but ecologically important proportion. A substantial fraction of genomes (257, 27%) were from sources that were not specified (Figure 1b). This uneven representation across habitats and lineages may reflect the historical emphasis on freshwater toxin producers and marine primary producers. Despite this uneven representation, the dataset provides a comprehensive overview of cyanobacterial diversity, capturing both metabolically streamlined taxa and lineages known for extensive specialised metabolism.

**Figure 1.**
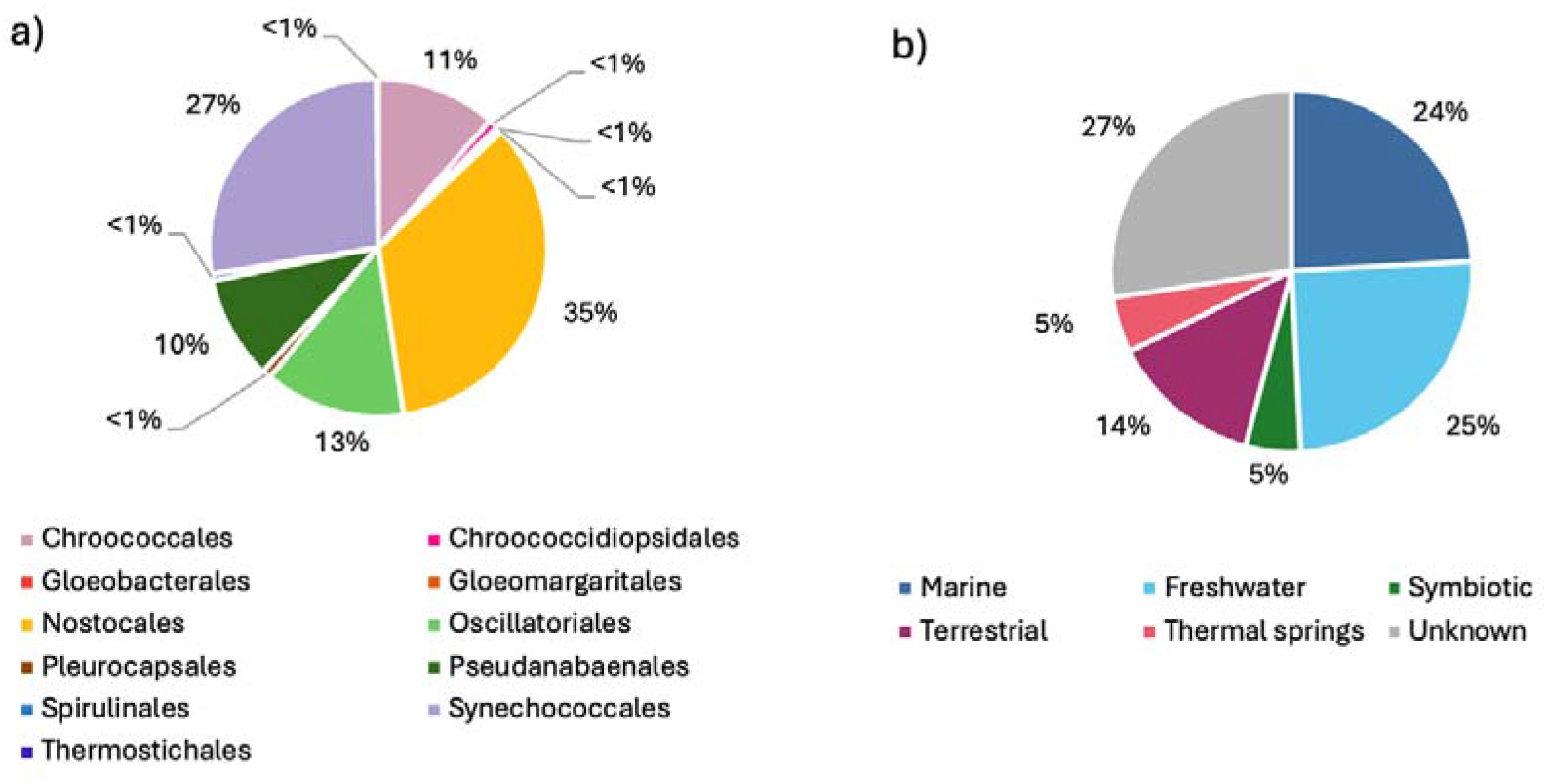
Representation of cyanobacterial orders a) and habitats b) in the curated dataset of 939 cyanobacterial genomes.

### Phylogenomic Summary

Phylogenomic analysis for majority of this dataset was previously report by Timms, et al., 2023 (Timms, et al. 2023). Hence, we do not go into detail here. The phylogeny demonstrated the previously described monophyletic clades of Nostocales and Chroococcidiopsidales and Spirulinales. Chroococcales, Gloeobacterales and Gloeomargaritales also formed their distinct phylogenetic clusters, though a small number of outliers were noted (Figure 2). In contrast, Oscillatoriales, Synechococcales, Pleurocapsales and Pseudanabaenales displayed polyphyletic distributions consistent with previous work, indicating the suitability of this phylogenic framework for subsequent analyses of BGC distribution (Willis and Woodhouse 2020; Chen, et al. 2021; Strunecký, et al. 2023).

**Figure 2.**
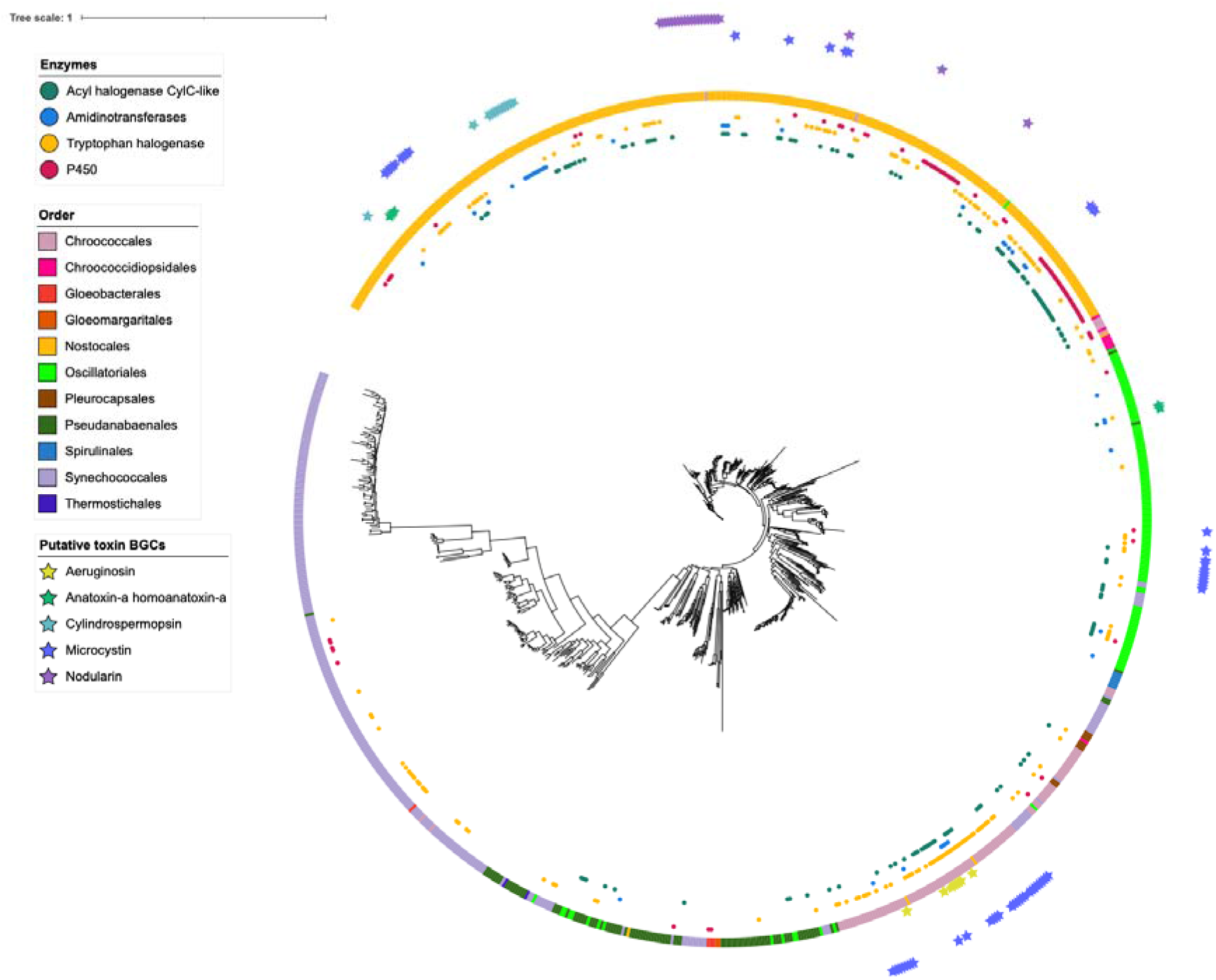
Maximum likelihood phylogenomic tree inferred from 400 concatenated universal proteins, representing 939 cyanobacterial genomes. The ring indicates the cyanobacterial order assigned to each genome. Coloured dots indicate the presence of specific biosynthetic enzymes (halogenases, amidinotransferases or cytochrome P450 monooxygenases) encoded within predicted BGCs. Coloured stars indicate putative toxin BGCs.

#### BGC abundances

Previous studies estimate that cyanobacterial genomes typically include an average of five BGCs, representing approximately 5-6% percent of the genome, with some strains carrying up to 23 BGCs in extreme cases (Calteau, et al. 2014; Ferrinho, et al. 2024). In line with previous work, within the 939 genomes examined, we identified 9,617 BGCs. The dataset exhibited a mean of 10.24 and a median of nine BGCs per genome.

Nostocales demonstrated the highest BGC abundance and had significantly (P < 0.05) more BGCs than all other orders (mean□=□16.97, range□2–41), consistent with their role as prolific secondary metabolite producers (Calteau, et al. 2014; Ferrinho, et al. 2024). High BGC densities were also observed in Pleurocapsales (mean□=□14), Chroococcales (mean□=□10.44) and Chroococcidiopsidales (mean□=□10.25), exceeding those of Oscillatoriales (mean□=□8.72) and contrasting with earlier reports, potentially due to improved taxonomic sampling (Calteau, et al. 2014) (Figure 3). In contrast, Synechococcales had significantly fewer BGCs than all other orders (P < 0.05)(mean□=□3.74; 257 genomes) exhibiting reduced biosynthetic capacity, trends that were also demonstrated previously (Calteau, et al. 2014; Cameron, et al. 2024; Ferrinho, et al. 2024).

**Figure 3.**
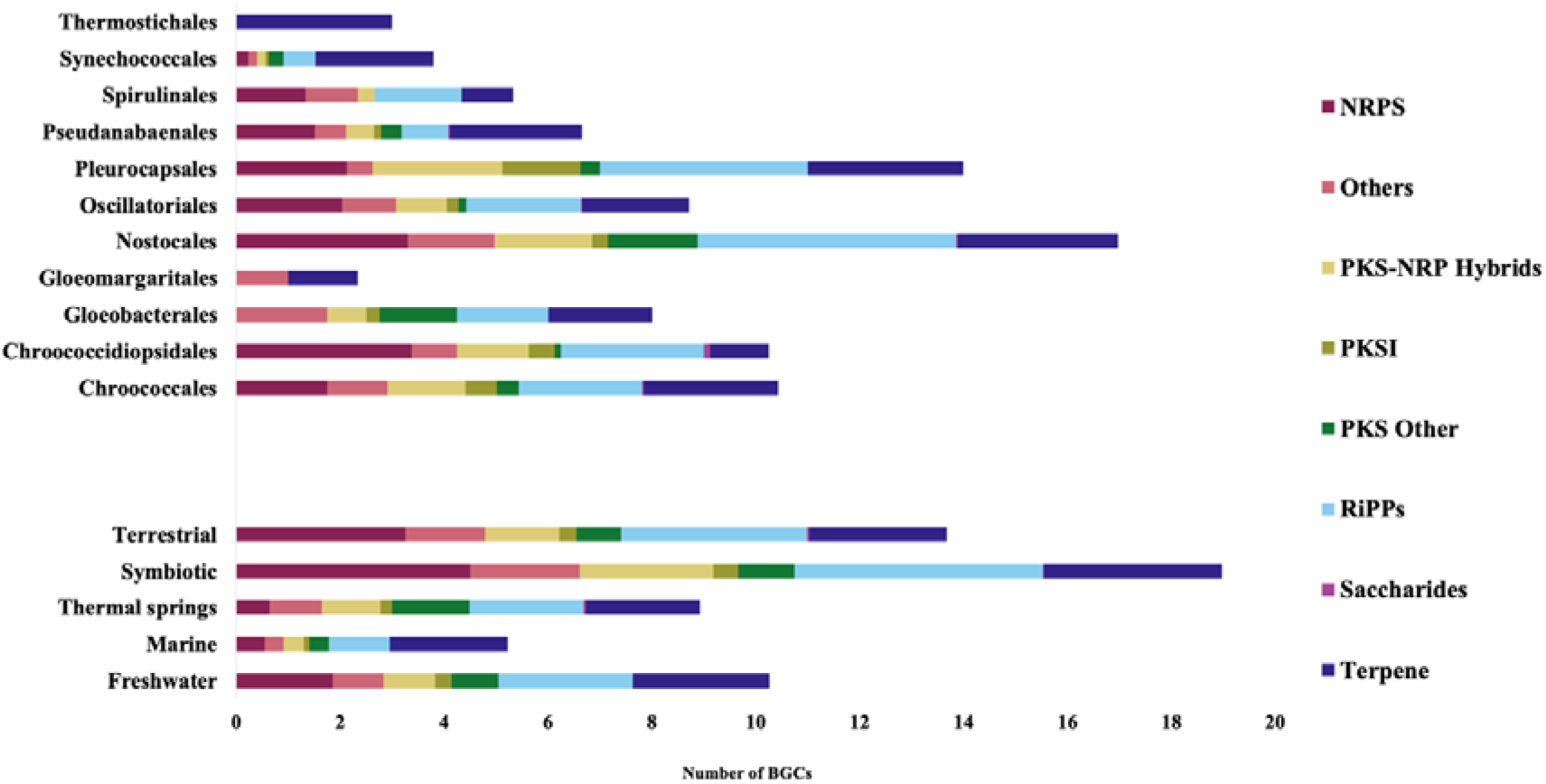
Distribution of BGC classes across cyanobacterial orders and hahbitats. Bar plots show the average number of BGCs per genome, grouped by (a) taxonomic order and (b) habitat. Bars are subdivided by BGC class, as predicted by antiSMASH, with colours corresponding to classes indicated in the legend. Data represent 9,617 putative BGCs identified from 939 cyanobacterial genomes.

BGC abundance also varied significantly between symbiotic, freshwater, terrestrial and thermal-spring genomes (P□<□0.001). Symbiotic cyanobacteria showed the highest BGC densities, consistent with the metabolic demands of host-associated lifestyles and coevolutionary pressures. This contrasted with previous work which demonstrated streamlining of obligate cyanobacterial genomes (Cameron, et al. 2024). Unfortunately, we were unable to distinguish between obligate and facultative symbionts and the differences in findings may be reflective of this. Also akin to previous work (Cameron, et al. 2024), marine taxa exhibited significantly reduced BGC abundance, indicative of genome streamlining in oligotrophic environments, while terrestrial and thermal-spring cyanobacteria possessed richer and more variable BGC complements (Figure 3).

#### Diversity of BGC classes

BGC class distribution among cyanobacterial orders has been studied extensively elsewhere (Popin, et al. 2021; Cameron, et al. 2024; Ferrinho, et al. 2024), therefore we focused on within-class diversity and markers of potential novelty. Additional breakdown by order and source are detailed in the Supplementary Tables 1-6 and Supplementary Figures 1-8.

Sequence similarity network analysis revealed substantial differences in diversity within BGC classes, measured by the proportion of singletons (clusters which contain no links to any others) and average connectivity (links per BGC). Links represent a similarity in sequence, gene content, domain architecture or predicted products (Navarro-Munoz, et al. 2020).

##### Ribosomally synthesized and post-translationally modified peptides (RiPPs)

RiPP clusters were the most abundant classes comprising approximately one quarter of the total BGC complement (2,473 clusters, 26% of all BGCs) and were widely distributed across the phylum. This finding reenforces their role as fundamental chemical mediators in cyanobacterial ecology, where they are known to contribute to defence, signalling, and niche establishment. These clusters form peptide natural products generated from short precursor peptides that are enzymatically tailored to produce structurally diverse and often highly bioactive compounds (Cao, et al. 2021). Their structural diversity and potent bioactivities, exemplified by lanthipeptides, cyanobactins, and microviridins, also make RiPPs valuable candidates for pharmaceutical and industrial applications, with several subclasses already serving as scaffolds for antimicrobial and enzyme-modulating compounds.

Predicted products (n = 53) were dominated by lanthipeptide-class-v biochemistries (28.6% of RiPPs), followed by RRE-containing (11.8%), cyanobactin (10.4%), lanthipeptide-class-ii (8%), lassopeptide (7.3%), microviridin (6.4%) and RiPP-like (5.7%) classes. An additional 46 unique RiPPs product types such as ranthipeptides and spliceotides were observed at low frequencies (1 to 100 occurrences, Supplementary Table 4). Despite their extensive distribution, these clusters were highly conserved (28.1% singletons, 3.06 links) (Figure 4), reflecting functional conservation across cyanobacteria, as in indicated across other bacteria (Li, et al. 2024). Future work should look at these less frequently observed RiPPs products to see if they exist in the same genomes as other RiPPs as accessory genome or they are the sole RiPPs within genomes as a functional replacement to more conversed clusters.

**Figure 4.**
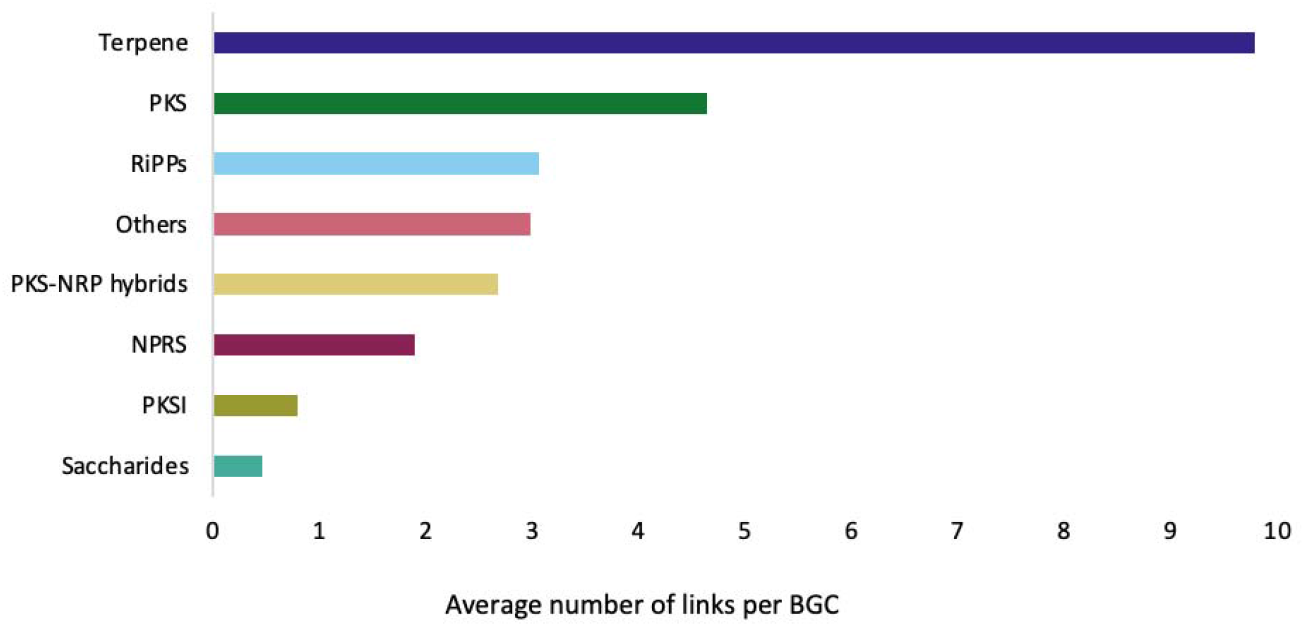
Sequence similarity network connectivity across BGC classes. Bar plot illustrating the mean connectivity within BiG-SCAPE-generated networks sequence similarity network for major BGC classes, based on 9,617 predicted BGCs from 939 cyanobacterial genomes. Connectivity was measured as the average node degree (number of links per cluster), reflecting similarity in gene content, domain architecture, or predicted products.

##### Terpenes

Terpenes are a large and functionally diverse class of isoprenoid compounds that play essential roles in photosynthesis, membrane stability, stress tolerance, and chemical signalling. Beyond their ecological roles, terpenoids are also of substantial applied value, forming the basis of many pharmaceuticals, fragrances, food additives and natural biocontrol agents. Their chemical versatility and tuneable structural scaffolds additionally position them as promising substrates for green chemistry, sustainable materials and industrial biotechnology (Mund, et al. 2022). Similar to RiPP clusters, terpene clusters exhibited a similar broad distribution and high frequency (2,420 clusters, 25%). Occurring in 934 of the 939 genomes analysed. With an average of 2.6 and range of zero to six clusters per genome, they ranked second the most frequently observed BGC classes.

Terpene clusters were the most conserved BGC type (16.41% singletons, average 9.79 links per cluster) (Figure 4). Despite the chemical diversity of terpenoids (key compounds in photosynthesis, stress responses, and defence) their shared isoprenoid backbone likely constrains biosynthetic pathway diversity (Pattanaik and Lindberg 2015).

Five Oscillatoriales genomes, primarily from terrestrial environments, lacked simple terpene clusters but instead contained terpene-NRPS-like hybrids. Such hybridization may represent an evolutionary strategy for generating novel terpenoid products, as seen in some extremophilic cyanobacteria (Machado, et al. 2025). Given the metabolic diversity and industrial potential of terpenoids (Pattanaik and Lindberg 2015), these rare hybrid clusters warrant further investigation as targets for bioprospecting. Considering the high degree of conservation of this BGC class, laboratory investigations should focus on these for cloning investigations.

##### Nonribosomal peptide synthetase (NRPS)

Nonribosomal peptides constitute one of the most structurally diverse and bioactive classes of natural products, with many cyanobacterial compounds, such as microcystins, nodularins, aeruginosins and cryptophycins, playing key ecological roles in competition, defence and signalling, while also serving as important scaffolds for anticancer, antimicrobial and enzyme-modulating drug development (Süssmuth and Mainz 2017).

The third most common class (1,771 clusters, 18%) was also the third most diverse, with 41.14% singletons and averaging 1.9 links per BGC (Figure 4). Their previous correlation with larger genome sizes (Wang, et al. 2014) suggesting they belong to the accessory rather than core genome. Furthermore, these BGCs were most prevalent in the Nostocales and symbiotic and terrestrial environments where comparatively larger genome sizes are prevalent (Calteau, et al. 2014). Together, their high diversity and association with the accessory genome supports their role in evolutionary innovation.

##### Type I polyketide synthase (TIPKS)

Polyketides produced by cyanobacteria represent a chemically diverse and often highly toxic class of natural products with complex modular architectures, yielding metabolites such as curacins, jamaicamides and scytophycins that mediate defence, predation resistance and ecological competition. They have also been recognised for their anticancer, antiparasitic and neuroactive properties (François, et al. 2025).

TIPKS clusters made up only 2% of all BGCs found (n = 237) and their markedly different prevalence by order and environment places them as likely candidates for biosynthetic innovation to new environments. TIPKS clusters were also highly diverse (50.63% singletons, 0.79 links) (Figure 4), further indicating their role niche adaptation for Cyanobacteria in diverse habitats.

##### Type III polyketide synthase (TIIIPKS)

TIIIPKS (n =186) was the only cluster type enriched in marine cyanobacteria and the streamlined order Synechococcales (25.3%, mean = 0.2 per genome) compared to the typical enriched orders such as Chroococcales (17%, mean = 0.17), Oscillatoriales (3.1%, mean = 0.03) and Nostocales (2.5%, mean = 0.02). TIIIPKS clusters were absent from Gloeomargaritales, Spirulinales, and Thermostichales, indicating possible lineage specificity. As these clusters were not a separate category within BiG-SCAPE, identification of singletons was not completed.

In Synechococcales, nearly all TIIIPKS clusters were simple, with only one terpene-hybrid exception. The scarcity of hybrid TIIIPKS clusters in this order suggests retention of the ancestral TIIIPKS forms, while hybrids, common in Nostocales and Oscillatoriales, likely evolved later, consistent with their designation as younger lineages (Komarek 2016; Willis and Woodhouse 2020). This trend aligns with the structural simplicity and broad substrate specificity of TIIIPKS enzymes compared to other PKS types (Katsuyama and Ohnishi 2012). The functional relevance of the simple TIIIPKS clusters in marine environments verses the hybrid clusters in later branches is an interesting evolutionary question and further investigation here may provide insights on the transition of ancient Cyanobacteria from aquatic to terrestrial habitats.

##### PKS-NRPS hybrids

Hybrid clusters (n = 1,028, 11%) are typically larger and more complex than those that can be categorised as a single class. Singletons made up roughly a third of BGCs (33.97%, average links 2.67). This is likely reflective of their roles as larger, multi-domain enzymes compared to other PKS clusters.

Within this cluster type, the most common predicated product appears to be biased towards simpler architectures, with TIPKS.NRPS combinations dominant (50.2%). A minority (2.6%) of these clusters were observed in four or fewer genomes. This pattern suggests that while hybridization between PKS and NRPS pathways is widespread in cyanobacteria, evolutionary pressures may favour more streamlined biosynthetic systems. Alternatively, these more complex structures may be the result of extensive horizontal gene transfer and continual adaptation to changing environments, or limitation of using draft genomes in BGC analyses.

Within Synechococcales, hybrid clusters were almost exclusively found in polyphyletic clades, distinct from the main marine-dominant lineage. This is a further indication that these clusters are the result of niche adaptation in less stable non-aquatic environments. Expansion of these BGCs in non-marine habitats even within the Synechococcales is further evidence that hybridization of pathways may be how Cyanobacteria create novel biosynthetic products as they transition to new environments. Comparative genomes of Synechococcales within the main branch and the polyphyletic lineages could demonstrate how these ancient life forms transitioned to new environments and should be a future direction of Cyanobacterial studies.

##### Polyketide synthase (PKS) others

PKS other clusters were the most commonly called PKS category (746 clusters, 8%). In contrast to the highly diverse TIPKS, PKS “other” subtypes were more conserved (19.79% singletons, 4.64 links). Although previous studies have acknowledged PKS diversity, most have focussed on individual subtypes (Jenke-Kodama and Dittmann 2009). These findings suggest functional and evolutionary divergence among PKS subtypes, particularly when comparing TIPKS to others.

PKS other clusters were the only category that was more dominant in thermal spring derived genomes than other habitats (64%, mean = 1.49). Previous research into thermal spring cyanobacteria demonstrated that these species had smaller genomes than their non-thermal counterparts but were enriched in some gene categories (Alcorta, et al. 2020). Results here suggest that cyanobacteria from this environment may have unique PKS clusters that confer a survival advantage here and other non-essential clusters have been reduced.

##### Saccharides

Saccharide-derived natural products are specialised metabolites built from carbohydrate precursors, often functioning in cell-cell recognition, stress protection and antimicrobial defence (McCranie and Bachmann 2014). Further, in some microbes such as *Streptomyces* they form the basis of clinically important aminoglycoside antibiotics (Yu, et al. 2017). Saccharide clusters were the rarest class in the dataset with only 15 identified (<1%), consistent with previous metabolomic and genomic studies (Li, et al. 2019; He, et al. 2025) indicating that glycosylated specialised metabolites may be uncommon in cyanobacteria. However, this may be due to AntiSMASH being optimised for more modular pathways with conserved domains and saccharide secondary metabolism can overlap with primary metabolic pathways.

Singletons were present in this category (60%, 0.47 average links) but due to difficulties in identifying secondary metabolite clusters that produce saccharides, it is unlikely that these products are rare or truly novel.

##### Other clusters

The “Other” category encompasses a broad range of cluster types that did not fall into the aforementioned classes (921 clusters, 10% of total). Despite encompassing a high number of predicted products (n = 135) and widespread abundance this category was more conserved than NRPs, PKSI and hybrid categories (29.86% singletons, 2.99 links). This unexpected trend suggests that even diverse subclasses and predicted products may share core biosynthetic frameworks.

The most common products (36% of total) in this class included ladderane (n = 101), NRPS.betalactone (n = 63), arylpolyene (n = 59), resorcinol (n = 57) and indole (n = 56). Contrastingly, 265 predicted products were only observed 30 times or less within the dataset. This indicates that these poorly characterised BGCs are an untapped trove of novelty. However, any rare clusters would require long read sequencing before characterisation.

#### Distribution of target biosynthesis enzymes across taxa and habitats

Recent advances in genome sequencing and bioinformatics have accelerated the identification of cyanobacterial BGCs, the functional annotation of their encoded enzymes, and the prediction of the structures and bioactivities of their associated metabolites. Within BGCs, certain genes may be particularly valuable “biosynthetic hooks” for metabolites that have useful or rare biochemistries (e.g., halogen and guanidine groups) and are difficult to produce synthetically (Crowe, et al. 2021).

Identifying such enzymes within large genomic datasets is a critical first step toward understanding the distribution of these compounds and determining which taxa may possess the capacity for their biosynthesis. To this end, we conducted a targeted survey of genes putatively encoding halogenases, cytochrome P450 (CYP450) and amidinotransferases within BGCs.

##### Halogenases

Halogenation occurs throughout nature, contributes to natural product diversity (Agarwal, et al. 2017) and can improve properties such as lipophilicity and metabolic stability. In drug design, for example, fluorination of 3-piperidinylindole antipsychotics improves serotonin receptor binding and bioavailability (Benedetto Tiz, et al. 2022), while halogenation can enhance bioadhesive performance via hydrophobic and electronic effects (Yang, et al. 2025).

In the examined cyanobacteria, BGC-associated halogenases were common and broadly distributed across taxa, consistent with the presence of previously described CylC-like enzymes (Eusebio, et al. 2021). Tryptophan halogenases (PF04820 were detected in 326 BGCs across 171 genomes, occurring predominantly in Nostocales (n = 145, 2.6% of BGCs from this order) and Chroococcales (n = 124, 10.9%, with smaller representation in Synechococcales (n = 28, 2.9%), Oscillatoriales (n = 17, 1.6%), Chroococcidiopsidales (n = 9, 11%), and Pseudanabaenales (n = 2, 0.3%).

Across environments, BGC-associated halogenases occurred in freshwater (n = 109, 4.5% 0f BGCs in this source), host-associated (n = 29, 3.3%), marine (n = 29, 2.4%), terrestrial (n = 53, 3%), and thermal spring (n = 5, 1.2%) cyanobacteria and those of unknown origin (n = 100, 3.4%). These enzymes were associated primarily with NRPS (n = 39, 2.2%), “others” (n = 32, 3.4%), PKS-NRP hybrid (n = 56, 5.3%), TIPKS (n = 105, 43.9%), PKS other (n = 30, 4%), RiPPs (38, 1.5%) and terpene (n = 24, 1%) clusters. Genomes from several sources and lineages encoded more than one halogenase, and several clusters encoded multiple halogenases, suggesting functional redundancy or pathway modularity (Figure 2).

BGC-associated acyl halogenase CylC-like proteins (A0A0A1VQW2) were detected 146 times in 132 genomes, primarily in PKS-NRP hybrid clusters (n = 54, 37% of detections), T1PKS (n = 43, 29.5%), “others” (n = 35, 24%), PKS others (n = 10, 6.8%) and NRPS (n =4, 2.7%) clusters.

Most detections occurered in Nostocales (n = 90, prevalence = 27.6% of genomes), followed by Chroococcales (n = 25, 23.1%), Oscillatoriales (15, 12%), Chroococcidiopsidales and Pleurocapsales (n = 2, 25% for both orders) and Pseudanabaenales and Synechococcales (n = 6, 6.5% and 2.3% respectively).

Environmental distribution included freshwater (37, prevalence = 15.7% of genomes), terrestrial (37, 29.1%), unknown source (33, 12.8%), thermal springs (20, 42.6%), symbiotic (13, 28.3%) and marine (6, 2.6%) (Figure 2). Notably, Nostocales from thermal springs, showed disproportionately high frequency of CylC-like halogenases, whereas Synechococcales from the same environments contained few, suggesting lineage-specific ecological adaptation.

##### Amidinotransferases

Amidinotransferases are enzymes that catalyse the transfer of an amidino group from one donor molecular (usually L-arginine) to an acceptor molecule containing a primary amino group. These enzymes are involved in the biosynthesis of guanidino-containing compounds, including several cyanobacterial toxins (Muenchhoff, et al. 2010; Barón Sola, et al. 2013; Hackett, et al. 2013). In this dataset, they were rarely observed within BGCs and were not specific to any particular taxonomic order, habitat or cluster type. 44 matches to SxtG (A0A1M4BLL8) from the saxitoxin pathway were found across 39 genomes and 40 different clusters. Five genomes contained two clusters encoding amidinotransferases, and in four of these clusters, two amidinotransferases co-occurred.

Taxonomically, BGC-associated amidinotransferases were most frequent in Nostocales (68.2% of total enzyme matches, 9.2% of genomes in order), Oscillatoriales (16%, 5.5%), Chroococcales (13.6%, 5.5%), and Pseudanabaenales (2.3%, 1.1%). Genomes encoding these enzymes did not form distinct phylogenetic clusters, except for a small group within Nostocales. Environmentally, these clusters were found in terrestrial (22.7% of total matches, 7.9% genomes from this source), freshwater (36.4%, 6.8%), symbiotic (6.8%, 6.5%), unknown (31.8%, 5.4%), and marine (2.3%, 0.4%).

BiGSCAPE predicted BGC products included simple NRPS (NRPS n = 6, NRPS-like n = 2), ‘other’ BGCs (one cyanobactin-NRPS and one phosphonate-lanthipeptide-class-v.hglE-KS.PUFA and betalactone-TIPKS-NRPS-NRPS-like n = 2 of each and six hglE-KS PUFA), PKS others (hglE-KS n = 1), PKS-NRP hybrids (TIPKS-NRPS n = 2, TIPKS-NRPS-like n = 16 and TIPKS-NRPS-NRPS-like n = 3) and terpene (n = 2).

Further interrogation of genomes harbouring amidinotransferases within BGCs is warranted to clarify their role in toxin biosynthesis.

##### Cytochrome P450 monooxidases

Cytochrome P450 monooxidase (CYP450) enzymes catalyse the oxidation of inert C-H bonds and play essential roles in the biosynthesis of carotenoids and other light-harvesting pigments. These enzymes may represent an evolutionary link between plants and cyanobacteria ^38^ and have significant industrial and pharmaceutical applications, including in the production of metabolites for bioremediation and drug delivery (Notonier, et al. 2016).

Previous studies have shown that cyanobacteria have fewer CYP450 enzymes compared to *Bacillus*, *Streptomyces* and *Mycobacterium* species but exhibit greater diversity, with only a minority encoded within BGCs (Khumalo, et al. 2020). Using the same query sequences as prior work but restricted to BGCs, our analysis found enzymes predominantly associated with NRPS and terpene clusters. Of the 341 unique BGC-associated CYP450 sequences identified, 41 different sequences were detected in 80 of 939 cyanobacterial genomes, resulting in 122 BGC-associated matches (Figure 2).

These enzymes were most frequent in Nostocales (82.8% total matches, 31% genomes in order), with fewer occurrences in Synechococcales and Oscillatoriales (4.9% matches each, 2.3%, 4.8% respectively), Chroococcales and Gloeobacterales (3.3% each, 3.7% and 100%, respectively), and Chroococcidiopsidales (0.8%, 12.5%).

Environmentally, BGC-associated CYP450s were widely distributed, with the highest frequency in genomes from thermal springs (40.2% of matches, 100% of genomes). CYP450 enzymes from these environments have been shown to exhibit enhanced thermal stability (Kundral, et al. 2025), highlighting their potential for industrial exploitation. Other sources, including symbiotic (7.4%, 19.6%), terrestrial (10.7%, 10.2%), freshwater (15.6%, 8.1%), unknown (16.4%, 7.8%), and marine (9.8%, 5.3%) environments, had lower representation.

These CYP450 enzymes were associated with 22 different predicted products across 7 BGC classes. The BGC class breakdowns were terpenes (39.3% of identified matches) and PKS-NRPS hybrids (24.6%), followed by NRPS (13.1%), PKS-other (9.8%), RiPPs (8.2%), others (4.1%), and a single T1PKS cluster (0.8%), as detailed in Supplementary Table 8.

Two Nostocales genomes, GCF_000316625.1 (freshwater) and GCF_000317535.1 (terrestrial), each encoded four CYP450 enzymes. In GCF000316625.1, the CYP450s were located in a ladderane-associated “others” cluster, a lassopeptide-producing RiPP cluster, and a hybrid TIPKS-NRPS cluster. In GCF000317535.1, CYP450s occurred in a hybrid TIPKS-NRPS cluster, a terpene cluster and an “others” cluster. The latter was predicted by BiG-SLiCE to participate in the biosynthesis of a complex multinodular product incorporating elements of thiopeptides, terpenes, NRPSs, thioamitides, and TIPKS domains.

#### Distribution and frequency of putative toxin BGCs

Cyanotoxins represent a significant global public health concern (Mihali, et al. 2008) and elucidating the genetic basis for their biosynthesis has already enabled the development of rapid diagnostic tools to support environmental monitoring and management (Al-Tebrineh, et al. 2012; Woodhouse, et al. 2016). Nevertheless, continued investigation of toxin BGCs is essential, as sequence divergence and the emergence of novel variants may lead to false negatives in current genetic assays. Beyond their ecological and health impacts, cyanotoxins and other natural products constitute a rich source of structurally unique compounds with potential medical and industrial applications (Swain, et al. 2017). Therefore, understanding their distribution and diversity across cyanobacterial lineages and environments is essential for both risk assessment and biotechnological exploitation.

A total of 111 BGCs from 106 genomes within our dataset were grouped into gene cluster families (GCFs) containing a characterised toxin BGC. The majority were NRPS-PKS hybrids (n = 98, 88.3%), followed by NRPS (n = 8, 7.2%) and PKS (n = 5, 4.5%). Most putative toxins (n = 60, 54.1%) were associated with GCFs containing microcystin accessions from the MIBiG database. Additional associations included nodularin (n = 24, 22.5%), cylindrospermopsin (n = 13, 11.7%), aeruginosin (n = 8, 7.2%), anatoxin-a/homoanatoxin-a (n = 5, 4.5%) (Figure 5a).

**Figure 5.**
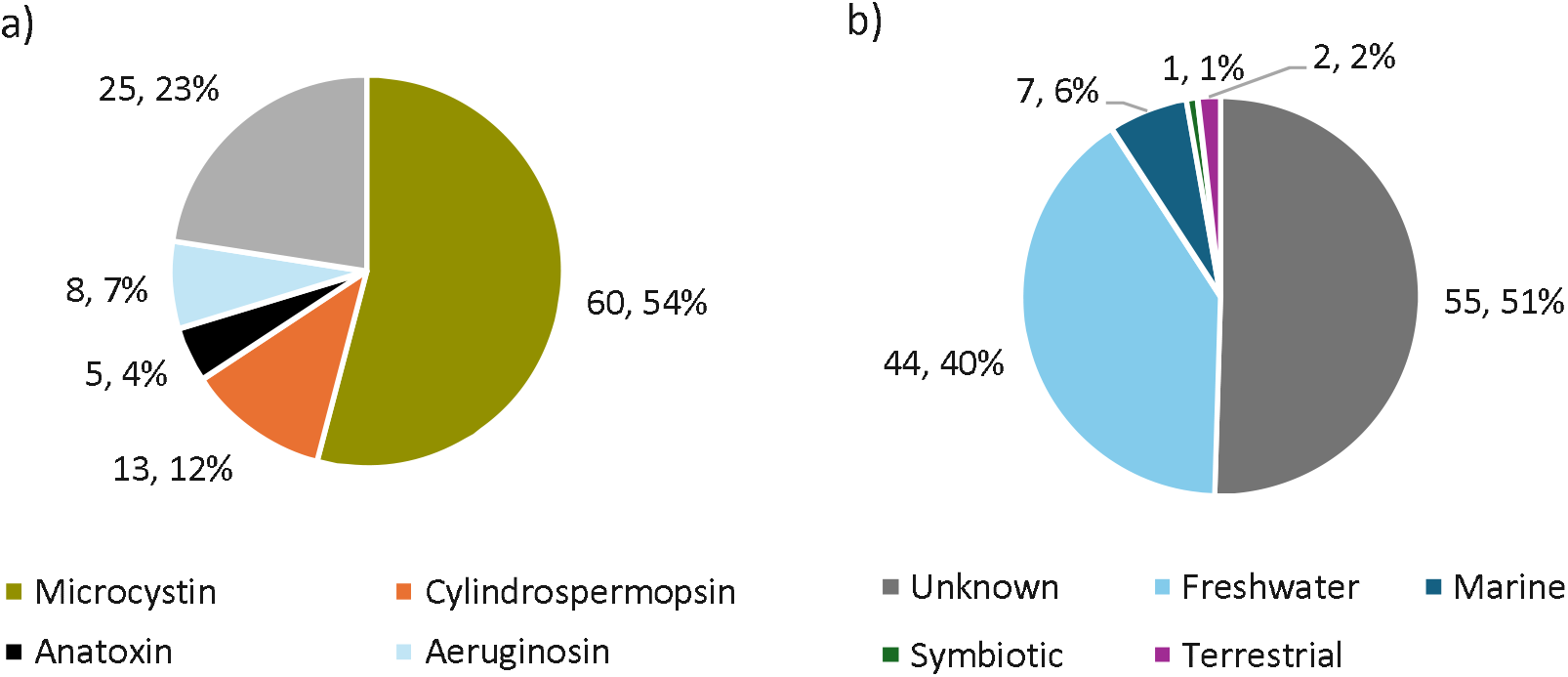
Distribution and frequency of putative cyanotoxin and other BGCs of interest in the dataset with respect to a) gene cluster families and b) genome source environments.

Our analysis revealed a widespread but uneven distribution across orders and environments for several of our BGCs of interest. Taxonomically, Nostocales genomes had the highest prevalence toxin BGCs (n = 61, 18.7%), followed by Chroococcales (n = 38, 35.2%) and Oscillatoriales (n = 12, 9.5%) (Figure 2). These orders have all been recognised as toxin producers (Bashir, et al. 2023). Several toxin BGCs were restricted to a single order, and some were further restricted to a small phylogenetic branch. For example, cylindrospermopsin were limited to a closely related group of freshwater Nostocales, while nodularin BGCs were predominantly found within a single lineage of *Nodularia spumigena*, with a few unspeciated Nostocales scattered elsewhere. Previous studies demonstrate that nodularin-producing species include *N. spumigena, Nodularia harveyana* and other *Nostoc* species (Nowruzi and Porzani 2021), consistent with our findings and supporting the validity of our identification method.

Putative aeruginosin BGCs were detected only in freshwater Chroococcales, although literature reports their presence in other orders (Ishida, et al. 2009). This discrepancy may reflect high BGC and gene sequence divergence between orders, preventing BiG-SCAPE from grouping them into the same gene cluster family. Alternately, the use of draft genome sequences may have resulted in incomplete genomic and therefore BGC data from genomes that contained these clusters, leading to false negatives in our dataset.

Microcystin associated BGCs were the most abundant ground in the dataset, and were identified across all known toxin-producing genera (*Anabaena, Oscillatoria, Nostoc, Phormidium* and *Fischerella*). The broad taxonomic distribution of these BGCs suggests an ancestral origin of microcystins relative to other cyanotoxins. Earlier studies have suggested that the nodularin synthetase evolved from a microcystin synthetase progenitorv(Moffitt and Neilan 2004; Rantala, et al. 2004) which is consistent with our observations.

With respect to habitat, genomes of unknown origin had the highest frequency of BGCs of interest, with 21.4% genomes (n = 55 BGCs) containing at least one cluster. This was followed by freshwater genomes (n = 44, 18.7%), marine genomes (n = 7, 3.1%) terrestrial genomes (n = 2, 4.3%) and symbiotic genomes (n = 1, 2.1%) (Figure 5b). Notably, freshwater genomes were the sole identified source for several toxin BGCs, including cylindrospermopsin, aeruginosin and anatoxin. This pattern aligns with the hypothesis that some of these metabolites may confer a competitive advantage within microbial communities such as deterring predation during bloom formation.

## Conclusions and future directions

By integrating BGC diversity, sequence similarity networks and rare biosynthetic enzymes, we identified cyanobacterial taxa and habitats that demonstrate biosynthetic innovation and represent promising targets for novel chemistry. The recovery of known toxin BGCs demonstrates the utility of this approach for contextualising potentially novel clusters within larger genomic datasets and minimising compound rediscovery.

Our analysis suggests that specialised metabolism in cyanobacteria is partitioned into conserved and adaptive components. Widespread and highly conserved BGC classes such as terpenes and RiPPs likely support broad ecological functions, whereas the more diverse and low-connectivity of the NRPS and PKSI classes may reflect adaptation to specific ecological niches. The distribution of halogenases and cytochrome P450 enzymes further refines this prioritisation, highlighting lineages and habitats enriched in unusual biochemical capabilities, particularly the Nostocales and symbiotic and terrestrial cyanobacteria.

The diversity of cyanobacterial metabolism from non-aquatic habitats demonstrates the importance of targeting underexplored environments and non-model taxa to maximise discovery outcomes. This study identifies high-priority cyanobacterial taxa, habitats and biosynthetic pathways for future genome-guided natural product discovery and bioprospecting. Further interrogation of metagenomics, long-read sequencing and functional characterisation will be critical for capturing uncultured diversity, resolving complete gene clusters and linking genomic predictions to metabolite production.

## Supporting information

Sup 3 detailed figure 2

Sup 1 Metadata

## Acknowledgments

The authors would like to thank CISIRO for providing 56 cyanobacterial genomes. This research used resources from the National Computational Infrastructure (NCI Australia), an NCRIS enabled capability supported by the Australian Government and the Australian Research Council Centre of Excellence in Synthetic Biology through project grant CE200100029.

## Supplementary material

Supplementary material 1 – Dataset

Supplementary material 2 – Additional tables and figures

Supplementary material 3 – High resolution phylogeny with additional metadata

## Data availability statement

All whole genome sequencing data generated for this study is available in the NCBI Sequence Read Archive under bioproject ID PRJNA924361. Public data was downloaded from the Sequence Read Archive and GenBank with accessions available in the supplementary material.

## CRediT statement

Rachel Mizzi: data curation, formal analysis, investigation, methodology, visualization, software and writing/reviewing of the manuscript. Verlaine Timms: conceptualization, data curation, funding acquisition, methodology, project administration, resources, software, supervision and writing/reviewing of the manuscript. Brett Neilan and Leanne Pearson: conceptualization, funding acquisition, supervision and writing/reviewing of the manuscript.

## References

Agarwal V, Miles ZD, Winter JM, Eustáquio AS, El Gamal AA, Moore BS. 2017. Enzymatic halogenation and dehalogenation reactions: pervasive and mechanistically diverse. Chemical reviews 117:5619–5674.

Al-Tebrineh J, Merrick C, Ryan D, Humpage A, Bowling L, Neilan BA. 2012. Community composition, toxigenicity, and environmental conditions during a cyanobacterial bloom occurring along 1,100 kilometers of the Murray River. Applied and Environmental Microbiology 78:263–272.

Alcorta J, Alarcón-Schumacher T, Salgado O, Díez B. 2020. Taxonomic Novelty and Distinctive Genomic Features of Hot Spring Cyanobacteria. Frontiers in genetics 11:568223.

Asnicar F, Thomas AM, Beghini F, Mengoni C, Manara S, Manghi P, Zhu Q, Bolzan M, Cumbo F, May U, et al. 2020. Precise phylogenetic analysis of microbial isolates and genomes from metagenomes using PhyloPhlAn 3.0. Nature Communications 11:2500–2500.

Balskus EP, Walsh CT. 2010. Genetic and molecular basis for sunscreen biosynthesis in cyanobacteria. Science (American Association for the Advancement of Science) 329:1653–1656.

Barón Sola Á, Gutiérrez Villanueva MA, del Campo FF, Sanz Alférez S. 2013. Characterization of Aphanizomenon ovalisporum amidinotransferase involved in cylindrospermopsin synthesis. MicrobiologyOpen (Weinheim) 2:447–458.

Bashir F, Bashir A, Bouaïcha N, Chen L, Codd GA, Neilan B, Xu W-L, Ziko L, Rajput VD, Minkina T, et al. 2023. Cyanotoxins, biosynthetic gene clusters, and factors modulating cyanotoxin biosynthesis. World journal of microbiology & biotechnology 39:241–241.

Belknap KC, Park CJ, Barth BM, Andam CP. 2020. Genome mining of biosynthetic and chemotherapeutic gene clusters in Streptomyces bacteria. Scientific Reports 10:2003–2003.

Benedetto Tiz D, Bagnoli L, Rosati O, Marini F, Sancineto L, Santi C. 2022. New Halogen-Containing Drugs Approved by FDA in 2021: An Overview on Their Syntheses and Pharmaceutical Use. Molecules 27:1643.

Blin K, Shaw S, Kloosterman AM, Charlop-Powers Z, Van Wezel GP, Medema MH, Weber T. 2021. AntiSMASH 6.0: Improving cluster detection and comparison capabilities. Nucleic acids research 49:W29–W35.

Calteau A, Fewer DP, Latifi A, Coursin T, Laurent T, Jokela J, Kerfeld CA, Sivonen K, Piel J, Gugger M. 2014. Phylum-wide comparative genomics unravel the diversity of secondary metabolism in Cyanobacteria. BMC Genomics 15:977–977.

Cameron ES, Sanchez S, Goldman N, Blaxter ML, Finn RD. 2024. Diversity and specificity of molecular functions in cyanobacterial symbionts. Scientific Reports 14:18658–18615.

Cao L, Do T, Link AJ. 2021. Mechanisms of action of ribosomally synthesized and posttranslationally modified peptides (RiPPs). Journal of industrial microbiology & biotechnology 48:1.

Cardona T. 2019. Thinking twice about the evolution of photosynthesis. Open biology 9:180246–180246.

Chen M-Y, Teng W-K, Zhao L, Hu C-X, Zhou Y-K, Han B-P, Song L-R, Shu W-S. 2021. Comparative genomics reveals insights into cyanobacterial evolution and habitat adaptation. The ISME Journal 15:211–227.

Chevrette MG, Gutiérrez-García K, Selem-Mojica N, Aguilar-Martínez C, Yañez-Olvera A, Ramos-Aboites HE, Hoskisson PA, Barona-Gómez F. 2020. Evolutionary dynamics of natural product biosynthesis in bacteria. Natural Product Reports 37:566–599.

Codd GA, Morrison LF, Metcalf JS. 2005. Cyanobacterial toxins: risk management for health protection. Toxicology and Applied Pharmacology 203:264–272.

Crowe C, Molyneux S, Sharma SV, Zhang Y, Gkotsi DS, Connaris H, Goss RJM. 2021. Halogenases: a palette of emerging opportunities for synthetic biology-synthetic chemistry and C-H functionalisation. Chemical Society reviews 5:9443–9481.

Deviram G, Mathimani T, Anto S, Ahamed TS, Ananth DA, Pugazhendhi A. 2020. Applications of microalgal and cyanobacterial biomass on a way to safe, cleaner and a sustainable environment. Journal of cleaner production 253:119770.

Ding CYG, Pang LM, Liang Z-X, Goh KKK, Glukhov E, Gerwick WH, Tan LT. 2018. MS/MS-based molecular networking approach for the detection of aplysiatoxin-related compounds in environmental marine cyanobacteria. Marine drugs 16:505.

Dittmann E, Gugger M, Sivonen K, Fewer DP. 2015. Natural product biosynthetic diversity and comparative genomics of the Cyanobacteria. Trends in microbiology (Regular ed.) 23:642–652.

Eusebio N, Rego A, Glasser NR, Castelo-Branco R, Balskus EP, Leão PN. 2021. Distribution and diversity of dimetal-carboxylate halogenases in cyanobacteria. BMC Genomics 22:1–633.

Ferrinho S, Connaris H, Mouncey NJ, Goss RJM. 2024. Compendium of metabolomic and genomic datasets for cyanobacteria: mined the gap. Water research (Oxford) 256:121492–121492.

Finn RD, Clements J, Eddy SR. 2011. HMMER web server: interactive sequence similarity searching. Nucleic acids research 39:W29–W37.

François RMM, Massicard J-M, Weissman KJ. 2025. The chemical ecology and physiological functions of type I polyketide natural products: the emerging picture. Natural Product Reports 42:324–358.

Gurevich A, Saveliev V, Vyahhi N, Tesler G. 2013. QUAST: quality assessment tool for genome assemblies. Bioinformatics 29:1072–1075.

Hackett JD, Wisecaver JH, Brosnahan ML, Kulis DM, Anderson DM, Bhattacharya D, Plumley FG, Erdner DL. 2013. Evolution of saxitoxin synthesis in Cyanobacteria and Dinoflagellates. Molecular biology and evolution 30:70–78.

He Y, Chen Y, Tao H, Zhou X, Liu J, Liu Y, Yang B. 2025. Secondary metabolites from cyanobacteria: source, chemistry, bioactivities, biosynthesis and total synthesis. Phytochemistry reviews 24:483–525.

Ishida K, Welker M, Christiansen G, Cadel-Six S, Bouchier C, Dittmann E, Hertweck C, Tandeau de Marsac N. 2009. Plasticity and evolution of aeruginosin biosynthesis in cyanobacteria. Applied and Environmental Microbiology 75:2017–2026.

Jenke-Kodama H, Dittmann E. 2009. Evolution of metabolic diversity: Insights from microbial polyketide synthases. Phytochemistry (Oxford) 70:1858–1866.

Katoch M, Mazmouz R, Chau R, Pearson LA, Pickford R, Neilan BA. 2016. Heterologous production of cyanobacterial mycosporine-like amino acids mycosporine-ornithine and mycosporine-lysine in Escherichia coli. Applied and Environmental Microbiology 82:6167–6173.

Katsuyama Y, Ohnishi Y. 2012. Type III Polyketide Synthases in Microorganisms. Methods in enzymology 515:359–377.

Khumalo MJ, Nzuza N, Padayachee T, Chen W, Yu J-H, Nelson DR, Syed K. 2020. Comprehensive analyses of cytochrome P450 monooxygenases and secondary metabolite biosynthetic gene clusters in Cyanobacteria. International Journal of Molecular Sciences 21:656.

Komarek J. 2016. A polyphasic approach for the taxonomy of cyanobacteria: principles and applications. European journal of phycology 51:346–353.

Krug D, Müller R. 2014. Secondary metabolomics: the impact of mass spectrometry-based approaches on the discovery and characterization of microbial natural products. Natural Product Reports 31:768–783.

Kultschar B. 2018. Secondary Metabolites in Cyanobacteria: IntechOpen.

Kundral S, Giang PD, Grundon LR, Supper JM, Khare SK, Bernhardt PV, Evans P, Bell SG, De Voss JJ. 2025. Characterisation of the thermophilic P450 CYP116B305 identified using metagenomics-derived sequence data from an Australian hot spring. Applied Microbiology and Biotechnology 109:133.

Letunic I, Bork P. 2019. Interactive Tree Of Life (iTOL) v4: recent updates and new developments. Nucleic acids research 47.

Li H, Ding W, Zhang Q. 2024. Discovery and engineering of ribosomally synthesized and post-translationally modified peptide (RiPP) natural products. RSC chemical biology 5:9–18.

Li K, Cai J, Su Z, Yang B, Liu Y, Zhou X, Huang J, Tao H. 2019. Glycosylated natural products from marine microbes. Frontiers in Chemistry 7.

Machado MJ, Jacinavicius FR, Médice RV, Dextro RB, Feitosa AMT, Weiss MB, Pellegrinetti TA, Cotta SR, Crnkovic CM, Fiore MF. 2025. Genetic and biochemical diversity of terpene biosynthesis in cyanobacterial strains from tropical soda lakes. Frontiers in microbiology 16.

McCranie EK, Bachmann BO. 2014. Bioactive oligosaccharide natural products. Natural Product Reports 31:126–142.

Mihali TK, Kellmann R, Muenchhoff J, Barrow KD, Neilan BA. 2008. Characterization of the gene cluster responsible for cylindrospermopsin biosynthesis. Applied and Environmental Microbiology 74:716–722.

Moffitt MC, Neilan BA. 2004. Characterization of the nodularin synthetase gene cluster and proposed theory of the evolution of cyanobacterial hepatotoxins. Applied and Environmental Microbiology 70:6353–6362.

Muenchhoff J, Siddiqui KS, Poljak A, Raftery MJ, Barrow KD, Neilan BA. 2010. novel prokaryotic l-arginine:glycine amidinotransferase is involved in cylindrospermopsin biosynthesis. The FEBS journal 277:3844–3860.

Mund NK, Liu Y, Chen S. 2022. Advances in metabolic engineering of cyanobacteria for production of biofuels. Fuel (Guildford) 322:124117.

Navarro-Munoz JC, Selem-Mojica N, Mullowney MW, Kautsar SA, Tryon JH, Parkinson EI, De Los Santos ELC, Yeong M, Cruz-Morales P, Abubucker S, et al. 2020. A computational framework to explore large-scale biosynthetic diversity. Nature chemical biology 16:60–68.

Nguyen L-T, Schmidt HA, Von Haeseler A, Minh BQ. 2015. IQ-TREE: a fast and effective stochastic algorithm for estimating maximum-likelihood phylogenies. Molecular biology and evolution 32:268.

Notonier S, Alexander M, Jayakody LN. 2016. An Overview of P450 Enzymes: Opportunity and Challenges in Industrial Applications. Enzyme engineering (Los Angeles, Calif.) 5.

Nowruzi B, Porzani SJ. 2021. Toxic compounds produced by cyanobacteria belonging to several species of the order Nostocales: A review. Journal of applied toxicology 41:510–548.

O’Neil JM, Davis TW, Burford MA, Gobler CJ. 2012. The rise of harmful cyanobacteria blooms: The potential roles of eutrophication and climate change. Harmful Algae 14:313–334.

Ongley SE, Bian X, Zhang Y, Chau R, Gerwick WH, Mu ller R, Neilan BA. 2013. High-titer heterologous production in E. coli of lyngbyatoxin, a protein kinase C activator from an uncultured marine cyanobacterium. ACS chemical biology 8:1888–1893.

Patel S. 2016. Drivers of bacterial genomes plasticity and roles they play in pathogen virulence, persistence and drug resistance. Infection, Genetics and Evolution 45:151–164.

Pattanaik B, Lindberg P. 2015. Terpenoids and their biosynthesis in cyanobacteria. Life 5:269–293.

Pearson LA, Dittmann E, Mazmouz R, Ongley SE, D’Agostino PM, Neilan BA. 2016. The genetics, biosynthesis and regulation of toxic specialized metabolites of cyanobacteria. Harmful Algae 54:98–111.

Pinheiro LB, Van Asten M, Antin L, Adams H, Qiu JY, Robinson M, DeLorenzo S, Holmes R, Hurd M, Tang R, et al. 2025. Interlaboratory performance study of cyanobacteria DNA reference materials using a qPCR format for monitoring cyanobacterial blooms. AWWA water science 7:n/a.

Popin RV, Alvarenga DO, Castelo-Branco R, Fewer DP, Sivonen K. 2021. Mining of cyanobacterial genomes indicates natural product biosynthetic gene clusters located in conjugative plasmids. Frontiers in microbiology 12:684565–684565.

Potter SC, Luciani A, Eddy SR, Park Y, Lopez R, Finn RD. 2018. HMMER web server: 2018 update. Nucleic acids research 46:W200–W204.

Rantala A, Fewer DP, Hisbergues M, Rouhiainen L, Vaitomaa J, Börner T, Sivonen K. 2004. Phylogenetic Evidence for the Early Evolution of Microcystin Synthesis. Proceedings of the National Academy of Sciences - PNAS 101:568–573.

Robert FO, Pandhal J, Wright PC. 2010. Exploiting cyanobacterial P450 pathways. Current Opinion in Microbiology 13:301–306.

Romanis CS, Pearson LA, Neilan BA. 2021. Cyanobacterial blooms in wastewater treatment facilities: Significance and emerging monitoring strategies. Journal of microbiological methods 180:106123.

Seemann T. 2014. Prokka: rapid prokaryotic genome annotation. Bioinformatics 30:2068–2069.

Soeriyadi AH, Mazmouz R, Pickford R, Al Sinawi B, Kellmann R, Pearson LA, Neilan BA. 2021. Heterologous Expression of an Unusual Ketosynthase, SxtA, Leads to Production of Saxitoxin Intermediates in Escherichia coli. Chembiochem : a European journal of chemical biology 22:845–849.

Strunecký O, Ivanova AP, Mareš J. 2023. An updated classification of cyanobacterial orders and families based on phylogenomic and polyphasic analysis. Journal of phycology 59:12–51.

Süssmuth RD, Mainz A. 2017. Nonribosomal Peptide Synthesis—Principles and Prospects. Angewandte Chemie International Edition 56:3770–3821.

Swain SS, Paidesetty SK, Padhy RN. 2017. Antibacterial, antifungal and antimycobacterial compounds from cyanobacteria. Biomedicine & Pharmacotherapy 90:760–776.

Timms VJ, Hassan KA, Pearson LA, Neilan BA. 2023. Cyanobacteria as a critical reservoir of the environmental antimicrobial resistome. Environmental microbiology 25:2266–2276.

Walter JM, Coutinho FH, Dutilh BE, Swings J, Thompson FL, Thompson CC. 2017. Ecogenomics and taxonomy of the cyanobacteria phylum. Frontiers in microbiology 8:2132–2132.

Wang H, Fewer DP, Holm L, Rouhiainen L, Sivonen K. 2014. Atlas of nonribosomal peptide and polyketide biosynthetic pathways reveals common occurrence of nonmodular enzymes. Proceedings of the National Academy of Sciences - PNAS 111:9259–9264.

Willis A, Woodhouse JN. 2020. Defining cyanobacterial species: diversity and description through genomics. Critical reviews in plant sciences 39:101–124.

Woodhouse JN, Kinsela AS, Collins RN, Bowling LC, Honeyman GL, Holliday JK, Neilan BA. 2016. Microbial communities reflect temporal changes in cyanobacterial composition in a shallow ephemeral freshwater lake. The ISME Journal 10:1337–1351.

Yang J, Wang Z, Li Z, Xu H, Xue B, Cao Y, Li Z, Li Y. 2025. Halogen-atom-aubstituted DOPA with enhanced wet adhesion and antioxidization ability. Biomacromolecules 26:2146–2156.

Yu G, Ge X, Li W, Ji L, Yang S. 2024. Interspecific cross-talk: The catalyst driving microbial biosynthesis of secondary metabolites. Biotechnology advances 76:108420.

Yu Y, Zhang Q, Deng Z. 2017. Parallel pathways in the biosynthesis of aminoglycoside antibiotics. F1000 research 6:723.

Zdouc Mitja M, Blin K, Louwen Nico LL, Navarro J, Loureiro C, Bader Chantal D, Bailey Constance B, Barra L, Booth Thomas J, Bozhüyük Kenan AJ, et al. 2025. MIBiG 4.0: advancing biosynthetic gene cluster curation through global collaboration. Nucleic acids research 53:D678–D690.

