## Supplementary figures and images for "Markers of biosynthetic innovation in the Cyanobacteria"

### Sup 3 detailed figure 2

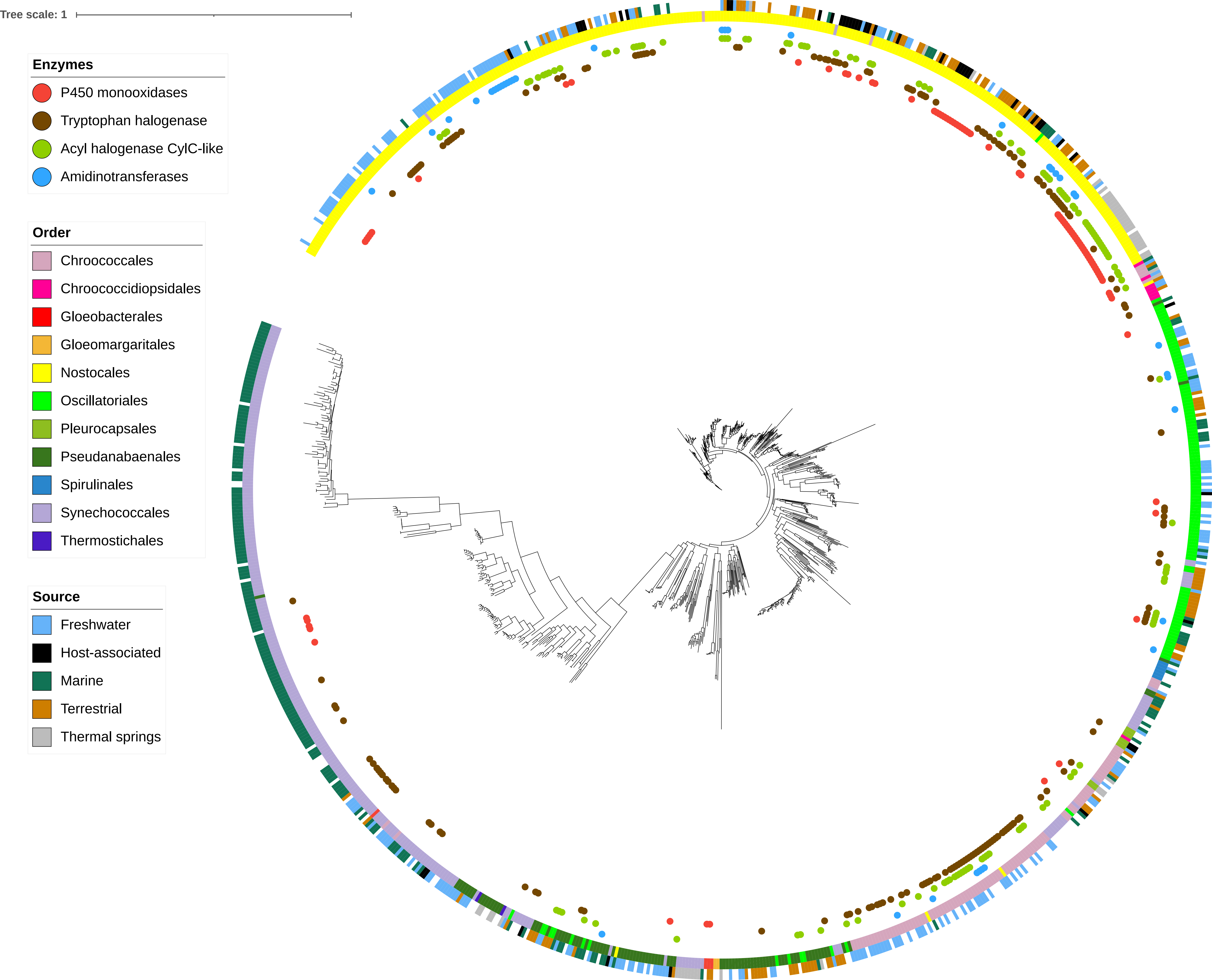
