## Supplementary material for "Markers of biosynthetic innovation in the Cyanobacteria": Sup 2 tables

Supplementary table 1 Average number of total BGCs by order and average number of each class of BGCs per genome within each order.

|  | NPRS | | Others | | PKS-NRP_Hybrids | | PKSI | | PKS Other | | RiPPs | | Saccharides | | Terpene | | Average no. BGCs per genome | No. genomes | Total BGCs per order |
| --- | --- | --- | --- | --- | --- | --- | --- | --- | --- | --- | --- | --- | --- | --- | --- | --- | --- | --- | --- |
| Chroococcales | 188 | 1.74 | 126 | 1.17 | 162 | 1.50 | 65 | 0.60 | 46 | 0.43 | 256 | 2.37 | 4 | 0.04 | 280 | 2.59 | 10.44 | 108 | 1127 |
| Chroococcidiopsidales | 27 | 3.38 | 7 | 0.88 | 11 | 1.38 | 4 | 0.50 | 1 | 0.13 | 22 | 2.75 | 1 | 0.13 | 9 | 1.13 | 10.25 | 8 | 82 |
| Gloeobacterales | 0 | 0.00 | 7 | 1.75 | 3 | 0.75 | 1 | 0.25 | 6 | 1.50 | 7 | 1.75 | 0 | 0.00 | 8 | 2.00 | 8 | 4 | 32 |
| Gloeomargaritales | 0 | 0.00 | 3 | 1.00 | 0 | 0.00 | 0 | 0.00 | 0 | 0.00 | 0 | 0.00 | 0 | 0.00 | 4 | 1.33 | 3.5 | 3 | 7 |
| Nostocales | 1077 | 3.30 | 541 | 1.66 | 616 | 1.89 | 98 | 0.30 | 561 | 1.72 | 1627 | 4.99 | 6 | 0.02 | 1005 | 3.08 | 16.97 | 326 | 5531 |
| Oscillatoriales | 254 | 2.03 | 130 | 1.04 | 123 | 0.98 | 27 | 0.22 | 19 | 0.15 | 277 | 2.22 | 2 | 0.02 | 258 | 2.06 | 8.72 | 125 | 1090 |
| Pleurocapsales | 17 | 2.13 | 4 | 0.50 | 20 | 2.50 | 12 | 1.50 | 3 | 0.38 | 32 | 4.00 | 0 | 0.00 | 24 | 3.00 | 14 | 8 | 112 |
| Pseudanabaenales | 140 | 1.51 | 56 | 0.60 | 50 | 0.54 | 13 | 0.14 | 36 | 0.39 | 85 | 0.91 | 2 | 0.02 | 236 | 2.54 | 6.65 | 93 | 618 |
| Spirulinales | 8 | 1.33 | 6 | 1.00 | 2 | 0.33 | 0 | 0.00 | 0 | 0.00 | 10 | 1.67 | 0 | 0.00 | 6 | 1.00 | 5.33 | 6 | 32 |
| Synechococcales | 60 | 0.23 | 41 | 0.16 | 41 | 0.16 | 17 | 0.07 | 74 | 0.29 | 157 | 0.61 | 0 | 0.00 | 584 | 2.27 | 3.79 | 257 | 974 |
| Thermostichales | 0 | 0.00 | 0 | 0.00 | 0 | 0.00 | 0 | 0.00 | 0 | 0.00 | 0 | 0.00 | 0 | 0.00 | 6 | 3.00 | 3 | 2 | 6 |
| Total | 1771 | 1.88 | 921 | 0.98 | 1028 | 1.09 | 237 | 0.25 | 746 | 0.79 | 2473 | 2.63 | 15 | 0.02 | 2420 | 2.57 | 10.24 | 940 | 9611 |

Supplementary table 2 number of BGC per source of cyanobacteria and the average of each BGC class per source

|  | **Freshwater** | | **Marine** | | **Thermal Springs** | | **Host-associated** | | **Terrestrial** | | **Unknown** | |
| --- | --- | --- | --- | --- | --- | --- | --- | --- | --- | --- | --- | --- |
|  | **Total** | **Average** | **Total** | **Average** | **Total** | **Average** | **Total** | **Average** | **Total** | **Average** | **Total** | **Average** |
| **NPRS**  **(n=1,771)** | 436 | 1.86 | 122 | 0.54 | 30 | 0.64 | 207 | 4.5 | 413 | 3.25 | 563 | 2.19 |
| **Others**  **(n=921)** | 228 | 0.97 | 83 | 0.37 | 47 | 1 | 97 | 2.11 | 194 | 1.53 | 272 | 1.06 |
| **PKS-NRP_Hybrids**  **(n=1,028)** | 235 | 1 | 87 | 0.38 | 53 | 1.13 | 118 | 2.57 | 182 | 1.43 | 353 | 1.37 |
| **PKSI**  **(n=237)** | 74 | 0.31 | 26 | 0.11 | 11 | 0.23 | 22 | 0.48 | 43 | 0.34 | 61 | 0.24 |
| **PKS Other**  **(n=746)** | 212 | 0.90 | 87 | 0.38 | 70 | 1.49 | 50 | 1.09 | 108 | 0.85 | 219 | 0.85 |
| **RiPPs**  **(n=2,473)** | 609 | 2.59 | 266 | 1.17 | 103 | 2.19 | 220 | 4.78 | 455 | 3.58 | 820 | 3.19 |
| **Saccarides**  **(n=15)** | 0 | 0 | 1 | 0.00 | 2 | 0.04 | 1 | 0.02 | 6 | 0.05 | 2 | 0.01 |
| **Terpene**  **(n=2,420)** | 621 | 2.64 | 516 | 2.27 | 104 | 2.21 | 158 | 3.43 | 337 | 2.65 | 684 | 2.66 |
| **Total BGCs in source** | 2418 | 10.29 | 1188 | 5.23 | 420 | 8.94 | 873 | 18.98 | 1738 | 13.69 | 2974 | 11.57 |
| **Standard deviation** | 5.79 | | 5.68 | | 5.57 | | 9.84 | | 7.84 | | 6.68 | |
| **Range** | 2-33 | | 1-42 | | 1-26 | | 1-41 | | 3-38 | | 1-29 | |

| A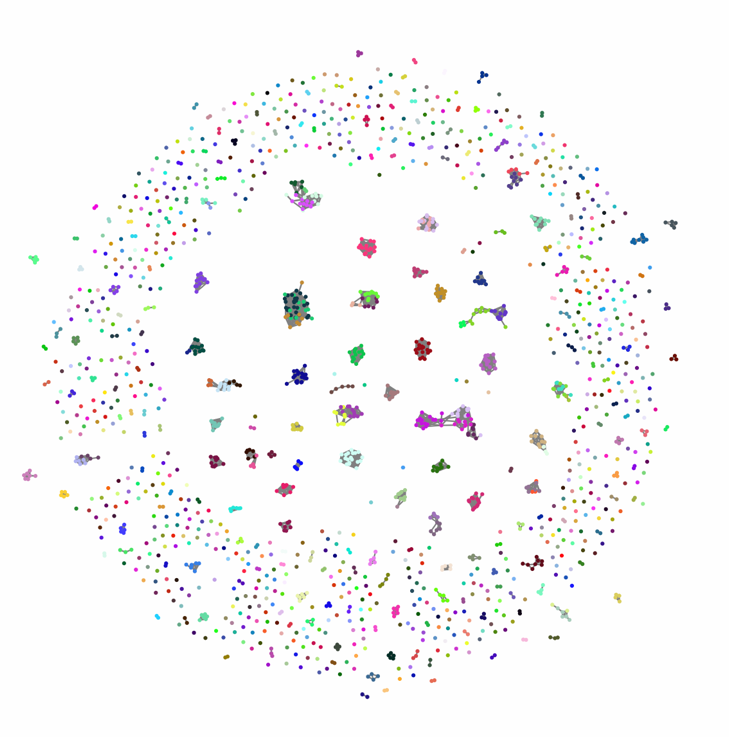 | B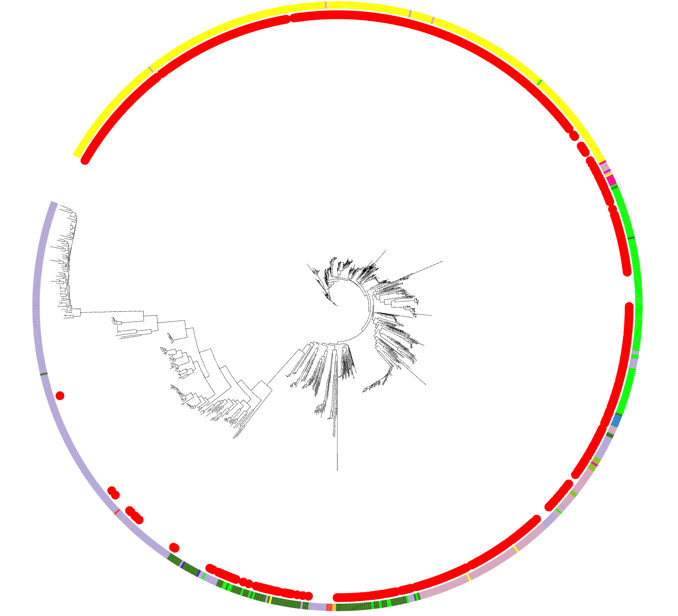 |
| --- | --- |
| Supplementary figure 1 A) network of NPRS clusters, B) phylogenomic distribution of NPRS clusters. 1,789 NPRS clusters were identified by antismash in the dataset. These clusters were organised into 975 families with 3,401 links and 736 singletons. Within the phylogenomic tree, these clusters are widespread in all cyanobacterial orders except for *Synechococcales*, in which they are sparser. Majority of the NRPS clusters identified had a predicted product of an NRPS (n=1,065), NRPS-like (n=538) or NRPS-NRPS-like (n=150). NRPS clusters were more abundant in genomes sourced from host-associated or terrestrial environments. These genome sources had an average of 3.98 and 3.15 clusters per genome, respectively. In contrast, genomes from marine and thermal spring environments had an average of one NRPS in half the genomes used in this study (n=0.56, n=0.567 respectively). No NRPS clusters were observed in genomes from the orders *Gloeobacterales*, *Gloeomargaritales* and *Thermostichales*. *Chroococcidiopsidales* and *Nostocales* had the highest average NRPS per genome at 3.38 and 3.3 respectively. *Chroococcidiopsidales* was the only order where this was on average the most frequently observed cluster per genome. | |

| A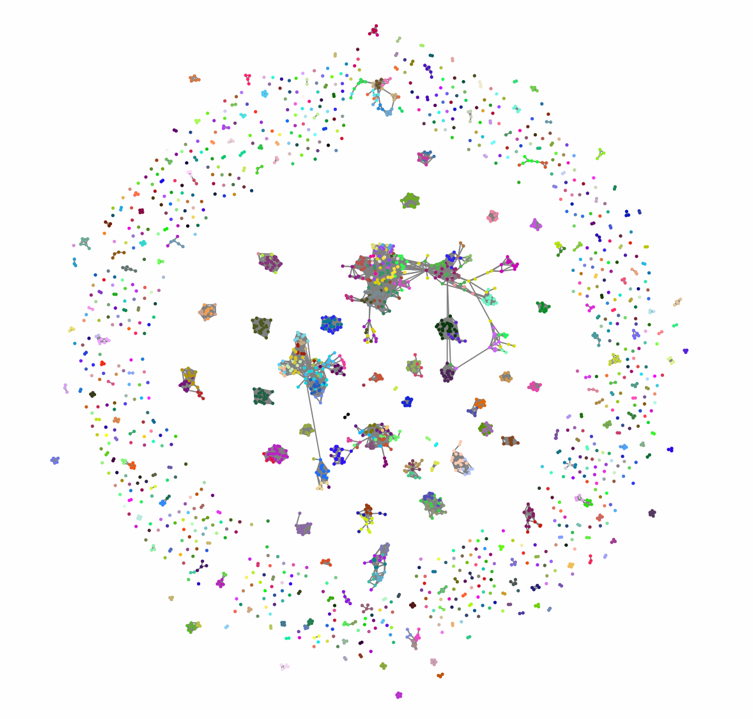 | B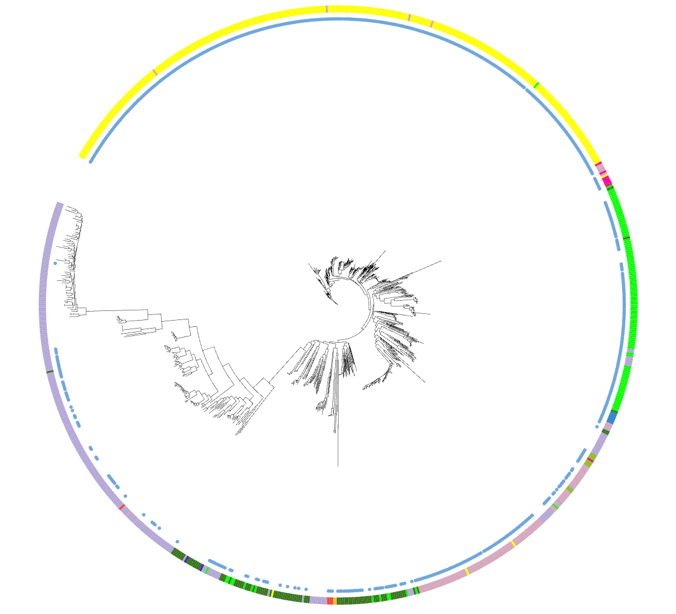 |
| --- | --- |
| Supplementary figure 2 A) Network of RiPPS clusters, B) phylogenomic distribution of RiPPS | |

The most commonly found cluster category was RiPPs with a total of 2,473 clusters that produce 55 different products. The network for RiPPs clusters consisted of 1,061 families, 7,654 links and 703 singletons. These clusters were common in all orders of cyanobacteria. 78% of products (n=1,953) fell into the lanthipeptide-class-v, RRE-containing, cyanobactin, lanthipeptide-class-ii, lassopeptide, microviridin and RiPP-like product categories at 708, 291, 257, 197, 180, 159 and 142 respectively.

Multiple RiPPs clusters were frequently found in host-associated and terrestrial genomes (average of n=4.78 and n=3.85 per genome respectively). RiPPs was the most frequently observed cluster for the *Nostocales, Oscillatoriales, Pleurocapsales* and *Spirulinales*. *Nostocales* and *Pleurocapsales* genomes had the highest average number of RiPPs per genome (n=4.99 and n=4 respectively). No RiPPs clusters were found in the *Gloeomargaritales* and *Thermostichales* genomes.

| A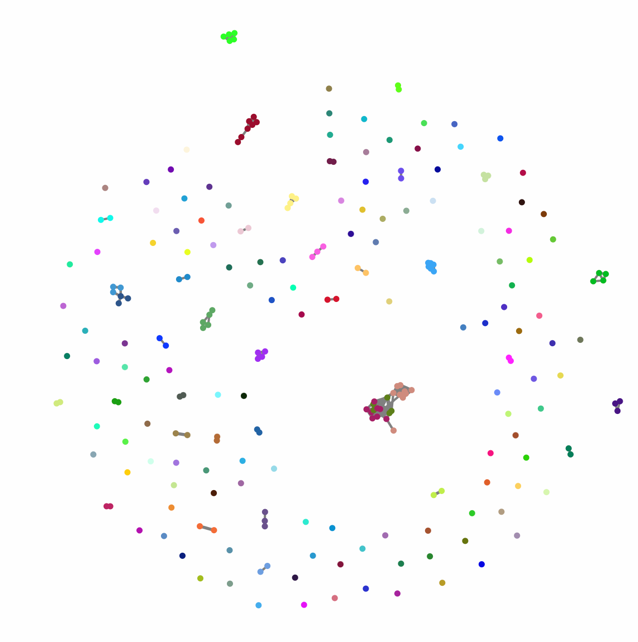 | B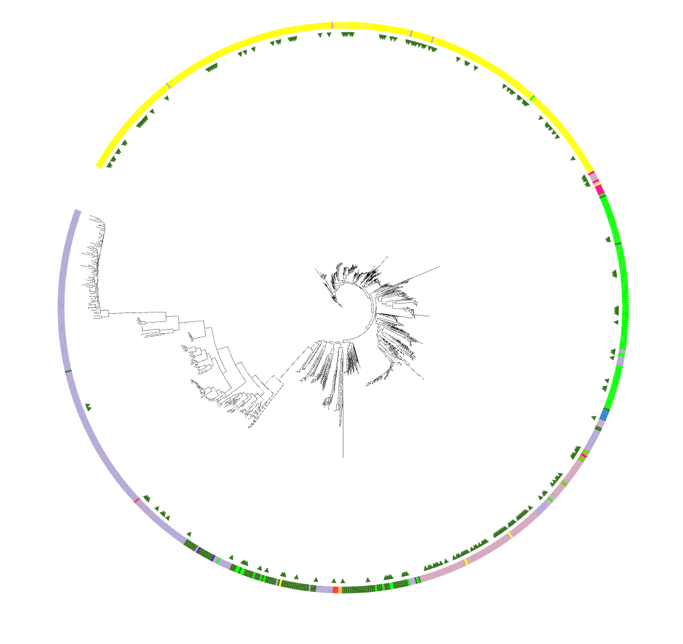 |
| --- | --- |
| Supplementary figure 3 A) Network of T1PKS clusters, B) phylogenomic distribution of PKSIs. Within the dataset, 237 Type I Polyketide Synthase (PKSI) BGCs were identified. These clusters consisted of 158 families with 190 links and 121 singletons. The sources with the highest frequencies of PKSI clusters were host-associated and terrestrial genomes (n=0.48 and n=0.34 average per genome respectively). *Pleurocapsales* (n=1.5) and *Chroococcales* (n=0.60) were the orders that had the highest average number of PKSI clusters per genome. These clusters were uncommon in *Synechococcales* (average of 0.06 per genome) and not found in any of the *Gloeomargaritales, Spirulinales* and *Thermostichales* genomes. | |

Supplementary table 3 Genome source, order, genus and species of NRPS clusters found 12 or less times throughout the dataset. A total of 18 NRPS clusters had 12 or less occurrences in the dataset. These included 11 cyclodipeptides (CDPs), 6 Non-alpha poly-amino acids (NAPAA) and one NRPS-thioamide-NRP products (Table 6). These BGCs came from 17 different genomes. Genomes were from freshwater (n=4), host-associated (n=4), marine (n=4), terrestrial (n=4), thermal springs (n=1) and one unknown source. Majority of the genomes were from *Nostocales* (n=13) the remainder were from Oscillatoriales (n=4) and Synechococcales (n=1). A single genome, GCF014132355, contained more than one rare NRPS cluster. This genome was sourced from a marine environment and is an *Okeania* spp. from *Oscillatoriales* and contains two cyclodipeptide clusters.

| **Product Prediction** | **Source** | **Order** |
| --- | --- | --- |
| CDPS (n=11) | Freshwater (n=2) | Nostocales |
|  | Host-associated (n=2) |  |
|  | Terrestrial (n=2) |  |
|  | Thermal springs |  |
|  | Marine (n=4) | Oscillatoriales |
| NAPAA (n=6) | Freshwater | Nostocales |
|  | Host-associated (n=2) |  |
|  | Terrestrial (n=2) |  |
|  | Freshwater | Synechococcales |
| NRPS.thioamide-NRP (n=1) | Unknown | Nostocales |

Supplementary table 4 Product prediction of RiPPS clusters observed four to 75 times throughout the dataset and the sources and orders these clusters are found in.

The remaining 539 RiPPs clusters produced 46 different products. The product breakdown for the remainder of the rare RiPPS clusters is summarised in Table 9. These less common RiPPS were not found in genomes from the orders *Gloeomargaritales*, *Spirulinales* or *Thermostichales*. Two genomes from *Gloeobacterales* genus, *Gloeobacter* species *violaceus* and *morelensis* contained a thioamitides thiopeptide that did not appear to be specific to these genomes.

| **Product prediction** | **Source** | **Order** |
| --- | --- | --- |
| lanthipeptide-class-v.microviridin (n=4) | Host-associated | Nostocales |
|  | Marine | Oscillatoriales |
|  | Terrestrial | Nostocales |
|  | Terrestrial | Oscillatoriales |
| cyanobactin.thiopeptide (n=4) | Freshwater | Chroococcales |
|  | Terrestrial (n=2) | Nostocales |
|  | Freshwater |  |
| ranthipeptide (n=5) | Freshwater | Chroococcidiopsidales |
|  | Terrestrial |  |
|  | Unknown (n=2) |  |
|  | Unknown (n=1) | Synechococcales |
|  | Unknown | Chroococcidiopsidales |
| proteusin.RiPP-like.spliceotide.lanthipeptide-class-ii (n=6) | Freshwater (n=2) | Nostocales |
|  | Terrestrial (n=1) |  |
|  | Unknown (n=3) |  |
| thioamitides.lanthipeptide-class-ii (n=7) | Freshwater (n=6) | Oscillatoriales |
|  | Unknown (n=1) |  |
| RRE-containing.lanthipeptide-class-ii (n=9) | Terrestrial (n=1) | Nostocales |
|  | Unknown (n=8) |  |
| thiopeptide.thioamitides (n=10) | Terrestrial | Gloeobacterales |
|  | Unknown |  |
|  | Freshwater (n=2) | Nostocales |
|  | Marine |  |
|  | Terrestrial (n=3) |  |
|  | Freshwater (n=2) | Oscillatoriales |
| lassopeptide.RRE-containing (n=11) | Terrestrial | Chroococcales |
|  | Freshwater | Nostocales |
|  | Host-associated |  |
|  | Marine |  |
|  | Terrestrial |  |
|  | Unknown |  |
|  | Freshwater | Oscillatoriales |
|  | Marine |  |
|  | Unknown | Pleurocapsales |
|  | Marine | Synechococcales |
|  | Unknown |  |
| cyanobactin.lanthipeptide-class-v (n=13) | Unknown | Chroococcales |
|  | Marine | Nostocales |
|  | Unknown (n=11) | Nostocales |
| lanthipeptide-class-v.lanthipeptide-class-ii (n=14) | Freshwater (n=3) |  |
|  | Host-associated |  |
|  | Terrestrial |  |
|  | Unknown (n=4) |  |
|  | Freshwater | Oscillatoriales |
|  | Marine (n=2) |  |
|  | Unknown |  |
|  | Host-associated | Pleurocapsales |
| thioamitides (n=14) | Unknown | Chroococcales |
|  | Freshwater | Nostocales |
|  | Host-associated (n=3) | Nostocales |
|  | Marine (n=2) | Nostocales |
|  | Thermal springs | Nostocales |
|  | Terrestrial (n=4) | Oscillatoriales |
|  | Unknown | Oscillatoriales |
|  | Unknown | Synechococcales |
| spliceotide (n=14) | Freshwater (n=4) | Chroococcales |
|  | Unknown (n=6) | Chroococcales |
|  | Freshwater | Nostocales |
|  | Host-associated (n=2) |  |
|  | Unknown | Pseudanabaenales |
| proteusin.RiPP-like (n=14) | Freshwater | Chroococcales |
|  | Unknown |  |
|  | Host-associated (n=5) | Nostocales |
|  | Terrestrial (n=2) |  |
|  | Thermal springs |  |
|  | Unknown (n=3) |  |
|  | Marine | Synechococcales |
| thiopeptide (n=25) | Freshwater (n=4) | Chroococcales |
|  | Marine |  |
|  | Unknown (n=2) |  |
|  | Freshwater (n=2) | Nostocales |
|  | Host-associated (n=3) |  |
|  | Marine |  |
|  | Terrestrial (n=5) |  |
|  | Unknown (n=5) |  |
|  | Freshwater | Oscillatoriales |
|  | Unknown |  |
| cyanobactin.RRE-containing (n=26) | Freshwater (n=4) | Chroococcales |
|  | Marine |  |
|  | Terrestrial |  |
|  | Unknown | Chroococcales |
|  | Freshwater (n=2) | Nostocales |
|  | Marine |  |
|  | Terrestrial (n=4) |  |
|  | Unknown (n=9) |  |
|  | Marine (n=3) | Oscillatoriales |
| LAP.thiopeptide (n=26) | Freshwater (n=10) | Nostocales |
|  | Host-associated |  |
|  | Marine |  |
|  | Terrestrial |  |
|  | Thermal springs |  |
|  | Unknown (n=11) |  |
|  | Marine | Synechococcales |
| redox-cofactor (n=30) | Marine | Chroococcidiopsidales |
|  | Freshwater | Nostocales |
|  | Host-associated |  |
|  | Marine |  |
|  | Terrestrial (n=8) |  |
|  | Thermal springs |  |
|  | Unknown (n=14) |  |
|  | Freshwater | Pleurocapsales |
|  | Terrestrial | Pseudanabaenales |
|  | Unknown | Synechococcales |
| proteusin.lanthipeptide-class-ii (n=46) | Freshwater | Chroococcales |
|  | Unknown (n=2) |  |
|  | Freshwater (n=5) | Nostocales |
|  | Host-associated (n=3) |  |
|  | Marine (n=2) |  |
|  | Terrestrial (n=8) |  |
|  | Unknown (n=13) |  |
|  | Freshwater (n=2) | Oscillatoriales |
|  | Unknown (n=2) |  |
|  | Host-associated | Pleurocapsales |
|  | Terrestrial (n=3) | Pseudanabaenales |
| lanthipeptide-class-v.RRE-containing.spliceotide (n=46) | Freshwater (n=2) | Chroococcales |
|  | Terrestrial (n=2) |  |
|  | Unknown (n=2) |  |
|  | Freshwater (n=6) | Nostocales |
|  | Host-associated (n=4) |  |
|  | Marine (n=2) |  |
|  | Terrestrial (n=3) |  |
|  | Unknown (n=23) |  |
|  | Unknown |  |
|  | Freshwater | Pseudanabaenales |
|  | Unknown | Synechococcales |
| proteusin (n=51) | Freshwater | Chroococcales |
|  | Freshwater (n=2) | Chroococcidiopsidales |
|  | Terrestrial |  |
|  | Unknown (n=2) |  |
|  | Freshwater (n=6) | Nostocales |
|  | Host-associated (n=5) |  |
|  | Marine |  |
|  | Terrestrial (n=6) |  |
|  | Thermal springs |  |
|  | Unknown (n=13) |  |
|  | Marine | Oscillatoriales |
|  | Terrestrial |  |
|  | Unknown |  |
|  | Marine | Pseudanabaenales |
|  | Terrestrial |  |
|  | Unknown |  |
|  | Marine (n=4) | Synechococcales |
|  | Terrestrial |  |
|  | Unknown (n=2) |  |
| RRE-containing.spliceotide (n=65) | Freshwater (n=2) | Chroococcales |
|  | Unknown |  |
|  | Freshwater (n=20) | Nostocales |
|  | Host-associated (n=4) |  |
|  | Marine (n=4) |  |
|  | Terrestrial (n=12) |  |
|  | Thermal springs |  |
|  | Unknown (n=11) |  |
|  | Host-associated | Oscillatoriales |
|  | Marine (n=2) |  |
|  | Unknown (n=4) |  |
|  | Thermal springs (n=2) | Pleurocapsales |
|  | Marine | Pseudanabaenales |
| LAP (n=75) | Terrestrial | Chroococcales |
|  | Freshwater (n=10) | Nostocales |
|  | Host-associated (n=5) |  |
|  | Marine (n=4) |  |
|  | Terrestrial (n=8) |  |
|  | Thermal springs |  |
|  | Unknown (n=25) |  |
|  | Marine (n=3) | Oscillatoriales |
|  | Unknown |  |
|  | Thermal springs (n=2) | Pleurocapsales |
|  | Freshwater | Pseudanabaenales |
|  | Unknown |  |
|  | Marine (n=10) | Synechococcales |
|  | Unknown (n=3) |  |

| A  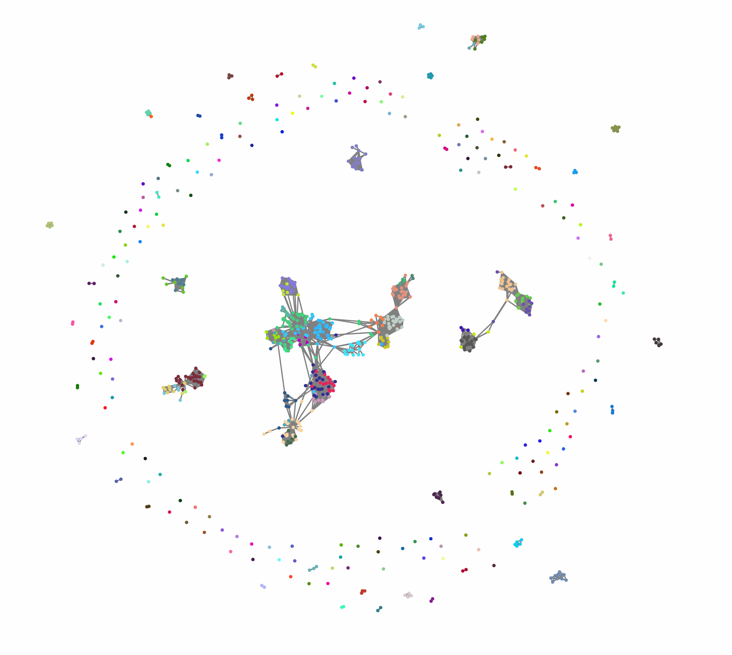 | B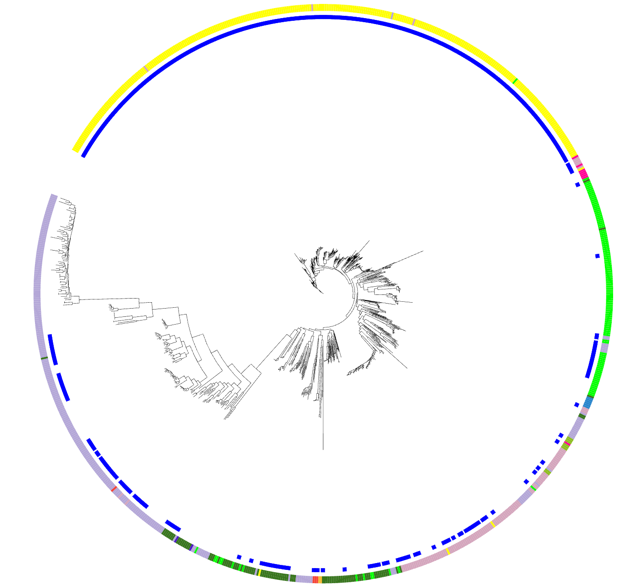 |
| --- | --- |
| Supplementary figure 4 A) Network of other PKS others clusters, B) phylogenomic distribution of other PKS clusters. 746 BGCs were identified as other PKS and produced 12 different products. This class was categorised into 229 families with 3,469 links and 148 singletons. Three of these products make up the bulk of the clusters, with T1PKS-hglE-KS, hglE-KS and T3PKS products found 330, 260 and 118 times in the dataset, respectively. Other PKS clusters were found in all *Nostocales* genomes and are distributed throughout all other orders more sparsely except for the *Oscillatoriales*, in which they appear to be less frequent. Genomes sourced from thermal springs had the highest occurrence of PKS other clusters (average of n=1.49 per genome) and marine-sourced genomes were the least likely to contain these clusters (average of n=0.38 per genome). No *Spirulinales*, *Gloeomargaritales* or *Thermostichales* in this dataset contained any PKS other clusters. The *Nostocales* genomes contained the highest average number of PKS other clusters per genome at 1.72. | |

Supplementary table 5. Genome source and order of PKS other clusters found 11 or less times throughout the dataset. Ten PKS clusters classified as ‘other’ were found in 11 or less occurrences within the dataset (total = 39). T3PKS.T1PKS producing clusters were found 11 times, transAT-PKS-like and transAT-PKS were found seven times and T2PKS were found in four instances. Five less common PKS were also found including two transAT-PKS.T1PKS.hglE-KS, transAT-PKS-like.T1PKS, T3PKS.T1PKS.hglE-KS and PKS-like.T3PKS.T1PKS and one transAT-PKS.T3PKS.PKS-like.T1PKS, T1PKS.hglE-KS.T3PKS and a T1PKS.transAT-PKS.T3PKS.PKS-like (Table 7). These rare PKS clusters were from 10 genomes from freshwater, six from marine, 11 from terrestrial, two from thermal springs, two host-associated and six from unknown sources. The orders that these genomes were from was; nine from *Chroococcales*, one from *Chroococcidiopsidales*, two *Gloeobacterales*, 11 *Nostocales*, two *Oscillatoriales*, two *Pleurocapsales*, nine *Pseudanabaenales*, and one *Synechococcales*.

| **Product** | **Source** | **Order** |
| --- | --- | --- |
| PKS-like.T3PKS.T1PKS | Terrestrial (n=2) | Nostocales |
| T2PKS | Thermal springs (n=2) | Chroococcales |
|  | Marine | Chroococcales |
|  | Freshwater | Chroococcidiopsidales |
| T3PKS.T1PKS | Unknown | Chroococcales |
|  | Freshwater (n=3) | Chroococcales |
|  | Unknown (n=2) | Chroococcales |
|  | Terrestrial (n=4) | Nostocales |
|  | Host-associated | Pleurocapsales |
| T3PKS.T1PKS.hglE-KS | Marine | Nostocales |
|  | Marine | Oscillatoriales |
| transAT-PKS | Freshwater | Nostocales |
|  | Terrestrial | Nostocales |
|  | Freshwater (n=4) | Pseudanabaenales |
|  | Terrestrial | Pseudanabaenales |
| transAT-PKS-like | Terrestrial | Nostocales |
|  | Unknown | Pleurocapsales |
|  | Host-associated | Pleurocapsales |
|  | Terrestrial (n=2) | Pseudanabaenales |
|  | Marine | Pseudanabaenales |
|  | Unknown | Synechococcales |
| transAT-PKS-like.T1PKS | Host-associated | Nostocales |
|  | Freshwater | Nostocales |
| transAT-PKS.T1PKS.hglE-KS | Unknown | Gloeobacterales |
|  | Terrestrial | Gloeobacterales |
| transAT-PKS.T3PKS.PKS-like | Marine | Pseudanabaenales |
| transAT-PKS.T3PKS.PKS-like.T1PKS | Marine | Oscillatoriales |

| A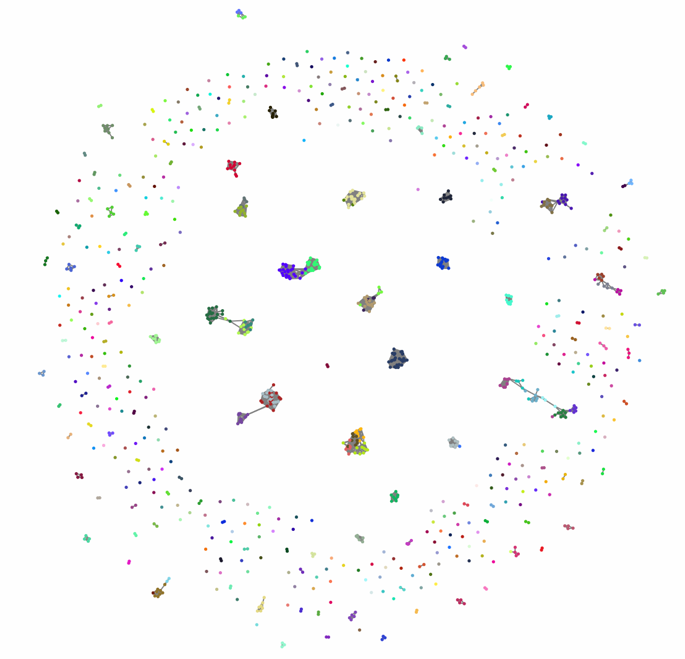 | B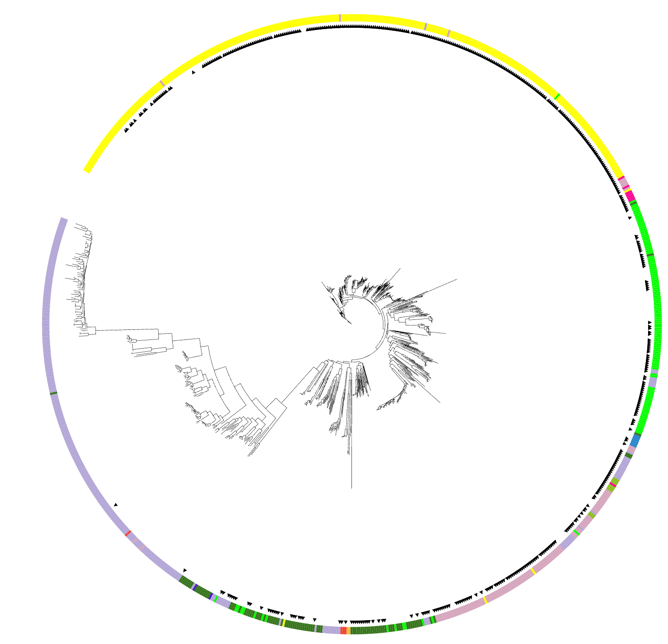 |
| --- | --- |
| Supplementary figure 5 A) Network of PKS-NRP hybrid clusters, B) phylogenomic distribution of PKS-NRP clusters. These clusters were spread throughout the phylogeny in all clades except for the *Synechococcales* clade, which only contained two of these clusters. These clusters were organised into 511 families with 2,803 families and 356 singletons.  1,028 PKS-NRP hybrid BGCs producing 22 different products were found in the dataset. These clusters were spread throughout the phylogeny in all clades except for the *Synechococcales* clade, which only contained two of these clusters. These clusters were organised into 511 families with 2,803 families and 356 singletons. The most common products were T1PKS.NRPS (n=516), T1PKS.NRPS.NRPS-like (n=232), T1PKS.NRPS-like (n=172), T1PKS.NRPS.hglE-KS (n=41), T1PKS.T3PKS.NRPS-like (n=28) and T3PKS.PKS-like.transAT-PKS.NRPS.T1PKS (n=12). The sources that this class was most frequently found in was host-associated (average 2.56 per genome) and it was least common in marine sources (average 0.38 per genome). The orders *Pleurocapsales*, *Nostocales* and *Chroococcales* contained the most PKS-NRP hybrid clusters, with an average of 2.5, 1.89 and 1.5 clusters per genome, respectively. PKS-NRP hybrids were found in *Thermostichales* or *Gloeomargaritales* genomes in this dataset and they were rare in *Synechococcales*, *Spirulinales* and *Pseudanabaenales* with an average of 0.16, 0.33 and 0.54 per genome respectively. | |

Supplementary table 6. Genome source, order, genus and species of PKS-NRP hybrid found 12 or less times throughout the dataset.

1,028 PKS-NRP hybrid BGCs producing 22 different products were found in the dataset. The most common products were T1PKS.NRPS (n=516), T1PKS.NRPS.NRPS-like (n=232), T1PKS.NRPS-like (n=172), T1PKS.NRPS.hglE-KS (n=41), T1PKS.T3PKS.NRPS-like (n=28) and T3PKS.PKS-like.transAT-PKS.NRPS.T1PKS (n=12). The sources that this class was most frequently found in was host-associated (average 2.56 per genome) and it was least common in marine sources (average 0.38 per genome). The orders *Pleurocapsales*, *Nostocales* and *Chroococcales* contained the most PKS-NRP hybrid clusters, with an average of 2.5, 1.89 and 1.5 clusters per genome, respectively. PKS-NRP hybrids were found in *Thermostichales* or *Gloeomargaritales* genomes in this dataset and they were rare in *Synechococcales*, *Spirulinales* and *Pseudanabaenales* with an average of 0.16, 0.33 and 0.54 per genome respectively. The remaining 27 PKS-NRP hybrid clusters produced 16 different products from 23 different genomes. These cluster products were only present only one to four times in the dataset. These rarer PKS-NRP hybrid clusters were only found in genomes from *Oscillatoriales, Nostocales, Pseudanabaenales* and *Chroococcales.*

| **Product Prediction** | **Source** | **Order** |
| --- | --- | --- |
| hglE-KS.NRPS-like | Terrestrial | Oscillatoriales |
| NRPS.hglE-KS | Terrestrial | Oscillatoriales |
| prodigiosin.NRPS | Freshwater | Nostocales |
| T1PKS.hglE-KS.NRPS-like | Terrestrial | Nostocales |
|  | Unknown | Nostocales |
| T1PKS.NRPS-like.CDPS | Unknown | Oscillatoriales |
| T1PKS.NRPS.CDPS | Terrestrial (n=2) | Nostocales |
| T1PKS.NRPS.hglE-KS.T3PKS | Marine | Oscillatoriales |
|  | Host-associated | Oscillatoriales |
| T1PKS.NRPS.NRPS-like.CDPS | Marine | Oscillatoriales |
| T1PKS.NRPS.NRPS-like.transAT-PKS | Host-associated | Nostocales |
| T1PKS.NRPS.PKS-like | Host-associated | Oscillatoriales |
| T1PKS.NRPS.T3PKS.PKS-like | Unknown | Oscillatoriales |
|  | Marine (n=3) | Oscillatoriales |
| T2PKS.NRPS-like | Freshwater | Pseudanabaenales |
| transAT-PKS-like.NRPS-like | Marine | Oscillatoriales |
| transAT-PKS-like.T1PKS.NRPS | Unknown | Nostocales |
| transAT-PKS-like.T1PKS.NRPS-like | Freshwater | Oscillatoriales |
|  | Unknown | Oscillatoriales |
|  | Freshwater (n=2) | Chroococcales |
| transAT-PKS.NRPS-like | Unknown | Chroococcales |
|  | Freshwater (n=2) | Oscillatoriales |

| A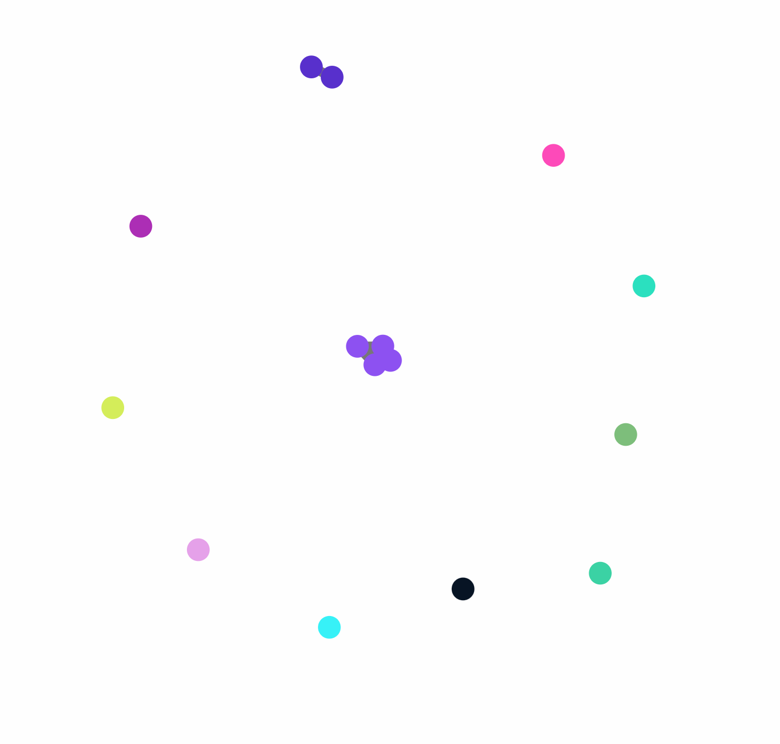 | B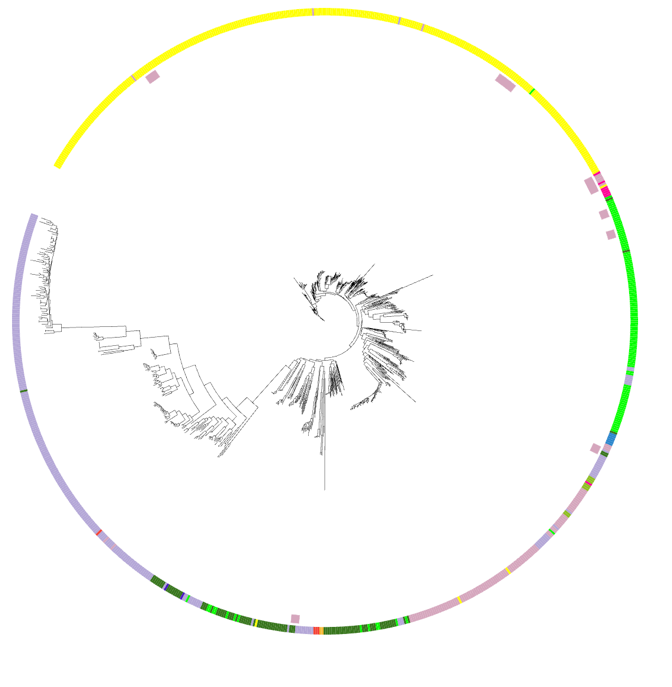 |
| --- | --- |

Supplementary figure 6 A) Network of saccharides clusters, B) phylogenomic distribution of saccharide clusters. Saccharides were the rarest type of BGC with only 15 clusters found from 15 different genomes. These were arranged into 11 families with seven links and nine singletons. Of these, seven produced an oligosaccharide and eight produced an amglyccycl. Sacccharide clusters were only found in *Chroococcales*, *Chroococcidiopsidales*, *Nostocales*, *Oscillatoriales* and *Pseudanabaenales.* No two genomes within the dataset contained more than one saccharide producing BGC. The BGCs within the central cluster in the network are from four different genomes that cluster tightly on the phylogeny within the *Nostocales*.

| A 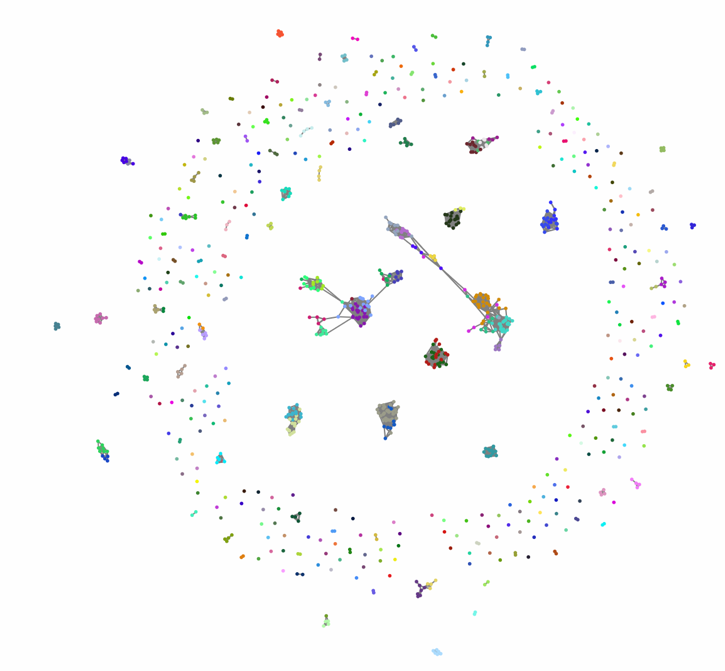 | B 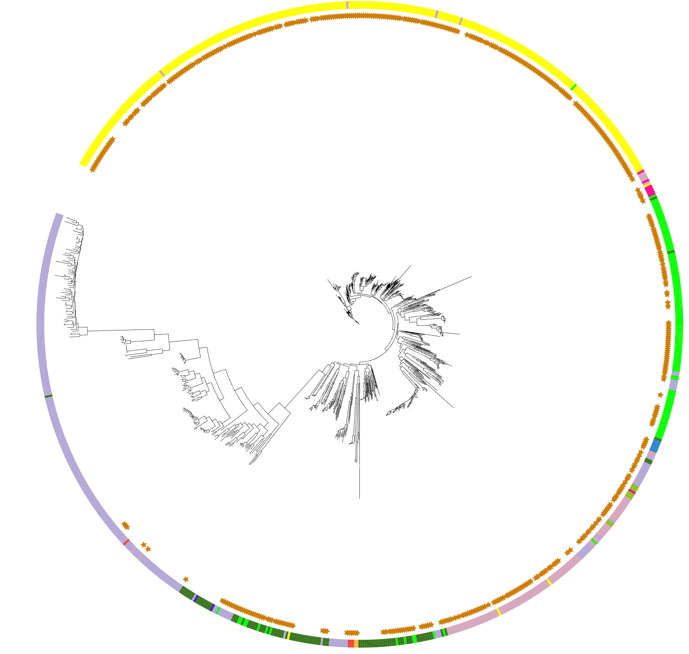 |
| --- | --- |
| Supplementary figure 7 A) Network of other clusters B) phylogenomic distribution of other clusters.  Other BGCs, a very broad category, were found 921 times within the dataset and produced the highest number of predicted products (n=135). The network consisted of 413 families of clusters, 2,780 links and 278 singletons. Clusters categorized as other were widespread in all orders except for *Synechococcales*. The most common products consisted of ladderane (n=101), NRPS.betalactone (n=63), arylpolyene (n=59), resorcinol (n=57), indole (n=56), siderophore (n=48), phosphonate (n=44), NRPS.microviridin (n=42), resorcinol.hglE-KS and NRPS.betalactone.microviridin (both n=32). Of the remaining 388 clusters, 150 products occurred a total of 11-28 instances, 53 occurred 2-9 times and 62 products were only found once in the dataset. Other clusters ranged from an average of 0.37 per marine-sourced genome and 2.11 clusters per host-associated genome. *Gloeobacterales* had the highest average frequency of these clusters with an average of 1.75 per genome, *Synechococcales* had the lowest (n=0.16) apart from *Thermostichales* which had none observed. | |

| A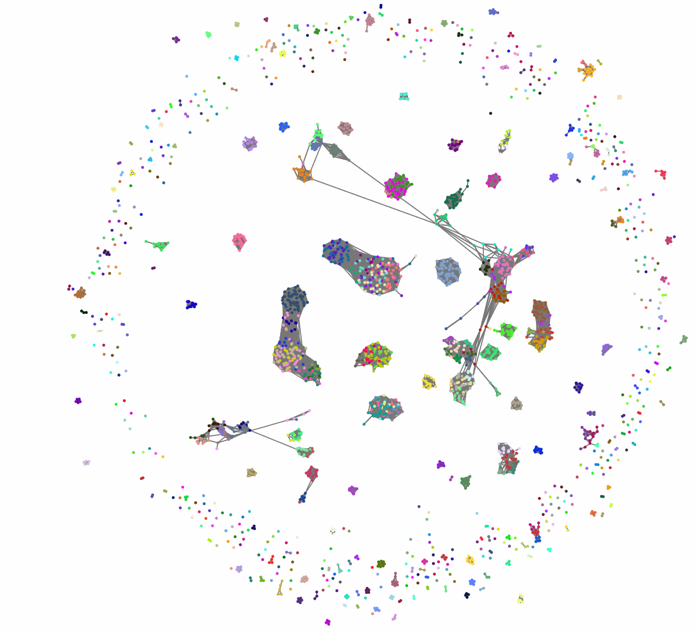 | B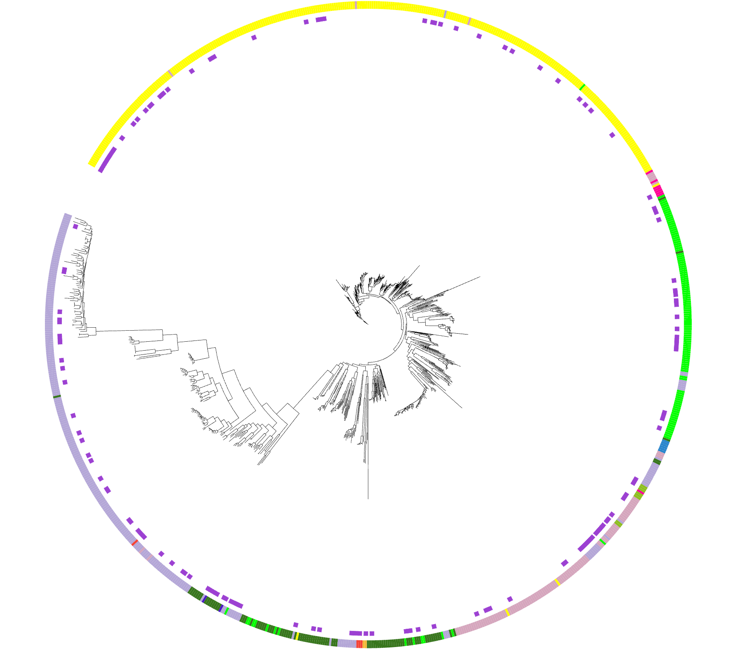 |
| --- | --- |

Supplementary figure 8 A) Network of terpene clusters, B) phylogenomic distribution of terpene clusters. 2,420 Terpene BGCs were found in the dataset and these were evenly distributed throughout the phylogeny. Terpene clusters were organised into 679 families with 23,810 links and 399 singletons. No further breakdown of the product type was available for this class. Terpene clusters were the most frequently observed class for the orders *Chroococcales, Gloeobacterales, Gloeomargaritales, Pseudanabaenales, Synechococcales* and the only detected clusters in the *Thermostichales* genomes. This cluster type was most frequent in host-associated genomes (n=3.44), and in the orders *Pleurocapsales, Thermostichales* and *Nostocales* (average of n=3, n=3, n=3.08 per genome).

Supplementary table 7 the number of BGCs in *Synechococcales* genomes in the dataset and their sources.

| **Genome Source** | **No. BGCs** | **No. genomes** | **Average no. BGCs per genome** |
| --- | --- | --- | --- |
| **Freshwater** | 85 | 24 | 3.54 |
| **Marine** | 573 | 168 | 3.41 |
| **Thermal Springs** | 11 | 9 | 1.22 |
| **Host-associated** | 25 | 5 | 5 |
| **Terrestrial** | 72 | 11 | 6.55 |
| **Unknown** | 208 | 40 | 5.2 |
| **Total** | 974 | 257 | 3.79 |

Supplementary table 8 BGCs that contained at least one cytochrome 450 monoxidase enzyme

| **BigScape category** | **Product Prediction** | **Count** |
| --- | --- | --- |
| NRPS | NRPS | 12 |
|  | NRPS-like | 3 |
|  | NRPS.NRPS-like | 1 |
| Others | ladderane | 1 |
|  | phenazine | 1 |
|  | resorcinol | 1 |
|  | thiopeptide.terpene.NRPS.thioamitides.T1PKS | 2 |
| PKS-NRP_Hybrids | T1PKS.NRPS | 22 |
|  | T1PKS.NRPS.hglE-KS | 1 |
|  | T1PKS.NRPS.hglE-KS.T3PKS | 2 |
|  | T1PKS.NRPS.NRPS-like | 5 |
|  | T1PKS.transAT-PKS.hglE-KS | 4 |
| PKSI | T1PKS | 1 |
| PKS other | T2PKS | 3 |
|  | T3PKS | 5 |
| RiPPs | lanthipeptide-class-v | 1 |
|  | lanthipeptide-class-v.lanthipeptide-class-ii | 2 |
|  | lassopeptide | 1 |
|  | microviridin | 1 |
|  | RRE-containing.spliceotide | 4 |
|  | spliceotide | 1 |
| Terpene | terpene | 48 |
