## Supplementary material for "Markers of biosynthetic innovation in the Cyanobacteria": Sup 1 Metadata

Genome ID
GCF_003003925.1_ASM300392v1_genomic
GCF_003007785.1_ASM300778v1_genomic
GCF_003864295.1_ASM386429v1_genomic
GCF_003864315.1_ASM386431v1_genomic
GCF_003003885.1_ASM300388v1_genomic
GCF_002368355.1_ASM236835v1_genomic
GCF_017312365.1_ASM1731236v1_genomic
GCF_001904655.1_ASM190465v1_genomic
GCF_000167195.1_ASM16719v1_genomic
GCF_000169335.1_ASM16933v1_genomic
GCF_000231425.2_ASM23142v3_genomic
GCA_000017845.1_ASM1784v1_genomic.fna
GCA_002813895.1_ASM281389v1_genomic.fna
GCA_947331815.1_uoEpiScrs1.Cyanobacterium_sp_1.1_genomic.fna
GCF_000317675.1_ASM31767v1_genomic
GCF_001747005.1_ASM174700v1_genomic
GCF_002736005.1_ASM273600v1_genomic
GCF_009710015.1_ASM971001v1_genomic
GCF_014697365.1_ASM1469736v1_genomic
GCF_015207825.1_ASM1520782v1_genomic
GCA_000317655.1_ASM31765v1_genomic.fna
GCF_007904085.1_ASM790408v1_genomic
GCF_000332235_1.chr
GCF_001548095.1_Gm3709_assembly_1.0_genomic
GCF_001548115.1_Gm3709_assembly_1.0_genomic
GCF_000317555.1_ASM31755v1_genomic
GCF_000332035.1_ASM33203v1_genomic
GCF_002964865.1_ASM296486v1_genomic
GCF_015207655.1_ASM1520765v1_genomic
GCF_000021825.1_ASM2182v1_genomic
GCF_000147335.1_ASM14733v1_genomic
GCF_000317635.1_ASM31763v1_genomic
GCA_019704275.1_ASM1970427v1_genomic.fna
GCA_020905635.1_ASM2090563v1_genomic.fna
GCA_021172085.1_ASM2117208v1_genomic.fna
GCA_026222535.1_ASM2622253v1_genomic.fna
GCF_000010625.1_ASM1062v1_genomic
GCF_000297435.1_ASM29743v1_genomic
GCF_000312165_1.scaf
GCF_000312185_1.scaf
GCF_000312205_1.scaf
GCF_000312225_1.scaf
GCF_000312245_1.scaf
GCF_000312265_1.scaf
GCF_000312285_1.scaf
GCF_000312725_1.scaf
GCF_000330925.1_MicAerT1.0_genomic

GCF_000332585.1_MicAerD1.0_genomic
GCF_00037995.1.scaf
GCF_000412595.1_spc777-v1_genomic
GCF_000787675.1_ASM78767v1_genomic
GCF_000981785.2_ASM98178v2_genomic
GCF_001578075.1_ASM157807v1_genomic
GCF_001704955.2_ASM170495v2_genomic
GCF_001725075.1_ASM172507v1_genomic
GCF_001885655.1_ASM188565v1_genomic
GCF_002025445.1_KWv02_genomic
GCF_002095975.1_ASM209597v1_genomic
GCF_002933835.1_ASM293383v1_genomic
GCF_003019735.1_ASM301973v1_genomic
GCF_003112475.1_ASM311247v1_genomic
GCF_003206555.1_ASM320655v1_genomic
GCF_003206625.1_ASM320662v1_genomic
GCF_003730145.1_ASM373014v1_genomic
GCF_003945305.1_ASM394530v1_genomic
GCF_004305995.1_ASM430599v1_genomic
GCF_008257445.1_ASM825744v1_genomic
GCF_008579225.1_ASM857922v1_genomic
GCF_008579325.1_ASM857932v1_genomic
GCF_008579445.1_ASM857944v1_genomic
GCF_008579565.1_ASM857956v1_genomic
GCF_008579685.1_ASM857968v1_genomic
GCF_008757435.1_ASM875743v1_genomic
GCF_008974145.1_ASM897414v1_genomic
GCF_009792235.1_ASM979223v1_genomic
GCF_009811815.1_ASM981181v1_genomic
GCF_009811835.1_ASM981183v1_genomic
GCF_009811855.1_ASM981185v1_genomic
GCF_009811875.1_ASM981187v1_genomic
GCF_010196425.1_ASM1019642v1_genomic
GCF_014218745.1_ASM1421874v1_genomic
GCF_014218765.1_ASM1421876v1_genomic
GCF_014696875.1_ASM1469687v1_genomic
GCF_014698335.1_ASM1469833v1_genomic
GCF_014698375.1_ASM1469837v1_genomic
GCF_014698445.1_ASM1469844v1_genomic
GCF_015206885.1_ASM1520688v1_genomic
GCF_015207115.1_ASM1520711v1_genomic
GCF_015207885.1_ASM1520788v1_genomic
GCF_015207925.1_ASM1520792v1_genomic
Microcystis_aeruginosa_CS_1036_genomic
Microcystis_aeruginosa_CS_33801_genomic
Microcystis_aeruginosa_CS_55201
Microcystis_aeruginosa_CS_555A01A07
Microcystis_aeruginosa_CS_55603

**Microcystis_aeruginosa_CS_55801A06**
**Microcystis_aeruginosa_CS_56304**
**Microcystis_aeruginosa_CS_56401**
**Microcystis_aeruginosa_CS_56702**
**Microcystis_aeruginosa_CS_56702A1**
**Microcystis_aeruginosa_CS_573**
**Microcystis_aeruginosa_CS_579**
**Microcystis_aeruginosa_CS_583**
**Microcystis_sp_CS_574**
**GCA_001264245.1_ASM126424v1_genomic.fna**
**GCF_000021805.1_ASM2180v1_genomic**
**GCF_000024045.1_ASM2404v1_genomic**
**GCF_000473895.1_KS51_v1_genomic**
**GCF_000952155.1_asmbly001_genomic**
**GCA_023546805.1_ASM2354680v1_genomic.fna**
**GCF_000317125.1_ASM31712v1_genomic**
**GCF_002939305.1_ASM293930v1_genomic**
**GCF_003015105.1_ASM301510v1_genomic**
**GCF_003991895.1_ASM399189v1_genomic**
**GCF_012932785.1_ASM1293278v1_genomic**
**GCF_014696895.1_ASM1469689v1_genomic**
**GCF_018502385.1_ASM1850238v1_genomic**
**GCF_000011385.1_ASM1138v1_genomic**
**GCF_000484535.1_ASM48453v1_genomic**
**GCA_021018745.1_ASM2101874v1_genomic.fna**
**GCF_001870225.1_ASM187022v1_genomic**
**GCA_022206215.1_ASM2220621v1_genomic.fna**
**GCF_017591665.1_ASM1759166v1_genomic**
**GCF_016056305.1_ASM1605630v1_genomic**
**Anabaena_sp_CS_54202**
**GCF_000312705.1_ASM31270v1_genomic**
**GCF_000317695.1_ASM31769v1_genomic**
**GCF_000332135_1.scaf**
**GCF_001277295.1_ASM127729v1_genomic**
**GCF_001597745.1_ASM159774v1_genomic**
**GCF_001597855.1_ASM159785v1_genomic**
**GCF_002367975.1_ASM236797v1_genomic**
**GCF_002368015.1_ASM236801v1_genomic**
**GCF_009498015.1_ASM949801v1_genomic**
**GCF_009711975.1_ASM971197v1_genomic**
**GCF_009712035.1_ASM971203v1_genomic**
**GCF_009712085.1_ASM971208v1_genomic**
**GCF_014696225.1_ASM1469622v1_genomic**
**GCF_014696335.1_ASM1469633v1_genomic**
**GCF_014696465.1_ASM1469646v1_genomic**
**GCF_014696755.1_ASM1469675v1_genomic**
**GCF_014696765.1_ASM1469676v1_genomic**
**GCF_014696775.1_ASM1469677v1_genomic**

GCF_014696825.1_ASM1469682v1_genomic
GCF_014697105.1_ASM1469710v1_genomic
GCF_014697125.1_ASM1469712v1_genomic
GCF_014697155.1_ASM1469715v1_genomic
GCF_014697335.1_ASM1469733v1_genomic
GCF_014697485.1_ASM1469748v1_genomic
GCF_014697625.1_ASM1469762v1_genomic
GCF_014698305.1_ASM1469830v1_genomic
GCF_014698355.1_ASM1469835v1_genomic
GCF_014698735.1_ASM1469873v1_genomic
GCF_015207835.1_ASM1520783v1_genomic
GCF_015245355.1_ASM1524535v1_genomic
Anabaenopsis_arnoldii_genomic
Aphanizomenon_sp_CS_73332
GCA_017346855.1_ASM1734685v1_genomic.fna
GCA_017346875.1_ASM1734687v1_genomic.fna
GCF_000521175_1.scaf
GCF_000789435.1_ASM78943v1_genomic
GCF_009712065.1_ASM971206v1_genomic
GCF_014696815.1_ASM1469681v1_genomic
GCF_014697315.1_ASM1469731v1_genomic
GCF_014698245.1_ASM1469824v1_genomic
GCF_014698265.1_ASM1469826v1_genomic
GCF_014698295.1_ASM1469829v1_genomic
GCF_014698695.1_ASM1469869v1_genomic
GCF_014698705.1_ASM1469870v1_genomic
GCF_014698725.1_ASM1469872v1_genomic
GCF_014698755.1_ASM1469875v1_genomic
GCF_016056295.1_ASM1605629v1_genomic
GCF_002368055.1_ASM236805v1_genomic
GCF_014696595.1_ASM1469659v1_genomic
GCF_014698045.1_ASM1469804v1_genomic
GCF_006968745.1_ASM696874v1_genomic
GCF_012912105.1_ASM1291210v1_genomic
GCF_012912125.1_ASM1291212v1_genomic
GCF_012912135.1_ASM1291213v1_genomic
GCF_017982655.1_ASM1798265v1_genomic
GCA_000317435.1_ASM31743v1_genomic.fna
GCA_000734895.2_ASM73489v2_genomic.fna
GCA_019977735.1_ASM1997773v1_genomic.fna
GCF_000316575.1_ASM31657v1_genomic
GCF_000317435.1_ASM31743v1_genomic
GCF_000331305.scaf
GCF_000734895.2_ASM73489v2_genomic
GCF_001904745.1_ASM190474v1_genomic
GCF_002367995.1_ASM236799v1_genomic
GCF_002368095.1_ASM236809v1_genomic
GCF_002368175.1_ASM236817v1_genomic

GCF_002368195.1_ASM236819v1_genomic
GCF_002368375.1_ASM236837v1_genomic
GCF_002368395.1_ASM236839v1_genomic
GCF_002368415.1_ASM236841v1_genomic
GCF_002368455.1_ASM236845v1_genomic
GCF_007830875.1_ASM783087v1_genomic
GCF_014696315.1_ASM1469631v1_genomic
GCF_014696435.1_ASM1469643v1_genomic
GCF_014696495.1_ASM1469649v1_genomic
GCF_014696555.1_ASM1469655v1_genomic
GCF_014696565.1_ASM1469656v1_genomic
GCF_014697055.1_ASM1469705v1_genomic
GCF_900185595.1_CalSC01_2013_genomic
GCF_000317285.1_ChlPCC6912_1.0_genomic
GCF_015207795.1_ASM1520779v1_genomic
GCA_021650815.1_ASM2165081v1_genomic.fna
GCF_000175835.1_ASM17583v1_genomic
GCF_001432185.1_ASM143218v1_genomic
GCF_001586755.1_ASM158675v1_genomic
GCF_001858115.1_ASM185811v1_genomic
GCF_001858125.1_ASM185812v1_genomic
GCF_002027345.1_ASM202734v1_genomic
GCF_002114155.1_ASM211415v1_genomic
GCF_002321945.1_ASM232194v1_genomic
GCF_002893125.1_ASM289312v1_genomic
GCF_002893145.1_ASM289314v1_genomic
GCF_002893155.1_ASM289315v1_genomic
GCF_002893185.1_ASM289318v1_genomic
GCF_002893205.1_ASM289320v1_genomic
GCF_002893215.1_ASM289321v1_genomic
GCF_002893245.1_ASM289324v1_genomic
GCF_002893265.1_ASM289326v1_genomic
GCF_002893285.1_ASM289328v1_genomic
GCF_003367075.1_ASM336707v1_genomic
GCF_006523545.1_ASM652354v1_genomic
GCF_012583295.1_ASM1258329v1_genomic
GCF_012583345.1_ASM1258334v1_genomic
GCF_014489415.1_ASM1448941v1_genomic
GCF_018139025.1_ASM1813902v1_genomic
GCA_003994795.1_ASM399479v1_genomic
GCF_000317535.1_ASM31753v1_genomic
GCF_014697195.1_ASM1469719v1_genomic
GCF_016056275.1_ASM1605627v1_genomic
Dolichospermum_circinale_CS_1031_genomic
Dolichospermum_circinale_CS_1225_genomic
Dolichospermum_circinale_CS_53405_genomic
Dolichospermum_circinale_CS_53701_genomic
Dolichospermum_circinale_CS_53703_genomic

**Dolichospermum_circinale_CS_53711**
**Dolichospermum_circinale_CS_539**
**Dolichospermum_circinale_CS_53909**
**Dolichospermum_circinale_CS_54104**
**Dolichospermum_circinale_CS_54106**
**Dolichospermum_circinale_CS_547**
**Dolichospermum_lemmermannii_CS_548**
**Dolichospermum_planctonicum_CS_1226_genomic**
**GCA_017346795.1_ASM1734679v1_genomic.fna**
**GCA_017346815.1_ASM1734681v1_genomic.fna**
**GCA_017346835.1_ASM1734683v1_genomic.fna**
**GCA_017355425.1_ASM1735542v1_genomic.fna**
**GCA_024584745.1_ASM2458474v1_genomic.fna**
**GCF_000426905_1.scaf**
**GCF_000426925_1.scaf**
**GCF_002368115.1_ASM236811v1_genomic**
**GCF_009712125.1_ASM971212v1_genomic**
**GCF_012516395.1_ASM1251639v1_genomic**
**GCF_015207785.1_ASM1520778v1_genomic**
**GCF_005402965.1_ASM540296v1_genomic**
**GCF_008121535.1_ASM812153v1_genomic**
**GCF_009711925.1_ASM971192v1_genomic**
**GCF_009711935.1_ASM971193v1_genomic**
**GCF_009711965.1_ASM971196v1_genomic**
**GCF_009711985.1_ASM971198v1_genomic**
**GCF_009712025.1_ASM971202v1_genomic**
**GCF_009712075.1_ASM971207v1_genomic**
**GCF_014697585.1_ASM1469758v1_genomic**
**GCF_015207875.1_ASM1520787v1_genomic**
**GCA_025999015.1_ASM2599901v1_genomic.fna**
**GCF_000231365.1_ASM23136v1_genomic**
**GCF_000315585_1.scaf**
**GCF_000317205.1_FisPCC7414_1.0_genomic**
**GCF_000317225.1_FisPCC7521_1.0_genomic**
**GCF_000317245.1_FisPCC73103_1.0_genomic**
**GCF_000447295_1.scaf**
**GCF_000517105_1.scaf**
**GCF_001548455.1_ASM154845v1_genomic**
**GCF_001904645.1_ASM190464v1_genomic**
**GCF_002368315.1_ASM236831v1_genomic**
**GCF_002870185.1_ASM287018v1_genomic**
**GCF_002870195.1_ASM287019v1_genomic**
**GCF_002870205.1_ASM287020v1_genomic**
**GCF_002870255.1_ASM287025v1_genomic**
**GCF_002870265.1_ASM287026v1_genomic**
**GCF_002870305.1_ASM287030v1_genomic**
**GCF_002870325.1_ASM287032v1_genomic**
**GCF_002870345.1_ASM287034v1_genomic**

GCF_002870365.1_ASM287036v1_genomic
GCF_002870385.1_ASM287038v1_genomic
GCF_002870405.1_ASM287040v1_genomic
GCF_002870415.1_ASM287041v1_genomic
GCF_002870445.1_ASM287044v1_genomic
GCF_002870465.1_ASM287046v1_genomic
GCF_002870475.1_ASM287047v1_genomic
GCF_002870505.1_ASM287050v1_genomic
GCF_002870525.1_ASM287052v1_genomic
GCF_002870545.1_ASM287054v1_genomic
GCF_002870565.1_ASM287056v1_genomic
GCF_002870705.1_ASM287070v1_genomic
GCF_002870755.1_ASM287075v1_genomic
GCF_003346965.1_ASM334696v1_genomic
GCF_014697535.1_ASM1469753v1_genomic
GCF_000332295_1.scaf
GCF_015206815.1_ASM1520681v1_genomic
GCF_002368275.1_ASM236827v1_genomic
GCA_029953635.1_ASM2995363v1_genomic.fna
GCF_001275395.1_ASM127539v1_genomic
GCF_000817785.2_ASM81778v2_genomic
GCF_014698965.1_ASM1469896v1_genomic
GCF_014831675.1_ASM1483167v1_genomic
GCF_000315565.scaf
GCF_001456025.1_ASM145602v1_genomic
GCA_022376295.1_ASM2237629v1_genomic.fna
GCF_000340565.2_ASM34056v3_genomic
GCF_001623485.1_ASM162348v1_genomic
GCF_002218065.1_ASM221806v1_genomic
GCF_003054475.1_ASM305447v1_genomic
GCF_015207755.1_ASM1520775v1_genomic
Nodularia_sphaerocarpa_CS_585
Nodularia_spumigena_CS_1038_genomic
Nodularia_spumigena_CS_33602_genomic
Nodularia_spumigena_CS_584
Nodularia_spumigena_CS_58605
Nodularia_spumigena_CS_58801
Nodularia_spumigena_CS_58802A
Nodularia_spumigena_CS_58802A10
Nodularia_spumigena_CS_58802B
Nodularia_spumigena_CS_58805
Nodularia_spumigena_CS_58806
Nodularia_spumigena_CS_58907
Nodularia_spumigena_CS_59001
Nodularia_spumigena_CS_59001A
Nodularia_spumigena_CS_59104
Nodularia_spumigena_CS_59107A
Nodularia_spumigena_CS_59112

GCA_003990685.1_ASM399068v1_genomic
GCA_003990705.1_ASM399070v1_genomic
GCA_013393905.1_ASM1339390v1_genomic.fna
GCA_019976755.1_ASM1997675v1_genomic.fna
GCA_022063185.1_ASM2206318v1_genomic.fna
GCA_023734775.1_ASM2373477v1_genomic.fna
GCA_028623165.1_ASM2862316v1_genomic.fna
GCF_000009705.1_ASM970v1_genomic
GCF_000020025.1_ASM2002v1_genomic
GCF_000196515.1_ASM19651v1_genomic
GCF_000316625.1_ASM31662v1_genomic
GCF_000316645.1_ASM31664v1_genomic
GCF_001298445.1_ASM129844v1_genomic
GCF_001548375.1_ASM154837v1_genomic
GCF_001712795_1.scaf
GCF_001904715.1_ASM190471v1_genomic
GCF_002154695.1_ASM215469v1_genomic
GCF_002154725.1_ASM215472v1_genomic
GCF_002155185.1_ASM215518v1_genomic
GCF_002245975.1_ASM224597v1_genomic
GCF_002245985.1_ASM224598v1_genomic
GCF_002246015.1_ASM224601v1_genomic
GCF_002368035.1_ASM236803v1_genomic
GCF_002368155.1_ASM236815v1_genomic
GCF_002368215.1_ASM236821v1_genomic
GCF_002368335.1_ASM236833v1_genomic
GCF_002607925.1_ASM260792v1_genomic
GCF_002607955.1_ASM260795v1_genomic
GCF_002607965.1_ASM260796v1_genomic
GCF_002608015.1_ASM260801v1_genomic
GCF_002608075.1_ASM260807v1_genomic
GCF_002608105.1_ASM260810v1_genomic
GCF_002608135.1_ASM260813v1_genomic
GCF_002608145.1_ASM260814v1_genomic
GCF_002608225.1_ASM260822v1_genomic
GCF_002608345.1_ASM260834v1_genomic
GCF_002813575.1_ASM281357v1_genomic
GCF_002896875.1_ASM289687v1_genomic
GCF_002897135_1.scaf
GCF_002949735.1_ASM294973v1_genomic
GCF_002949795.1_ASM294979v1_genomic
GCF_003113895.1_ASM311389v1_genomic
GCF_003326245.1_ASM332624v1_genomic
GCF_003443655.1_ASM344365v1_genomic
GCF_005869855.1_ASM586985v1_genomic
GCF_009372195.1_ASM937219v1_genomic
GCF_010091925.1_ASM1009192v1_genomic
GCF_013343235.1_ASM1334323v1_genomic

GCF_013393925.1_ASM1339392v1_genomic
GCF_013393945.1_ASM1339394v1_genomic
GCF_014023275.1_ASM1402327v1_genomic
GCF_014696615.1_ASM1469661v1_genomic
GCF_014696625.1_ASM1469662v1_genomic
GCF_014696655.1_ASM1469665v1_genomic
GCF_014696905.1_ASM1469690v1_genomic
GCF_014697355.1_ASM1469735v1_genomic
GCF_014697425.1_ASM1469742v1_genomic
GCF_014697455.1_ASM1469745v1_genomic
GCF_014697475.1_ASM1469747v1_genomic
GCF_014697495.1_ASM1469749v1_genomic
GCF_014697615.1_ASM1469761v1_genomic
GCF_014698035.1_ASM1469803v1_genomic
GCF_014698055.1_ASM1469805v1_genomic
GCF_014698115.1_ASM1469811v1_genomic
GCF_014698475.1_ASM1469847v1_genomic
GCF_014698495.1_ASM1469849v1_genomic
GCF_014698505.1_ASM1469850v1_genomic
GCF_014698525.1_ASM1469852v1_genomic
GCF_014698635.1_ASM1469863v1_genomic
GCF_014698795.1_ASM1469879v1_genomic
GCF_014698835.1_ASM1469883v1_genomic
GCF_014904695.1_ASM1490469v1_genomic
GCF_015206895.1_ASM1520689v1_genomic
GCF_015206985.1_ASM1520698v1_genomic
GCF_015207705.1_ASM1520770v1_genomic
GCF_015714705.1_ASM1571470v1_genomic
GCF_015714725.1_ASM1571472v1_genomic
GCF_017164015.1_ASM1716401v1_genomic
GCA_000175855.1_ASM17585v1_genomic
GCF_000350105.1_ASM35010v1_genomic
GCF_014698825.1_ASM1469882v1_genomic
GCF_000316665.1_ASM31666v1_genomic
GCF_015206995.1_ASM1520699v1_genomic
GCF_000346485_2.scaf
GCF_000817735.3_ASM81773v3_genomic
GCF_000828085.3_ASM82808v3_genomic
GCF_014698435.1_ASM1469843v1_genomic
GCF_001586785.1_ASM158678v2_genomic
GCF_002368075.1_ASM236807v1_genomic
GCF_014299995.1_ASM1429999v1_genomic
GCF_014696195.1_ASM1469619v1_genomic
GCF_014696235.1_ASM1469623v1_genomic
GCF_015207205.1_ASM1520720v1_genomic
Sphaerospermopsis_kisseleviana_CS_549
GCF_000300115_1.scaf
GCF_000760695.4_ASM76069v4_genomic

GCF_000828075.3_ASM82807v3_genomic
GCF_001858025.1_ASM185802v1_genomic
GCF_002218085.1_ASM221808v1_genomic
GCF_002368295.1_ASM236829v1_genomic
GCF_011769525.1_ASM1176952v1_genomic
GCF_014697075.1_ASM1469707v1_genomic
GCF_000204075.1_ASM20407v1_genomic
GCF_003991935.1_ASM399193v1_genomic
GCF_009856605.1_ASM985660v1_genomic
GCF_014222135_1.scaf
GCF_014222145_1.scaf
GCF_014222155_1.scaf
GCF_014222225_1.scaf
GCF_014222245_1.scaf
GCF_014697165.1_ASM1469716v1_genomic
GCA_002163975.1_ASM216397v1_genomic.fna
GCF_004323185.1_ASM432318v1_genomic
GCA_000317515.1_ASM31751v1_genomic.fna
GCA_000175415.3_ASM17541v3_genomic
GCA_003060805.1_ASM306080v1_genomic.fna
GCA_025200965.1_ASM2520096v1_genomic.fna
GCF_000210375.1_ASM21037v1_genomic
GCF_000307915_1.chr
GCF_001611905.1_ASM161190v1_genomic
GCF_009176225.1_ASM917622v1_genomic
GCF_014698385.1_ASM1469838v1_genomic
GCF_014698675.1_ASM1469867v1_genomic
GCF_014698815.1_ASM1469881v1_genomic
GCF_016745315.1_ASM1674531v1_genomic
GCF_000332355_1.scaf
GCF_000155555_1.scaf
GCF_014695285.1_ASM1469528v1_genomic
GCF_014695375.1_ASM1469537v1_genomic
GCF_014695435.1_ASM1469543v1_genomic
GCF_014695475.1_ASM1469547v1_genomic
GCF_014695485.1_ASM1469548v1_genomic
GCF_014695685.1_ASM1469568v1_genomic
GCF_014695825.1_ASM1469582v1_genomic
GCF_014698185.1_ASM1469818v1_genomic
GCF_014698845.1_ASM1469884v1_genomic
GCF_015207375.1_ASM1520737v1_genomic
GCF_015207445.1_ASM1520744v1_genomic
GCF_000317495.1_ASM31749v1_genomic
GCF_000022045.1_ASM2204v1_genomic
GCF_003013815.1_ASM301381v1_genomic
GCF_001746915.1_ASM174691v1_genomic
GCF_014696935.1_ASM1469693v1_genomic
GCF_014696985.1_ASM1469698v1_genomic

GCF_014697015.1_ASM1469701v1_genomic
GCF_000317045.1_ASM31704v1_genomic
GCF_001870905.1_ASM187090v1_genomic
GCF_000180455.1_ASM18045v1_genomic
GCF_000332155_1.scaf
GCA_016446395.1_ASM1644639v1_genomic.fna
GCF_000972705.2_ASM97270v2_genomic
GCA_014489865.1_ASM1448986v1_genomic
GCF_000173555.1_ASM17355v1_genomic
GCF_000973065.1_ASM97306v1_genomic
GCA_012516315.1_ASM1251631v1_genomic.fna
GCF_000169095.1_ASM16909v1_genomic
GCF_000478195.2_ASM47819v2_genomic
GCF_000817775.2_ASM81777v2_genomic
GCA_022701255.1_ASM2270125v1_genomic.fna
GCA_022701275.1_ASM2270127v1_genomic.fna
GCF_000214075.1_ASM21407v1_genomic
GCF_000317515.1_ASM31751v1_genomic
GCF_009846485_1.scaf
GCF_013179805.1_ASM1317980v1_genomic
GCF_014695175.1_ASM1469517v1_genomic
GCF_014695295.1_ASM1469529v1_genomic
GCF_014695525.1_ASM1469552v1_genomic
GCF_014695585.1_ASM1469558v1_genomic
GCF_014695715.1_ASM1469571v1_genomic
GCF_014695725.1_ASM1469572v1_genomic
GCF_014695895.1_ASM1469589v1_genomic
GCF_014695925.1_ASM1469592v1_genomic
GCF_014696105.1_ASM1469610v1_genomic
GCF_014696155.1_ASM1469615v1_genomic
GCF_014849525.1_ASM1484952v1_genomic
GCF_015207455.1_ASM1520745v1_genomic
GCF_000211815_1.scaf
GCF_001767235.1_ASM176723v1_genomic
GCF_001854205.1_ASM185420v1_genomic
GCF_001942495.1_ASM194249v1_genomic
GCA_001767235.1_ASM176723v1_genomic.fna
GCA_001854205.2_ASM185420v2_genomic.fna
GCF_003838225.1_ASM383822v1_genomic
GCF_014132355.1_ASM1413235v1_genomic
GCF_000317105.1_ASM31710v1_genomic
GCF_000317475.1_ASM31747v1_genomic
GCF_000332335_1.scaf
GCF_014697545.1_ASM1469754v1_genomic
GCF_014698145.1_ASM1469814v1_genomic
Oscillatoria_sp_CS_180_genomic
GCF_012295525.1_ASM1229552v1_genomic
GCF_016632315.1_ASM1663231v1_genomic

GCA_020386575.1_ASM2038657v1_genomic.fna
GCA_023983615.1_ASM2398361v1_genomic.fna
GCF_001637315.1_ASM163731v1_genomic
GCF_001904725.1_ASM190472v1_genomic
GCF_001904775.1_ASM190477v1_genomic
GCF_014695595.1_ASM1469559v1_genomic
GCF_014695795.1_ASM1469579v1_genomic
GCF_014695965.1_ASM1469596v1_genomic
GCF_014696675.1_ASM1469667v1_genomic
GCF_014697375.1_ASM1469737v1_genomic
GCF_014764505.1_ASM1476450v1_genomic
GCF_015207735.1_ASM1520773v1_genomic
GCF_016807185_1.scaf
GCF_001276715.1_ASM127671v1_genomic
GCF_014698175.1_ASM1469817v1_genomic
GCF_014698535.1_ASM1469853v1_genomic
GCA_003609755.1_ASM360975v1_genomic
GCA_904830775.1_P. agardhii_No.2A_genomic.fna
GCA_904830845.1_PA_No.365_genomic.fna
GCA_904830855.1_PA_No.66_genomic.fna
GCA_904830915.1_P. agardhii_PCC7805_genomic.fna
GCA_904830925.1_P. pseudagardhii_No.713_genomic.fna
GCA_904830935.1_P. agardhii_No.976_genomic.fna
GCF_000464665_1.scaf
GCF_000464745_1.scaf
GCF_000464765_1.scaf
GCF_000464785_1.scaf
GCF_000464805_1.scaf
GCF_000464825_1.scaf
GCF_000464845_1.scaf
GCF_000710505.chr
GCF_005402765.1_ASM540276v2_genomic
GCF_014696265.1_ASM1469626v1_genomic
GCF_014697575.1_ASM1469757v1_genomic
GCF_015207485.1_ASM1520748v1_genomic
GCF_900009135_1.scaf
GCF_900009145_1.scaf
GCF_900009265_2.scaf
GCF_900009275.2chr
GCF_900010725_2.scaf
GCF_903969095_1.scaf
GCF_015207665.1_ASM1520766v1_genomic
GCF_000014265.1_ASM1426v1_genomic
GCF_002412335.2_ASM241233v2_genomic
GCF_015207285.1_ASM1520728v1_genomic
GCF_015207295.1_ASM1520729v1_genomic
GCF_015207345.1_ASM1520734v1_genomic
GCF_015207505.1_ASM1520750v1_genomic

GCF_001904635.1_ASM190463v1_genomic
GCF_000756305.1_ASM75630v1_genomic
GCF_000317025.1_ASM31702v1_genomic
GCF_000332195_1.scaf
GCF_003003995.1_ASM300399v1_genomic
GCA_000317575.1_ASM31757v1_genomic.fna
GCA_002355455.1_ASM235545v1_genomic.fna
GCF_000332055.1_ASM33205v1_genomic
GCF_014697025.1_ASM1469702v1_genomic
GCF_002075285.3_ASM207528v3_genomic
GCF_004212215.1_ASM421221v1_genomic
GCF_016888165.1_ASM1688816v1_genomic
GCA_020091585.1_ASM2009158v1_genomic.fna
GCA_022372495.1_ASM2237249v1_genomic.fna
GCA_026802175.1_ASM2680217v1_genomic.fna
GCF_000316115.scaf
GCF_000316605.1_ASM31660v1_genomic
GCF_000332095_2.scaf
GCF_000353285_1.scaf
GCF_000482245.1_LepHIscaffolds_genomic
GCF_000763385.1_ASM76338v1_genomic
GCF_001485215.1_ASM148521v1_genomic
GCF_001548395.1_ASM154839v1_genomic
GCF_001548435.1_ASM154843v1_genomic
GCF_001631935.1_ASM163193v1_genomic
GCF_001637395.1_ASM163739v1_genomic
GCF_001939115.1_ASM193911v1_genomic
GCF_002142475.1_ASM214247v1_genomic
GCF_002142495.1_ASM214249v1_genomic
GCF_002215035.1_ASM221503v1_genomic
GCF_002286735.1_ASM228673v1_genomic
GCF_002368255.1_ASM236825v1_genomic
GCF_014695405.1_ASM1469540v1_genomic
GCF_014695575.1_ASM1469557v1_genomic
GCF_014695745.1_ASM1469574v1_genomic
GCF_014695775.1_ASM1469577v1_genomic
GCF_014695875.1_ASM1469587v1_genomic
GCF_014695975.1_ASM1469597v1_genomic
GCF_014696055.1_ASM1469605v1_genomic
GCF_014696065.1_ASM1469606v1_genomic
GCF_014696095.1_ASM1469609v1_genomic
GCF_014697225.1_ASM1469722v1_genomic
GCF_014697245.1_ASM1469724v1_genomic
GCF_014697265.1_ASM1469726v1_genomic
GCF_014697285.1_ASM1469728v1_genomic
GCF_016403105.1_ASM1640310v1_genomic
GCF_016888135.1_ASM1688813v1_genomic
GCF_018128485.1_ASM1812848v1_genomic

GCA_019891175.1_ASM1989117v1_genomic.fna
GCA_021379005.1_ASM2137900v1_genomic.fna
GCF_018739865.1_ASM1873986v1_genomic
GCF_018739885.1_ASM1873988v1_genomic
GCF_001904615.1_ASM190461v1_genomic
GCF_002742025.1_ASM274202v1_genomic
GCF_014696455.1_ASM1469645v1_genomic
GCF_014696485.1_ASM1469648v1_genomic
GCF_014698135.1_ASM1469813v1_genomic
GCF_014698215.1_ASM1469821v1_genomic
GCF_014698235.1_ASM1469823v1_genomic
GCF_015207425.1_ASM1520742v1_genomic
GCF_010091945.1_ASM1009194v1_genomic
GCF_000775285.1_ASM77528v1_genomic
GCF_000309385_1.scaf
GCF_012911975.1_ASM1291197v1_genomic
GCF_014695395.1_ASM1469539v1_genomic
GCF_014696135.1_ASM1469613v1_genomic
GCF_014696165.1_ASM1469616v1_genomic
GCF_015207395.1_ASM1520739v1_genomic
GCF_015207525.1_ASM1520752v1_genomic
GCF_014696015.1_ASM1469601v1_genomic
GCF_015207605.1_ASM1520760v1_genomic
GCF_001650195.1_ASM165019v1_genomic
GCF_003003695.1_ASM300369v1_genomic
GCF_000332315_1.scaf
GCA_027942295.1_ASM2794229v1_genomic.fna
GCA_029910235.1_ASM2991023v1_genomic.fna
GCF_000317065.1_ASM31706v1_genomic
GCF_000332175_1.scaf
GCF_002251945.1_ASM225194v1_genomic
GCF_002914585.1_ASM291458v1_genomic
GCF_003967015.1_ASM396701v1_genomic
GCF_006861605.1_ASM686160v1_genomic
GCF_006937785.1_ASM693778v1_genomic
GCF_012863495.1_ASM1286349v1_genomic
GCF_013361095_1.scaf
GCF_014696345.1_ASM1469634v1_genomic
GCF_014696355.1_ASM1469635v1_genomic
GCF_014696385.1_ASM1469638v1_genomic
GCF_014696715.1_ASM1469671v1_genomic
GCF_014696925.1_ASM1469692v1_genomic
GCF_016584665.1_ASM1658466v1_genomic
GCF_015207255.1_ASM1520725v1_genomic
GCA_013177315.1_ASM1317731v1_genomic.fna
GCA_023650955.1_ASM2365095v1_genomic.fna
GCA_025999095.1_ASM2599909v1_genomic.fna
GCF_000011345.1_ASM1134v1_genomic

GCF_000505665.1_ASM50566v1_genomic
GCF_003555505.1_ASM355550v2_genomic
GCF_003990665.1_ASM399066v2_genomic
GCF_008386235.1_ASM838623v1_genomic
GCF_017086385.1_ASM1708638v1_genomic
GCF_000314005_1.scaf
GCF_001890765_1.scaf
GCF_016888125.1_ASM1688812v1_genomic
Spirulina_major_CS_329
Spirulina_sp_CS_78501
Spirulina_subsalsa_CS_330
GCA_018336915.1_ASM1833691v1_genomic.fna
GCA_021497025.1_ASM2149702v1_genomic.fna
GCA_024347835.1_ASM2434783v1_genomic.fna
GCF_000018105.1_ASM1810v1_genomic
GCF_003231495.1_AcaThom1.0_genomic
GCF_018336915.1chr
GCF_000817745.2_ASM81774v2_genomic
GCF_000025125.1_ASM2512v1_genomic
GCF_000586015.1_SynSpo.0_genomic
GCF_002018015.1_ASM201801v1_genomic
GCF_900047545.1_m9_genomic
GCA_018141785.1_ASM1814178v1_genomic.fna
GCF_000317145.1_ASM31714v1_genomic
GCA_018399015.1_ASM1839901v1_genomic.fna
GCF_000155635_1.scaf
GCF_000316515.1_ASM31651v1_genomic
GCF_003011885.1_ASM301188v1_genomic
GCF_009834675.1_ASM983467v1_genomic
GCF_014280235.1_ASM1428023v1_genomic
GCF_014697415.1_ASM1469741v1_genomic
GCF_015207535.1_ASM1520753v1_genomic
GCF_015207615.1_ASM1520761v1_genomic
GCF_900088535.1_ASM90008853v1_genomic
GCF_000317615.1_ASM31761v1_genomic
GCF_003003775.1_ASM300377v1_genomic
GCA_001180325.1_CLC_assembled_contigs_genomic
GCF_000007925.1_ASM792v1_genomic
GCF_000011465.1_ASM1146v1_genomic
GCF_000011485.1_ASM1148v1_genomic
GCF_000012465.1_ASM1246v1_genomic
GCF_000012505.1_ASM1250v1_genomic
GCF_000012645.1_ASM1264v1_genomic
GCF_000015645.1_ASM1564v1_genomic
GCF_000015665.1_ASM1566v1_genomic
GCF_000015685.1_ASM1568v1_genomic
GCF_000015965.1_ASM1596v1_genomic
GCF_000018065.1_ASM1806v1_genomic

GCF_000018585.1_ASM1858v1_genomic
GCF_000158595_1.scaf
GCF_000757845.1_ASM75784v1_genomic
GCF_000757865.1_ASM75786v1_genomic
GCF_000759855.1_ASM75985v1_genomic
GCF_000759865.1_ASM75986v1_genomic
GCF_000759875.1_ASM75987v1_genomic
GCF_000759885.1_ASM75988v1_genomic
GCF_000759935.1_ASM75993v1_genomic
GCF_000759955.1_ASM75995v1_genomic
GCF_000759975.1_ASM75997v1_genomic
GCF_000760015.1_ASM76001v1_genomic
GCF_000760035.1_ASM76003v1_genomic
GCF_000760055.1_ASM76005v1_genomic
GCF_000760075.1_ASM76007v1_genomic
GCF_000760095.1_ASM76009v1_genomic
GCF_000760115.1_ASM76011v1_genomic
GCF_000760155.1_ASM76015v1_genomic
GCF_000760175.1_ASM76017v1_genomic
GCF_000760195.1_ASM76019v1_genomic
GCF_000760215.1_ASM76021v1_genomic
GCF_000760235.1_ASM76023v1_genomic
GCF_000760255.1_ASM76025v1_genomic
GCF_000760275.1_ASM76027v1_genomic
GCF_000760295.1_ASM76029v1_genomic
GCF_000760315.1_ASM76031v1_genomic
GCF_000760335.1_ASM76033v1_genomic
GCF_000760355.1_ASM76035v1_genomic
GCF_000760375.1_ASM76037v1_genomic
GCF_001180245.1_CLC_assembled_contigs_genomic
GCF_001180265.1_CLC_assembled_contigs_genomic
GCF_001180285.1_CLC_assembled_contigs_genomic
GCF_001180305.1_CLC_assembled_contigs_genomic
GCF_001180325.1_CLC_assembled_contigs_genomic
GCF_001631965.1_ASM163196v1_genomic
GCF_001631985.1_ASM163198v1_genomic
GCF_001632025.1_ASM163202v1_genomic
GCF_001632075.1_ASM163207v1_genomic
GCF_001989415.1_ASM198941v1_genomic
GCF_001989435.1_ASM198943v1_genomic
GCF_001989455.1_ASM198945v1_genomic
GCF_002025865.1_ASM202586v1_genomic
GCF_002025905.1_ASM202590v1_genomic
GCF_002025945.1_ASM202594v1_genomic
GCF_002025965.1_ASM202596v1_genomic
GCF_002025975.1_ASM202597v1_genomic
GCF_002026005.1_ASM202600v1_genomic
GCF_002026015.1_ASM202601v1_genomic

GCF_002026045.1_ASM202604v1_genomic
GCF_002026065.1_ASM202606v1_genomic
GCF_002026085.1_ASM202608v1_genomic
GCF_002026095.1_ASM202609v1_genomic
GCF_002026105.1_ASM202610v1_genomic
GCF_002026145.1_ASM202614v1_genomic
GCF_002026165.1_ASM202616v1_genomic
GCF_002026175.1_ASM202617v1_genomic
GCF_002026185.1_ASM202618v1_genomic
GCF_002026205.1_ASM202620v1_genomic
GCF_002026245.1_ASM202624v1_genomic
GCF_002156565.1_ASM215656v1_genomic
GCF_002812945.1_ASM281294v1_genomic
GCF_003208055.1_ASM320805v1_genomic
GCF_003208065.1_ASM320806v1_genomic
GCF_012933585.1_ASM1293358v1_genomic
GCF_012933595.1_ASM1293359v1_genomic
GCF_012933605.1_ASM1293360v1_genomic
GCF_013393905.1_ASM1339390v1_genomic
GCF_017695955.1_ASM1769595v1_genomic
GCF_017695965.1_ASM1769596v1_genomic
GCF_017695975.1_ASM1769597v1_genomic
GCF_017696015.1_ASM1769601v1_genomic
GCF_017696035.1_ASM1769603v1_genomic
GCF_017696055.1_ASM1769605v1_genomic
GCF_017696075.1_ASM1769607v1_genomic
GCF_017696085.1_ASM1769608v1_genomic
GCF_017696175.1_ASM1769617v1_genomic
GCF_017696205.1_ASM1769620v1_genomic
GCF_017696245.1_ASM1769624v1_genomic
GCF_017696275.1_ASM1769627v1_genomic
GCF_017696315.1_ASM1769631v1_genomic
GCA_000063505.1_ASM6350v1_genomic.fna
GCA_000063525.1_ASM6352v1_genomic.fna
GCA_000284135.1_ASM28413v1_genomic.fna
GCA_000284215.1_ASM28421v1_genomic.fna
GCA_000284455.1_ASM28445v1_genomic.fna
GCA_000817325.1_ASM81732v1_genomic.fna
GCA_003015225.1_ASM301522v1_genomic.fna
GCA_018282115.1_ASM1828211v1_genomic.fna
GCA_019924095.1_ASM1992409v1_genomic.fna
GCA_023078835.1_ASM2307883v1_genomic.fna
GCA_023115425.1_ASM2311542v1_genomic.fna
GCF_000010065.1_ASM1006v1_genomic
GCF_000012525.1_ASM1252v1_genomic
GCF_000012625.1_ASM1262v1_genomic
GCF_000013205.1_ASM1320v1_genomic
GCF_000013225.1_ASM1322v1_genomic

GCF_000014585.1_ASM1458v1_genomic
GCF_000019485.1_ASM1948v1_genomic
GCF_000153045_1.scaf
GCF_000153065_1.scaf
GCF_000153805_1.scaf
GCF_000153825_1.scaf
GCF_000155595_1.scaf
GCF_000161795_1.scaf
GCF_000179235.2_ASM17923v2_genomic
GCF_000179255.1_ASM17925v1_genomic
GCF_000195975.1_ASM19597v1_genomic
GCF_000230675.1_ASM23067v1_genomic
GCF_000316685.1_ASM31668v1_genomic
GCF_000317085.1_ASM31708v1_genomic
GCF_000332275_1.chr
GCF_000485815.1_ASM48581v1_genomic
GCF_000515235_1.scaf
GCF_000715475.1_ASM71547v1_genomic
GCF_000737535.1_ASM73753v1_genomic
GCF_000737575.1_ASM73757v1_genomic
GCF_000737595.1_ASM73759v1_genomic
GCF_000817325.1_ASM81732v1_genomic
GCF_001039265.1_ASM103926v1_genomic
GCF_001040845.1_ASM104084v1_genomic
GCF_001182765.1_WH8103.1_genomic
GCF_001521855.1_ASM152185v1_genomic
GCF_001632105.1_ASM163210v1_genomic
GCF_001632165.1_ASM163216v1_genomic
GCF_001693255.1_ASM169325v1_genomic
GCF_001693275.1_ASM169327v1_genomic
GCF_001693295.1_ASM169329v1_genomic
GCF_001885215.1_ASM188521v1_genomic
GCF_002252625.1_ASM225262v1_genomic
GCF_002252635.1_ASM225263v1_genomic
GCF_002252665.1_ASM225266v1_genomic
GCF_002252675.1_ASM225267v1_genomic
GCF_002356215.1_ASM235621v1_genomic
GCF_002721235.1_ASM272123v1_genomic
GCF_002754935.1_ASM275493v1_genomic
GCF_002760345.1_ASM276034v1_genomic
GCF_002760375.1_ASM276037v1_genomic
GCF_002760395.1_ASM276039v1_genomic
GCF_002760415.1_ASM276041v1_genomic
GCF_002760445.1_ASM276044v1_genomic
GCF_002760475.1_ASM276047v1_genomic
GCF_003011125.1_ASM301112v1_genomic
GCF_003846445.1_ASM384644v1_genomic
GCF_003957805.1_ASM395780v1_genomic

GCF_004332405.1_ASM433240v1_genomic
GCF_004332415.1_ASM433241v1_genomic
GCF_005577135.1_ASM557713v1_genomic
GCF_007828135.1_ASM782813v1_genomic
GCF_008807075.1_ASM880707v1_genomic
GCF_009498715.1_ASM949871v1_genomic
GCF_011365345.1_ASM1136534v1_genomic
GCF_014217855.1_ASM1421785v1_genomic
GCF_014217875.1_ASM1421787v1_genomic
GCF_014224335.1_ASM1422433v1_genomic
GCF_014224355.1_ASM1422435v1_genomic
GCF_014279535.1_ASM1427953v1_genomic
GCF_014279555.1_ASM1427955v1_genomic
GCF_014279575.1_ASM1427957v1_genomic
GCF_014279595.1_ASM1427959v1_genomic
GCF_014279615.1_ASM1427961v1_genomic
GCF_014279635.1_ASM1427963v1_genomic
GCF_014279655.1_ASM1427965v1_genomic
GCF_014279755.1_ASM1427975v1_genomic
GCF_014279775.1_ASM1427977v1_genomic
GCF_014279795.1_ASM1427979v1_genomic
GCF_014279815.1_ASM1427981v1_genomic
GCF_014279835.1_ASM1427983v1_genomic
GCF_014279855.1_ASM1427985v1_genomic
GCF_014279875.1_ASM1427987v1_genomic
GCF_014279895.1_ASM1427989v1_genomic
GCF_014279915.1_ASM1427991v1_genomic
GCF_014279955.1_ASM1427995v1_genomic
GCF_014279975.1_ASM1427997v1_genomic
GCF_014279995.1_ASM1427999v1_genomic
GCF_014280015.1_ASM1428001v1_genomic
GCF_014280035.1_ASM1428003v1_genomic
GCF_014280055.1_ASM1428005v1_genomic
GCF_014280075.1_ASM1428007v1_genomic
GCF_014280095.1_ASM1428009v1_genomic
GCF_014280115.1_ASM1428011v1_genomic
GCF_014280175.1_ASM1428017v1_genomic
GCF_014280195.1_ASM1428019v1_genomic
GCF_014280215.1_ASM1428021v1_genomic
GCF_014304795.1_ASM1430479v1_genomic
GCF_014304815.1_ASM1430481v1_genomic
GCF_014698415.1_ASM1469841v1_genomic
GCF_014698895.1_ASM1469889v1_genomic
GCF_014698945.1_ASM1469894v1_genomic
GCF_015840335.1_ASM1584033v1_genomic
GCF_015840525.1_ASM1584052v1_genomic
GCF_015840715.1_ASM1584071v1_genomic
GCF_015840915.1_ASM1584091v1_genomic

**GCF_015841355.1_ASM1584135v1_genomic**
**GCF_900177365.1_IMG-taxon_2708742537_annotated_assembly_genomic**
**GCF_900177825.1_IMG-taxon_2708742468_annotated_assembly_genomic**
**GCF_900473895.1_N32_genomic**
**GCF_900473925.1_N5_genomic**
**GCF_900473935.1_UW105_genomic**
**GCF_900473955.1_GEYO_genomic**
**GCF_900473965.1_UW179A_genomic**
**GCF_900473975.1_N26_genomic**
**GCF_900474015.1_UW106_genomic**
**GCF_900474045.1_N19_genomic**
**GCF_900474085.1_UW86_genomic**
**GCF_900474185.1_UW69_genomic**
**GCF_900474245.1_UW179B_genomic**
**GCF_900474295.1_UW140_genomic**
**GCF000153285_1.scaf**
**GCA_000478825.2_ASM47882v2_genomic.fna**
**GCF_014695385.1_ASM1469538v1_genomic**
**GCF_014695505.1_ASM1469550v1_genomic**
**GCF_014695535.1_ASM1469553v1_genomic**
**GCF_014695675.1_ASM1469567v1_genomic**
**GCF_014695815.1_ASM1469581v1_genomic**
**GCF_014695845.1_ASM1469584v1_genomic**
**GCF_014695915.1_ASM1469591v1_genomic**
**GCF_014696035.1_ASM1469603v1_genomic**
**GCA_026240635.1_ASM2624063v1_genomic.fna**
**GCF_002252705.1_ASM225270v1_genomic**
**GCA_025370055.1_ASM2537005v1_genomic.fna**
**GCA_002754935.1_ASM275493v1_genomic.fna**
**GCA_003990665.2_ASM399066v2_genomic.fna**

| Order | Genus | Species |
|---|---|---|
| Chroococcales | Aphanothece | minutissima |
| Chroococcales | Aphanothece | hegewaldii |
| Chroococcales | Aphanothece | sacrum |
| Chroococcales | Aphanothece | sacrum |
| Chroococcales | Chlorogloea | Unknown |
| Chroococcales | Chondrocystis | Unknown |
| Chroococcales | Chroococcus | Unknown |
| Chroococcales | Chroogloeocystis | siderophila |
| Chroococcales | Crocosphaera | watsonii |
| Chroococcales | Crocosphaera | chwakensis |
| Chroococcales | Crocosphaera | subtropica |
| Chroococcales | Crocosphaera | subtropica |
| Chroococcales | Cyanobacterium | Unknown |
| Chroococcales | Cyanobacterium | Unknown |
| Chroococcales | Cyanobacterium | aponinum |
| Chroococcales | Cyanobacterium | Unknown |
| Chroococcales | Cyanobacterium | aponinum |
| Chroococcales | Cyanobacterium | aponinum |
| Chroococcales | Cyanobacterium | aponinum |
| Chroococcales | Cyanobacterium | stanieri |
| Chroococcales | Cyanobacterium | stanieri |
| Chroococcales | Euhalothece | natronophila |
| Chroococcales | Geminocystis | herdmanii |
| Chroococcales | Geminocystis | Unknown |
| Chroococcales | Geminocystis | Unknown |
| Chroococcales | Gloeocapsa | Unknown |
| Chroococcales | Gloeocapsa | Unknown |
| Chroococcales | Gloeocapsopsis | dulcis |
| Chroococcales | Gloeocapsopsis | crepidinum |
| Chroococcales | Gloeothece | citriformis |
| Chroococcales | Gloeothece | verrucosa |
| Chroococcales | Halothece | Unknown |
| Chroococcales | Microcystis | aeruginosa |
| Chroococcales | Microcystis | aeruginosa |
| Chroococcales | Microcystis | aeruginosa |
| Chroococcales | Microcystis | aeruginosa |
| Chroococcales | Microcystis | aeruginosa |
| Chroococcales | Microcystis | Unknown |
| Chroococcales | Microcystis | aeruginosa |
| Chroococcales | Microcystis | aeruginosa |
| Chroococcales | Microcystis | aeruginosa |
| Chroococcales | Microcystis | aeruginosa |
| Chroococcales | Microcystis | aeruginosa |
| Chroococcales | Microcystis | aeruginosa |
| Chroococcales | Microcystis | aeruginosa |
| Chroococcales | Microcystis | aeruginosa |
| Chroococcales | Microcystis | aeruginosa |

| | | |
|---|---|---|
| Chroococcales | Microcystis | aeruginosa |
| Chroococcales | Microcystis | aeruginosa |
| Chroococcales | Microcystis | aeruginosa |
| Chroococcales | Microcystis | aeruginosa |
| Chroococcales | Microcystis | aeruginosa |
| Chroococcales | Microcystis | aeruginosa |
| Chroococcales | Microcystis | aeruginosa |
| Chroococcales | Microcystis | aeruginosa |
| Chroococcales | Microcystis | aeruginosa |
| Chroococcales | Microcystis | aeruginosa |
| Chroococcales | Microcystis | aeruginosa |
| Chroococcales | Microcystis | aeruginosa |
| Chroococcales | Microcystis | Unknown |
| Chroococcales | Microcystis | Unknown |
| Chroococcales | Microcystis | aeruginosa |
| Chroococcales | Microcystis | aeruginosa |
| Chroococcales | Microcystis | aeruginosa |
| Chroococcales | Microcystis | viridis |
| Chroococcales | Microcystis | aeruginosa |
| Chroococcales | Microcystis | aeruginosa |
| Chroococcales | Microcystis | aeruginosa |
| Chroococcales | Microcystis | aeruginosa |
| Chroococcales | Microcystis | aeruginosa |
| Chroococcales | Microcystis | aeruginosa |
| Chroococcales | Microcystis | aeruginosa |
| Chroococcales | Microcystis | aeruginosa |
| Chroococcales | Microcystis | aeruginosa |
| Chroococcales | Microcystis | aeruginosa |
| Chroococcales | Microcystis | aeruginosa |
| Chroococcales | Microcystis | aeruginosa |
| Chroococcales | Microcystis | aeruginosa |
| Chroococcales | Microcystis | aeruginosa |
| Chroococcales | Microcystis | aeruginosa |
| Chroococcales | Microcystis | aeruginosa |
| Chroococcales | Microcystis | aeruginosa |
| Chroococcales | Microcystis | wesenbergii |
| Chroococcales | Microcystis | viridis |
| Chroococcales | Microcystis | flos-aquae |
| Chroococcales | Microcystis | elabens |
| Chroococcales | Microcystis | aeruginosa |
| Chroococcales | Microcystis | aeruginosa |
| Chroococcales | Microcystis | aeruginosa |
| Chroococcales | Microcystis | Unknown |
| Chroococcales | Microcystis | aeruginosa |
| Chroococcales | Microcystis | aeruginosa |
| Chroococcales | Microcystis | aeruginosa |
| Chroococcales | Microcystis | aeruginosa |
| Chroococcales | Microcystis | aeruginosa |

| | | |
|---|---|---|
| Chroococcales | Microcystis | aeruginosa |
| Chroococcales | Microcystis | aeruginosa |
| Chroococcales | Microcystis | aeruginosa |
| Chroococcales | Microcystis | aeruginosa |
| Chroococcales | Microcystis | aeruginosa |
| Chroococcales | Microcystis | aeruginosa |
| Chroococcales | Microcystis | aeruginosa |
| Chroococcales | Microcystis | aeruginosa |
| Chroococcales | Microcystis | Unknown |
| Chroococcales | Microcystis | panniformis |
| Chroococcales | Rippkaea | orientalis |
| Chroococcales | Rippkaea | orientalis |
| Chroococcales | Rubidibacter | lacunae |
| Chroococcidiopsidales | Aliterella | atlantica |
| Chroococcidiopsidales | Chroococcidiopsis | Unknown |
| Chroococcidiopsidales | Chroococcidiopsis | thermalis |
| Chroococcidiopsidales | Chroococcidiopsis | Unknown |
| Chroococcidiopsidales | Chroococcidiopsis | Unknown |
| Chroococcidiopsidales | Chroococcidiopsis | cubana |
| Chroococcidiopsidales | Chroococcidiopsis | Unknown |
| Chroococcidiopsidales | Chroococcidiopsis | Unknown |
| Gloeobacterales | Anthocerotibacter | panamensis |
| Gloeobacterales | Gloeobacter | violaceus |
| Gloeobacterales | Gloeobacter | kilaueensis |
| Gloeobacterales | Gloeobacter | morelensis |
| Gloeomargaritales | Gloeomargarita | lithophora |
| Gloeomargaritales | Unknown | Unknown |
| Nostocales | Aetokthonos | hydrillicola |
| Nostocales | Amazonocrinis | nigriterrae |
| Nostocales | Anabaena | Unknown |
| Nostocales | Anabaena | Unknown |
| Nostocales | Anabaena | cylindrica |
| Nostocales | Anabaena | Unknown |
| Nostocales | Anabaena | Unknown |
| Nostocales | Anabaena | Unknown |
| Nostocales | Anabaena | Unknown |
| Nostocales | Anabaena | circularis |
| Nostocales | Anabaena | variabilis |
| Nostocales | Anabaena | Unknown |
| Nostocales | Anabaena | Unknown |
| Nostocales | Anabaena | Unknown |
| Nostocales | Anabaena | Unknown |
| Nostocales | Anabaena | Unknown |
| Nostocales | Anabaena | Unknown |
| Nostocales | Anabaena | cylindrica |
| Nostocales | Anabaena | Unknown |
| Nostocales | Anabaena | Unknown |
| Nostocales | Anabaena | cylindrica |

| | | |
|---|---|---|
| Nostocales | Anabaena | sphaerica |
| Nostocales | Anabaena | subtropica |
| Nostocales | Anabaena | minutissima |
| Nostocales | Anabaena | variabilis |
| Nostocales | Anabaena | cylindrica |
| Nostocales | Anabaena | Unknown |
| Nostocales | Anabaena | azotica |
| Nostocales | Anabaena | lutea |
| Nostocales | Anabaena | variabilis |
| Nostocales | Anabaena | catenula |
| Nostocales | Anabaena | aphanizomenioides |
| Nostocales | Anabaena | elenkinii |
| Nostocales | Anabaenopsis | arnoldii |
| Nostocales | Aphanizomenon | Unknown |
| Nostocales | Aphanizomenon | flos-aquae |
| Nostocales | Aphanizomenon | flos-aquae |
| Nostocales | Aphanizomenon | flos-aquae |
| Nostocales | Aphanizomenon | flos-aquae |
| Nostocales | Aphanizomenon | Unknown |
| Nostocales | Aphanizomenon | flos-aquae |
| Nostocales | Aphanizomenon | flos-aquae |
| Nostocales | Aphanizomenon | Unknown |
| Nostocales | Aphanizomenon | Unknown |
| Nostocales | Aphanizomenon | flos-aquae |
| Nostocales | Aphanizomenon | flos-aquae |
| Nostocales | Aphanizomenon | flos-aquae |
| Nostocales | Aphanizomenon | flos-aquae |
| Nostocales | Aphanizomenon | flos-aquae |
| Nostocales | Atlanticothrix | silvestris |
| Nostocales | Aulosira | laxa |
| Nostocales | Aulosira | Unknown |
| Nostocales | Aulosira | Unknown |
| Nostocales | Brasilonema | sennae |
| Nostocales | Brasilonema | Unknown |
| Nostocales | Brasilonema | octagenarum |
| Nostocales | Brasilonema | bromeliae |
| Nostocales | Brasilonema | Unknown |
| Nostocales | Calothrix | Unknown |
| Nostocales | Calothrix | Unknown |
| Nostocales | Calothrix | Unknown |
| Nostocales | Calothrix | Unknown |
| Nostocales | Calothrix | Unknown |
| Nostocales | Calothrix | Unknown |
| Nostocales | Calothrix | Unknown |
| Nostocales | Calothrix | Unknown |
| Nostocales | Calothrix | brevissima |
| Nostocales | Calothrix | parasitica |
| Nostocales | Calothrix | Unknown |

| | | |
|---|---|---|
| Nostocales | Calothrix | Unknown |
| Nostocales | Calothrix | Unknown |
| Nostocales | Calothrix | Unknown |
| Nostocales | Calothrix | Unknown |
| Nostocales | Calothrix | Unknown |
| Nostocales | Calothrix | desertica |
| Nostocales | Calothrix | membranacea |
| Nostocales | Calothrix | anomala |
| Nostocales | Calothrix | Unknown |
| Nostocales | Calothrix | parietina |
| Nostocales | Calothrix | Unknown |
| Nostocales | Calothrix | Unknown |
| Nostocales | Calothrix | rhizosoleniae |
| Nostocales | Chlorogloeopsis | fritschii |
| Nostocales | Cuspidothrix | issatschenkoi |
| Nostocales | Cylindrospermopsis | raciborskii |
| Nostocales | Cylindrospermopsis | raciborskii |
| Nostocales | Cylindrospermopsis | Unknown |
| Nostocales | Cylindrospermopsis | raciborskii |
| Nostocales | Cylindrospermopsis | raciborskii |
| Nostocales | Cylindrospermopsis | raciborskii |
| Nostocales | Cylindrospermopsis | raciborskii |
| Nostocales | Cylindrospermopsis | raciborskii |
| Nostocales | Cylindrospermopsis | raciborskii |
| Nostocales | Cylindrospermopsis | raciborskii |
| Nostocales | Cylindrospermopsis | raciborskii |
| Nostocales | Cylindrospermopsis | raciborskii |
| Nostocales | Cylindrospermopsis | raciborskii |
| Nostocales | Cylindrospermopsis | raciborskii |
| Nostocales | Cylindrospermopsis | raciborskii |
| Nostocales | Cylindrospermopsis | raciborskii |
| Nostocales | Cylindrospermopsis | raciborskii |
| Nostocales | Cylindrospermopsis | raciborskii |
| Nostocales | Cylindrospermopsis | raciborskii |
| Nostocales | Cylindrospermopsis | raciborskii |
| Nostocales | Cylindrospermopsis | raciborskii |
| Nostocales | Cylindrospermopsis | raciborskii |
| Nostocales | Cylindrospermopsis | curvispora |
| Nostocales | Cylindrospermopsis | raciborskii |
| Nostocales | Cylindrospermum | Unknown |
| Nostocales | Cylindrospermum | stagnale |
| Nostocales | Cylindrospermum | Unknown |
| Nostocales | Dendronalium | phyllosphericum |
| Nostocales | Dolichospermum | circinale |
| Nostocales | Dolichospermum | circinale |
| Nostocales | Dolichospermum | circinale |
| Nostocales | Dolichospermum | circinale |
| Nostocales | Dolichospermum | circinale |

| | | |
|---|---|---|
| Nostocales | Dolichospermum | circinale |
| Nostocales | Dolichospermum | circinale |
| Nostocales | Dolichospermum | circinale |
| Nostocales | Dolichospermum | circinale |
| Nostocales | Dolichospermum | circinale |
| Nostocales | Dolichospermum | circinale |
| Nostocales | Dolichospermum | lemmermannii |
| Nostocales | Dolichospermum | planctonicum |
| Nostocales | Dolichospermum | Unknown |
| Nostocales | Dolichospermum | Unknown |
| Nostocales | Dolichospermum | Unknown |
| Nostocales | Dolichospermum | Unknown |
| Nostocales | Dolichospermum | heterosporum |
| Nostocales | Dolichospermum | circinale |
| Nostocales | Dolichospermum | circinale |
| Nostocales | Dolichospermum | compactum |
| Nostocales | Dolichospermum | flos-aquae |
| Nostocales | Dolichospermum | flos-aquae |
| Nostocales | Dolichospermum | flos-aquae |
| Nostocales | Dolichospermum | planctonicum |
| Nostocales | Dolichospermum | Unknown |
| Nostocales | Dolichospermum | Unknown |
| Nostocales | Dolichospermum | Unknown |
| Nostocales | Dolichospermum | Unknown |
| Nostocales | Dolichospermum | Unknown |
| Nostocales | Dolichospermum | Unknown |
| Nostocales | Dolichospermum | planctonicum |
| Nostocales | Dolichospermum | Unknown |
| Nostocales | Dolichospermum | Unknown |
| Nostocales | Fischerella | Unknown |
| Nostocales | Fischerella | thermalis |
| Nostocales | Fischerella | Unknown |
| Nostocales | Fischerella | muscicola |
| Nostocales | Fischerella | thermalis |
| Nostocales | Fischerella | muscicola |
| Nostocales | Fischerella | Unknown |
| Nostocales | Fischerella | Unknown |
| Nostocales | Fischerella | Unknown |
| Nostocales | Fischerella | major |
| Nostocales | Fischerella | Unknown |
| Nostocales | Fischerella | thermalis |
| Nostocales | Fischerella | thermalis |
| Nostocales | Fischerella | thermalis |
| Nostocales | Fischerella | thermalis |
| Nostocales | Fischerella | thermalis |
| Nostocales | Fischerella | thermalis |
| Nostocales | Fischerella | thermalis |
| Nostocales | Fischerella | thermalis |

| | | |
|---|---|---|
| Nostocales | Fischerella | thermalis |
| Nostocales | Fischerella | thermalis |
| Nostocales | Fischerella | thermalis |
| Nostocales | Fischerella | thermalis |
| Nostocales | Fischerella | thermalis |
| Nostocales | Fischerella | thermalis |
| Nostocales | Fischerella | thermalis |
| Nostocales | Fischerella | thermalis |
| Nostocales | Fischerella | thermalis |
| Nostocales | Fischerella | thermalis |
| Nostocales | Fischerella | thermalis |
| Nostocales | Fischerella | thermalis |
| Nostocales | Fischerella | thermalis |
| Nostocales | Fischerella | thermalis |
| Nostocales | Fischerella | Unknown |
| Nostocales | Fortiea | contorta |
| Nostocales | Fortiea | Unknown |
| Nostocales | Fremyella | diplosiphon |
| Nostocales | Halotia | branconii |
| Nostocales | Hapalosiphon | Unknown |
| Nostocales | Hassallia | byssoidea |
| Nostocales | Iningainema | tapete |
| Nostocales | Komarekiella | delphini-convector |
| Nostocales | Mastigocladopsis | repens |
| Nostocales | Mastigocoleus | testarum |
| Nostocales | Nodularia | sphaerocarpa |
| Nostocales | Nodularia | spumigena |
| Nostocales | Nodularia | spumigena |
| Nostocales | Nodularia | Unknown |
| Nostocales | Nodularia | spumigena |
| Nostocales | Nodularia | Unknown |
| Nostocales | Nodularia | sphaerocarpa |
| Nostocales | Nodularia | spumigena |
| Nostocales | Nodularia | spumigena |
| Nostocales | Nodularia | spumigena |
| Nostocales | Nodularia | spumigena |
| Nostocales | Nodularia | spumigena |
| Nostocales | Nodularia | spumigena |
| Nostocales | Nodularia | spumigena |
| Nostocales | Nodularia | spumigena |
| Nostocales | Nodularia | spumigena |
| Nostocales | Nodularia | spumigena |
| Nostocales | Nodularia | spumigena |
| Nostocales | Nodularia | spumigena |
| Nostocales | Nodularia | spumigena |
| Nostocales | Nodularia | spumigena |
| Nostocales | Nodularia | spumigena |
| Nostocales | Nodularia | spumigena |

| | | |
|---|---|---|
| Nostocales | Nostoc | commune |
| Nostocales | Nostoc | Unknown |
| Nostocales | Nostoc | Unknown |
| Nostocales | Nostoc | Unknown |
| Nostocales | Nostoc | Unknown |
| Nostocales | Nostoc | Unknown |
| Nostocales | Nostoc | Unknown |
| Nostocales | Nostoc | Unknown |
| Nostocales | Nostoc | punctiforme |
| Nostocales | Nostoc | azollae |
| Nostocales | Nostoc | Unknown |
| Nostocales | Nostoc | Unknown |
| Nostocales | Nostoc | piscinale |
| Nostocales | Nostoc | Unknown |
| Nostocales | Nostoc | Unknown |
| Nostocales | Nostoc | calcicola |
| Nostocales | Nostoc | Unknown |
| Nostocales | Nostoc | Unknown |
| Nostocales | Nostoc | Unknown |
| Nostocales | Nostoc | Unknown |
| Nostocales | Nostoc | Unknown |
| Nostocales | Nostoc | Unknown |
| Nostocales | Nostoc | linckia |
| Nostocales | Nostoc | carneum |
| Nostocales | Nostoc | Unknown |
| Nostocales | Nostoc | Unknown |
| Nostocales | Nostoc | linckia |
| Nostocales | Nostoc | linckia |
| Nostocales | Nostoc | linckia |
| Nostocales | Nostoc | linckia |
| Nostocales | Nostoc | linckia |
| Nostocales | Nostoc | linckia |
| Nostocales | Nostoc | linckia |
| Nostocales | Nostoc | linckia |
| Nostocales | Nostoc | linckia |
| Nostocales | Nostoc | linckia |
| Nostocales | Nostoc | flagelliforme |
| Nostocales | Nostoc | Unknown |
| Nostocales | Nostoc | cycadae |
| Nostocales | Nostoc | Unknown |
| Nostocales | Nostoc | Unknown |
| Nostocales | Nostoc | commune |
| Nostocales | Nostoc | Unknown |
| Nostocales | Nostoc | sphaeroides |
| Nostocales | Nostoc | Unknown |
| Nostocales | Nostoc | sphaeroides |
| Nostocales | Nostoc | Unknown |
| Nostocales | Nostoc | Unknown |

| | | |
|---|---|---|
| Nostocales | Nostoc | Unknown |
| Nostocales | Nostoc | Unknown |
| Nostocales | Nostoc | edaphicum |
| Nostocales | Nostoc | linckia |
| Nostocales | Nostoc | parmelioides |
| Nostocales | Nostoc | Unknown |
| Nostocales | Nostoc | Unknown |
| Nostocales | Nostoc | calcicola |
| Nostocales | Nostoc | Unknown |
| Nostocales | Nostoc | Unknown |
| Nostocales | Nostoc | Unknown |
| Nostocales | Nostoc | Unknown |
| Nostocales | Nostoc | Unknown |
| Nostocales | Nostoc | muscorum |
| Nostocales | Nostoc | Unknown |
| Nostocales | Nostoc | Unknown |
| Nostocales | Nostoc | spongiaeforme |
| Nostocales | Nostoc | punctiforme |
| Nostocales | Nostoc | foliaceum |
| Nostocales | Nostoc | linckia |
| Nostocales | Nostoc | Unknown |
| Nostocales | Nostoc | Unknown |
| Nostocales | Nostoc | paludosum |
| Nostocales | Nostoc | Unknown |
| Nostocales | Nostoc | Unknown |
| Nostocales | Nostoc | Unknown |
| Nostocales | Nostoc | Unknown |
| Nostocales | Nostoc | Unknown |
| Nostocales | Nostoc | Unknown |
| Nostocales | Nostoc | Unknown |
| Nostocales | Raphidiopsis | brookii |
| Nostocales | Richelia | intracellularis |
| Nostocales | Richelia | sinica |
| Nostocales | Rivularia | Unknown |
| Nostocales | Roholtiella | Unknown |
| Nostocales | Scytonema | hofmannii |
| Nostocales | Scytonema | millei |
| Nostocales | Scytonema | tolypothrichoides |
| Nostocales | Scytonema | hofmannii |
| Nostocales | Sphaerospermopsis | aphanizomenoides |
| Nostocales | Sphaerospermopsis | kisseleviana |
| Nostocales | Sphaerospermopsis | Unknown |
| Nostocales | Sphaerospermopsis | Unknown |
| Nostocales | Sphaerospermopsis | Unknown |
| Nostocales | Sphaerospermopsis | Unknown |
| Nostocales | Sphaerospermopsis | kisseleviana |
| Nostocales | Tolypothrix | Unknown |
| Nostocales | Tolypothrix | bouteillei |

| | | |
|---|---|---|
| Nostocales | Tolypothrix | campylonemoides |
| Nostocales | Tolypothrix | Unknown |
| Nostocales | Tolypothrix | Unknown |
| Nostocales | Tolypothrix | tenuis |
| Nostocales | Tolypothrix | Unknown |
| Nostocales | Tolypothrix | Unknown |
| Nostocales | Trichormus | variabilis |
| Nostocales | Trichormus | variabilis |
| Nostocales | Trichormus | variabilis |
| Nostocales | Trichormus | variabilis |
| Nostocales | Trichormus | variabilis |
| Nostocales | Trichormus | variabilis |
| Nostocales | Trichormus | variabilis |
| Nostocales | Trichormus | variabilis |
| Nostocales | Trichormus | variabilis |
| Nostocales | Unknown | Unknown |
| Nostocales | Westiellopsis | prolifica |
| Oscillatoriales | Allocoleopsis | franciscana |
| Oscillatoriales | Arthrospira | platensis |
| Oscillatoriales | Arthrospira | Unknown |
| Oscillatoriales | Arthrospira | platensis |
| Oscillatoriales | Arthrospira | platensis |
| Oscillatoriales | Arthrospira | platensis |
| Oscillatoriales | Arthrospira | platensis |
| Oscillatoriales | Arthrospira | platensis |
| Oscillatoriales | Arthrospira | platensis |
| Oscillatoriales | Arthrospira | platensis |
| Oscillatoriales | Arthrospira | platensis |
| Oscillatoriales | Arthrospira | Unknown |
| Oscillatoriales | Baaleninema | simplex |
| Oscillatoriales | Coleofasciculus | chthonoplastes |
| Oscillatoriales | Coleofasciculus | Unknown |
| Oscillatoriales | Coleofasciculus | Unknown |
| Oscillatoriales | Coleofasciculus | Unknown |
| Oscillatoriales | Coleofasciculus | Unknown |
| Oscillatoriales | Coleofasciculus | Unknown |
| Oscillatoriales | Coleofasciculus | Unknown |
| Oscillatoriales | Coleofasciculus | Unknown |
| Oscillatoriales | Coleofasciculus | Unknown |
| Oscillatoriales | Coleofasciculus | Unknown |
| Oscillatoriales | Coleofasciculus | Unknown |
| Oscillatoriales | Coleofasciculus | Unknown |
| Oscillatoriales | Crinalium | epipsammum |
| Oscillatoriales | Cyanothece | Unknown |
| Oscillatoriales | Cyanothece | Unknown |
| Oscillatoriales | Desertifilum | Unknown |
| Oscillatoriales | Desertifilum | Unknown |
| Oscillatoriales | Desertifilum | Unknown |

| | | |
|---|---|---|
| Oscillatoriales | Desertifilum | Unknown |
| Oscillatoriales | Geitlerinema | Unknown |
| Oscillatoriales | Geitlerinema | Unknown |
| Oscillatoriales | Kamptonema | Unknown |
| Oscillatoriales | Kamptonema | formosum |
| Oscillatoriales | Koinonema | Unknown |
| Oscillatoriales | Limnoraphis | robusta |
| Oscillatoriales | Limnospira | fusiformis |
| Oscillatoriales | Limnospira | maxima |
| Oscillatoriales | Limnospira | indica |
| Oscillatoriales | Limnospira | fusiformis |
| Oscillatoriales | Lyngbya | Unknown |
| Oscillatoriales | Lyngbya | aestuarii |
| Oscillatoriales | Lyngbya | confervoides |
| Oscillatoriales | Microcoleus | vaginatus |
| Oscillatoriales | Microcoleus | vaginatus |
| Oscillatoriales | Microcoleus | vaginatus |
| Oscillatoriales | Microcoleus | Unknown |
| Oscillatoriales | Microcoleus | Unknown |
| Oscillatoriales | Microcoleus | asticus |
| Oscillatoriales | Microcoleus | Unknown |
| Oscillatoriales | Microcoleus | Unknown |
| Oscillatoriales | Microcoleus | Unknown |
| Oscillatoriales | Microcoleus | Unknown |
| Oscillatoriales | Microcoleus | Unknown |
| Oscillatoriales | Microcoleus | Unknown |
| Oscillatoriales | Microcoleus | Unknown |
| Oscillatoriales | Microcoleus | Unknown |
| Oscillatoriales | Microcoleus | Unknown |
| Oscillatoriales | Microcoleus | Unknown |
| Oscillatoriales | Microcoleus | Unknown |
| Oscillatoriales | Microcoleus | Unknown |
| Oscillatoriales | Moorea | producens |
| Oscillatoriales | Moorea | producens |
| Oscillatoriales | Moorea | producens |
| Oscillatoriales | Moorea | bouillonii |
| Oscillatoriales | Moorena | producens |
| Oscillatoriales | Moorena | producens |
| Oscillatoriales | Okeania | hirsuta |
| Oscillatoriales | Okeania | Unknown |
| Oscillatoriales | Oscillatoria | acuminata |
| Oscillatoriales | Oscillatoria | nigro-viridis |
| Oscillatoriales | Oscillatoria | Unknown |
| Oscillatoriales | Oscillatoria | Unknown |
| Oscillatoriales | Oscillatoria | Unknown |
| Oscillatoriales | Oscillatoria | Unknown |
| Oscillatoriales | Oxynema | aestuarii |
| Oscillatoriales | Oxynema | Unknown |

| | | |
|---|---|---|
| Oscillatoriales | Phormidium | Unknown |
| Oscillatoriales | Phormidium | yuhuli |
| Oscillatoriales | Phormidium | willei |
| Oscillatoriales | Phormidium | ambiguum |
| Oscillatoriales | Phormidium | tenue |
| Oscillatoriales | Phormidium | Unknown |
| Oscillatoriales | Phormidium | Unknown |
| Oscillatoriales | Phormidium | Unknown |
| Oscillatoriales | Phormidium | tenue |
| Oscillatoriales | Phormidium | Unknown |
| Oscillatoriales | Phormidium | tenue |
| Oscillatoriales | Phormidium | Unknown |
| Oscillatoriales | Phormidium | Unknown |
| Oscillatoriales | Planktothricoides | Unknown |
| Oscillatoriales | Planktothricoides | Unknown |
| Oscillatoriales | Planktothricoides | Unknown |
| Oscillatoriales | Planktothrix | agardhii |
| Oscillatoriales | Planktothrix | agardhii |
| Oscillatoriales | Planktothrix | agardhii |
| Oscillatoriales | Planktothrix | agardhii |
| Oscillatoriales | Planktothrix | agardhii |
| Oscillatoriales | Planktothrix | pseudagardhii |
| Oscillatoriales | Planktothrix | agardhii |
| Oscillatoriales | Planktothrix | agardhii |
| Oscillatoriales | Planktothrix | mougeotii |
| Oscillatoriales | Planktothrix | prolifica |
| Oscillatoriales | Planktothrix | rubescens |
| Oscillatoriales | Planktothrix | prolifica |
| Oscillatoriales | Planktothrix | agardhii |
| Oscillatoriales | Planktothrix | prolifica |
| Oscillatoriales | Planktothrix | agardhii |
| Oscillatoriales | Planktothrix | agardhii |
| Oscillatoriales | Planktothrix | Unknown |
| Oscillatoriales | Planktothrix | Unknown |
| Oscillatoriales | Planktothrix | mougeotii |
| Oscillatoriales | Planktothrix | Unknown |
| Oscillatoriales | Planktothrix | tepida |
| Oscillatoriales | Planktothrix | paucivesiculata |
| Oscillatoriales | Planktothrix | rubescens |
| Oscillatoriales | Planktothrix | serta |
| Oscillatoriales | Planktothrix | agardhii |
| Oscillatoriales | Plectonema | radiosum |
| Oscillatoriales | Trichodesmium | erythraeum |
| Oscillatoriales | Tychonema | bourrellyi |
| Oscillatoriales | Tychonema | Unknown |
| Oscillatoriales | Tychonema | Unknown |
| Oscillatoriales | Tychonema | Unknown |
| Oscillatoriales | Tychonema | Unknown |

| | | |
|---|---|---|
| Pleurocapsales | Hydrococcus | rivularis |
| Pleurocapsales | Myxosarcina | Unknown |
| Pleurocapsales | Pleurocapsa | Unknown |
| Pleurocapsales | Pleurocapsa | Unknown |
| Pleurocapsales | Pleurocapsa | Unknown |
| Pleurocapsales | Stanieria | cyanosphaera |
| Pleurocapsales | Stanieria | Unknown |
| Pleurocapsales | Xenococcus | Unknown |
| Pseudanabaenales | Alkalinema | Unknown |
| Pseudanabaenales | Halomicronema | hongdechloris |
| Pseudanabaenales | Halomicronema | excentricum |
| Pseudanabaenales | Halomicronema | Unknown |
| Pseudanabaenales | Kovacikia | minuta |
| Pseudanabaenales | Leptolyngbya | boryana |
| Pseudanabaenales | Leptolyngbya | Unknown |
| Pseudanabaenales | Leptolyngbya | Unknown |
| Pseudanabaenales | Leptolyngbya | Unknown |
| Pseudanabaenales | Leptolyngbya | Unknown |
| Pseudanabaenales | Leptolyngbya | boryana |
| Pseudanabaenales | Leptolyngbya | Unknown |
| Pseudanabaenales | Leptolyngbya | Unknown |
| Pseudanabaenales | Leptolyngbya | Unknown |
| Pseudanabaenales | Leptolyngbya | Unknown |
| Pseudanabaenales | Leptolyngbya | Unknown |
| Pseudanabaenales | Leptolyngbya | valderiana |
| Pseudanabaenales | Leptolyngbya | valderiana |
| Pseudanabaenales | Leptolyngbya | Unknown |
| Pseudanabaenales | Leptolyngbya | boryana |
| Pseudanabaenales | Leptolyngbya | boryana |
| Pseudanabaenales | Leptolyngbya | ohadii |
| Pseudanabaenales | Leptolyngbya | Unknown |
| Pseudanabaenales | Leptolyngbya | boryana |
| Pseudanabaenales | Leptolyngbya | Unknown |
| Pseudanabaenales | Leptolyngbya | Unknown |
| Pseudanabaenales | Leptolyngbya | Unknown |
| Pseudanabaenales | Leptolyngbya | Unknown |
| Pseudanabaenales | Leptolyngbya | Unknown |
| Pseudanabaenales | Leptolyngbya | Unknown |
| Pseudanabaenales | Leptolyngbya | Unknown |
| Pseudanabaenales | Leptolyngbya | Unknown |
| Pseudanabaenales | Leptolyngbya | Unknown |
| Pseudanabaenales | Leptolyngbya | Unknown |
| Pseudanabaenales | Leptolyngbya | Unknown |
| Pseudanabaenales | Leptolyngbya | Unknown |
| Pseudanabaenales | Leptolyngbya | Unknown |
| Pseudanabaenales | Leptolyngbya | Unknown |
| Pseudanabaenales | Leptolyngbya | Unknown |
| Pseudanabaenales | Leptolyngbya | Unknown |

| | | |
|---|---|---|
| Pseudanabaenales | Leptothermofonsia | sichuanensis |
| Pseudanabaenales | Leptothermofonsia | sichuanensis |
| Pseudanabaenales | Leptothoe | kymatousa |
| Pseudanabaenales | Leptothoe | spongobia |
| Pseudanabaenales | Limnothrix | rosea |
| Pseudanabaenales | Limnothrix | Unknown |
| Pseudanabaenales | Limnothrix | Unknown |
| Pseudanabaenales | Limnothrix | Unknown |
| Pseudanabaenales | Limnothrix | Unknown |
| Pseudanabaenales | Limnothrix | Unknown |
| Pseudanabaenales | Limnothrix | Unknown |
| Pseudanabaenales | Lusitaniella | coriacea |
| Pseudanabaenales | Myxacorys | almedinensis |
| Pseudanabaenales | Neosynechococcus | sphagnicola |
| Pseudanabaenales | Nodosilinea | nodulosa |
| Pseudanabaenales | Nodosilinea | Unknown |
| Pseudanabaenales | Nodosilinea | Unknown |
| Pseudanabaenales | Nodosilinea | Unknown |
| Pseudanabaenales | Nodosilinea | Unknown |
| Pseudanabaenales | Nodosilinea | Unknown |
| Pseudanabaenales | Nodosilinea | Unknown |
| Pseudanabaenales | Oculatella | Unknown |
| Pseudanabaenales | Oculatella | Unknown |
| Pseudanabaenales | Phormidesmis | priestleyi |
| Pseudanabaenales | Phormidesmis | priestleyi |
| Pseudanabaenales | Prochlorothrix | hollandica |
| Pseudanabaenales | Pseudanabaena | Unknown |
| Pseudanabaenales | Pseudanabaena | galeata |
| Pseudanabaenales | Pseudanabaena | Unknown |
| Pseudanabaenales | Pseudanabaena | Unknown |
| Pseudanabaenales | Pseudanabaena | Unknown |
| Pseudanabaenales | Pseudanabaena | Unknown |
| Pseudanabaenales | Pseudanabaena | Unknown |
| Pseudanabaenales | Pseudanabaena | Unknown |
| Pseudanabaenales | Pseudanabaena | Unknown |
| Pseudanabaenales | Pseudanabaena | yagii |
| Pseudanabaenales | Pseudanabaena | biceps |
| Pseudanabaenales | Pseudanabaena | Unknown |
| Pseudanabaenales | Pseudanabaena | Unknown |
| Pseudanabaenales | Pseudanabaena | Unknown |
| Pseudanabaenales | Pseudanabaena | Unknown |
| Pseudanabaenales | Pseudanabaena | Unknown |
| Pseudanabaenales | Pseudanabaena | Unknown |
| Pseudanabaenales | Romeria | gracilis |
| Pseudanabaenales | Thermoleptolyngbya | sichuanensis |
| Pseudanabaenales | Thermosynechococcus | Unknown |
| Pseudanabaenales | Thermosynechococcus | Unknown |
| Pseudanabaenales | Thermosynechococcus | elongatus |

| | | |
|---|---|---|
| Pseudanabaenales | Thermosynechococcus | Unknown |
| Pseudanabaenales | Thermosynechococcus | elongatus |
| Pseudanabaenales | Thermosynechococcus | vulcanus |
| Pseudanabaenales | Thermosynechococcus | Unknown |
| Pseudanabaenales | Thermosynechococcus | Unknown |
| Spirulinales | Spirulina | subsalsa |
| Spirulinales | Spirulina | major |
| Spirulinales | Spirulina | major |
| Spirulinales | Spirulina | major |
| Spirulinales | Spirulina | Unknown |
| Spirulinales | Spirulina | subsalsa |
| Synechococcales | Acaryochloris | marina |
| Synechococcales | Acaryochloris | Unknown |
| Synechococcales | Acaryochloris | marina |
| Synechococcales | Acaryochloris | marina |
| Synechococcales | Acaryochloris | Unknown |
| Synechococcales | Acaryochloris | marina |
| Synechococcales | Aphanocapsa | montana |
| Synechococcales | Candidatus | Atelocyanobacterium thalassa |
| Synechococcales | Candidatus | Synechococcus spongiarum |
| Synechococcales | Candidatus | Synechococcus spongiarum |
| Synechococcales | Candidatus | spongiarum |
| Synechococcales | Candidatus Paraprochlo | terpiosi |
| Synechococcales | Chamaesiphon | minutus |
| Synechococcales | Cyanobium | Unknown |
| Synechococcales | Cyanobium | Unknown |
| Synechococcales | Cyanobium | gracile |
| Synechococcales | Cyanobium | usitatum |
| Synechococcales | Cyanobium | Unknown |
| Synechococcales | Cyanobium | Unknown |
| Synechococcales | Cyanobium | Unknown |
| Synechococcales | Cyanobium | Unknown |
| Synechococcales | Cyanobium | Unknown |
| Synechococcales | Cyanobium | Unknown |
| Synechococcales | Dactylococcopsis | salina |
| Synechococcales | Merismopedia | glauca |
| Synechococcales | Prochlorococcus | marinus |
| Synechococcales | Prochlorococcus | marinus |
| Synechococcales | Prochlorococcus | marinus |
| Synechococcales | Prochlorococcus | marinus |
| Synechococcales | Prochlorococcus | marinus |
| Synechococcales | Prochlorococcus | marinus |
| Synechococcales | Prochlorococcus | marinus |
| Synechococcales | Prochlorococcus | marinus |
| Synechococcales | Prochlorococcus | marinus |
| Synechococcales | Prochlorococcus | marinus |
| Synechococcales | Prochlorococcus | marinus |
| Synechococcales | Prochlorococcus | marinus |

| | | |
|---|---|---|
| Synechococcales | Prochlorococcus | marinus |
| Synechococcales | Prochlorococcus | marinus |
| Synechococcales | Prochlorococcus | Unknown |
| Synechococcales | Prochlorococcus | Unknown |
| Synechococcales | Prochlorococcus | marinus |
| Synechococcales | Prochlorococcus | marinus |
| Synechococcales | Prochlorococcus | marinus |
| Synechococcales | Prochlorococcus | marinus |
| Synechococcales | Prochlorococcus | marinus |
| Synechococcales | Prochlorococcus | marinus |
| Synechococcales | Prochlorococcus | marinus |
| Synechococcales | Prochlorococcus | marinus |
| Synechococcales | Prochlorococcus | marinus |
| Synechococcales | Prochlorococcus | marinus |
| Synechococcales | Prochlorococcus | marinus |
| Synechococcales | Prochlorococcus | marinus |
| Synechococcales | Prochlorococcus | marinus |
| Synechococcales | Prochlorococcus | marinus |
| Synechococcales | Prochlorococcus | Unknown |
| Synechococcales | Prochlorococcus | Unknown |
| Synechococcales | Prochlorococcus | Unknown |
| Synechococcales | Prochlorococcus | marinus |
| Synechococcales | Prochlorococcus | marinus |
| Synechococcales | Prochlorococcus | marinus |
| Synechococcales | Prochlorococcus | Unknown |
| Synechococcales | Prochlorococcus | Unknown |
| Synechococcales | Prochlorococcus | Unknown |
| Synechococcales | Prochlorococcus | marinus |
| Synechococcales | Prochlorococcus | Unknown |
| Synechococcales | Prochlorococcus | marinus |
| Synechococcales | Prochlorococcus | marinus |
| Synechococcales | Prochlorococcus | marinus |
| Synechococcales | Prochlorococcus | marinus |
| Synechococcales | Prochlorococcus | marinus |
| Synechococcales | Prochlorococcus | Unknown |
| Synechococcales | Prochlorococcus | Unknown |
| Synechococcales | Prochlorococcus | marinus |
| Synechococcales | Prochlorococcus | marinus |
| Synechococcales | Prochlorococcus | Unknown |
| Synechococcales | Prochlorococcus | Unknown |
| Synechococcales | Prochlorococcus | Unknown |
| Synechococcales | Prochlorococcus | Unknown |
| Synechococcales | Prochlorococcus | Unknown |
| Synechococcales | Prochlorococcus | Unknown |
| Synechococcales | Prochlorococcus | Unknown |
| Synechococcales | Prochlorococcus | Unknown |
| Synechococcales | Prochlorococcus | Unknown |
| Synechococcales | Prochlorococcus | Unknown |

| | | |
|---|---|---|
| Synechococcales | Prochlorococcus | Unknown |
| Synechococcales | Prochlorococcus | Unknown |
| Synechococcales | Prochlorococcus | Unknown |
| Synechococcales | Prochlorococcus | Unknown |
| Synechococcales | Prochlorococcus | Unknown |
| Synechococcales | Prochlorococcus | Unknown |
| Synechococcales | Prochlorococcus | Unknown |
| Synechococcales | Prochlorococcus | Unknown |
| Synechococcales | Prochlorococcus | Unknown |
| Synechococcales | Prochlorococcus | Unknown |
| Synechococcales | Prochlorococcus | Unknown |
| Synechococcales | Prochlorococcus | Unknown |
| Synechococcales | Prochlorococcus | marinus |
| Synechococcales | Prochlorococcus | marinus |
| Synechococcales | Prochlorococcus | marinus |
| Synechococcales | Prochlorococcus | Unknown |
| Synechococcales | Prochlorococcus | Unknown |
| Synechococcales | Prochlorococcus | Unknown |
| Synechococcales | Prochlorococcus | Unknown |
| Synechococcales | Prochlorococcus | marinus |
| Synechococcales | Prochlorococcus | marinus |
| Synechococcales | Prochlorococcus | marinus |
| Synechococcales | Prochlorococcus | marinus |
| Synechococcales | Prochlorococcus | marinus |
| Synechococcales | Prochlorococcus | marinus |
| Synechococcales | Prochlorococcus | marinus |
| Synechococcales | Prochlorococcus | marinus |
| Synechococcales | Prochlorococcus | marinus |
| Synechococcales | Prochlorococcus | marinus |
| Synechococcales | Prochlorococcus | marinus |
| Synechococcales | Prochlorococcus | marinus |
| Synechococcales | Prochlorococcus | marinus |
| Synechococcales | Synechococcus | Unknown |
| Synechococcales | Synechococcus | Unknown |
| Synechococcales | Synechococcus | Unknown |
| Synechococcales | Synechococcus | Unknown |
| Synechococcales | Synechococcus | Unknown |
| Synechococcales | Synechococcus | elongatus |
| Synechococcales | Synechococcus | Unknown |
| Synechococcales | Synechococcus | Unknown |
| Synechococcales | Synechococcus | Unknown |
| Synechococcales | Synechococcus | Unknown |
| Synechococcales | Synechococcus | Unknown |
| Synechococcales | Synechococcus | elongatus |
| Synechococcales | Synechococcus | elongatus |
| Synechococcales | Synechococcus | Unknown |
| Synechococcales | Synechococcus | Unknown |
| Synechococcales | Synechococcus | Unknown |

| | | |
|---|---|---|
| Synechococcales | Synechococcus | Unknown |
| Synechococcales | Synechococcus | Unknown |
| Synechococcales | Synechococcus | Unknown |
| Synechococcales | Synechococcus | Unknown |
| Synechococcales | Synechococcus | Unknown |
| Synechococcales | Synechococcus | Unknown |
| Synechococcales | Synechococcus | Unknown |
| Synechococcales | Synechococcus | Unknown |
| Synechococcales | Synechococcus | Unknown |
| Synechococcales | Synechococcus | Unknown |
| Synechococcales | Synechococcus | Unknown |
| Synechococcales | Synechococcus | Unknown |
| Synechococcales | Synechococcus | Unknown |
| Synechococcales | Synechococcus | Unknown |
| Synechococcales | Synechococcus | Unknown |
| Synechococcales | Synechococcus | Unknown |
| Synechococcales | Synechococcus | Unknown |
| Synechococcales | Synechococcus | Unknown |
| Synechococcales | Synechococcus | Unknown |
| Synechococcales | Synechococcus | Unknown |
| Synechococcales | Synechococcus | Unknown |
| Synechococcales | Synechococcus | Unknown |
| Synechococcales | Synechococcus | Unknown |
| Synechococcales | Synechococcus | Unknown |
| Synechococcales | Synechococcus | Unknown |
| Synechococcales | Synechococcus | Unknown |
| Synechococcales | Synechococcus | Unknown |
| Synechococcales | Synechococcus | Unknown |
| Synechococcales | Synechococcus | Unknown |
| Synechococcales | Synechococcus | Unknown |
| Synechococcales | Synechococcus | Unknown |
| Synechococcales | Synechococcus | Unknown |
| Synechococcales | Synechococcus | Unknown |
| Synechococcales | Synechococcus | Unknown |
| Synechococcales | Synechococcus | Unknown |
| Synechococcales | Synechococcus | Unknown |
| Synechococcales | Synechococcus | Unknown |
| Synechococcales | Synechococcus | Unknown |
| Synechococcales | Synechococcus | lividus |
| Synechococcales | Synechococcus | Unknown |
| Synechococcales | Synechococcus | Unknown |
| Synechococcales | Synechococcus | Unknown |
| Synechococcales | Synechococcus | Unknown |
| Synechococcales | Synechococcus | Unknown |
| Synechococcales | Synechococcus | Unknown |
| Synechococcales | Synechococcus | lacustris |
| Synechococcales | Synechococcus | elongatus |
| Synechococcales | Synechococcus | elongatus |

| | | |
|---|---|---|
| Synechococcales | Synechococcus | Unknown |
| Synechococcales | Synechococcus | Unknown |
| Synechococcales | Synechococcus | Unknown |
| Synechococcales | Synechococcus | Unknown |
| Synechococcales | Synechococcus | Unknown |
| Synechococcales | Synechococcus | elongatus |
| Synechococcales | Synechococcus | Unknown |
| Synechococcales | Synechococcus | Unknown |
| Synechococcales | Synechococcus | Unknown |
| Synechococcales | Synechococcus | Unknown |
| Synechococcales | Synechococcus | Unknown |
| Synechococcales | Synechococcus | Unknown |
| Synechococcales | Synechococcus | Unknown |
| Synechococcales | Synechococcus | Unknown |
| Synechococcales | Synechococcus | Unknown |
| Synechococcales | Synechococcus | Unknown |
| Synechococcales | Synechococcus | Unknown |
| Synechococcales | Synechococcus | Unknown |
| Synechococcales | Synechococcus | Unknown |
| Synechococcales | Synechococcus | Unknown |
| Synechococcales | Synechococcus | Unknown |
| Synechococcales | Synechococcus | Unknown |
| Synechococcales | Synechococcus | Unknown |
| Synechococcales | Synechococcus | Unknown |
| Synechococcales | Synechococcus | Unknown |
| Synechococcales | Synechococcus | Unknown |
| Synechococcales | Synechococcus | Unknown |
| Synechococcales | Synechococcus | Unknown |
| Synechococcales | Synechococcus | Unknown |
| Synechococcales | Synechococcus | Unknown |
| Synechococcales | Synechococcus | Unknown |
| Synechococcales | Synechococcus | Unknown |
| Synechococcales | Synechococcus | Unknown |
| Synechococcales | Synechococcus | Unknown |
| Synechococcales | Synechococcus | Unknown |
| Synechococcales | Synechococcus | Unknown |
| Synechococcales | Synechococcus | Unknown |
| Synechococcales | Synechococcus | Unknown |
| Synechococcales | Synechococcus | Unknown |
| Synechococcales | Synechococcus | Unknown |
| Synechococcales | Synechococcus | Unknown |
| Synechococcales | Synechococcus | elongatus |
| Synechococcales | Synechococcus | elongatus |
| Synechococcales | Synechococcus | Unknown |
| Synechococcales | Synechococcus | Unknown |
| Synechococcales | Synechococcus | Unknown |
| Synechococcales | Synechococcus | Unknown |
| Synechococcales | Synechococcus | Unknown |

| | | |
|---|---|---|
| Synechococcales | Synechococcus | Unknown |
| Synechococcales | Synechococcus | Unknown |
| Synechococcales | Synechococcus | Unknown |
| Synechococcales | Synechococcus | Unknown |
| Synechococcales | Synechococcus | Unknown |
| Synechococcales | Synechococcus | Unknown |
| Synechococcales | Synechococcus | Unknown |
| Synechococcales | Synechococcus | Unknown |
| Synechococcales | Synechococcus | Unknown |
| Synechococcales | Synechococcus | Unknown |
| Synechococcales | Synechococcus | Unknown |
| Synechococcales | Synechococcus | Unknown |
| Synechococcales | Synechococcus | Unknown |
| Synechococcales | Synechococcus | Unknown |
| Synechococcales | Synechococcus | Unknown |
| Synechococcales | Synechococcus | Unknown |
| Synechococcales | Synechocystis | Unknown |
| Synechococcales | Trichocoleus | Unknown |
| Synechococcales | Trichocoleus | Unknown |
| Synechococcales | Trichocoleus | Unknown |
| Synechococcales | Trichocoleus | Unknown |
| Synechococcales | Trichocoleus | Unknown |
| Synechococcales | Trichocoleus | Unknown |
| Synechococcales | Trichocoleus | Unknown |
| Synechococcales | Trichocoleus | Unknown |
| Synechococcales | Trichothermofontia | sichuanensis |
| Synechococcales | Vulcanococcus | limneticus |
| Synechococcales | Woronichinia | naegeliana |
| Thermostichales | Thermostichus | lividus |
| Thermostichales | Thermostichus | vulcanus |
